# Cell and Transcriptomic Diversity of Infrapatellar Fat Pad during Knee Osteoarthritis

**DOI:** 10.1101/2024.04.04.588106

**Authors:** Hayley Peters, Pratibha Potla, Jason S. Rockel, Teodora Tockovska, Chiara Pastrello, Igor Jurisica, Keemo Delos Santos, Shabana Vohra, Starlee Lively, Kim Perry, Nikita Looby, Sheng Han Li, Vinod Chandran, Katrina Hueniken, Paramvir Kaur, Anthony V. Perruccio, Nizar N. Mahomed, Y. Raja Rampersaud, Khalid A. Syed, Eric Gracey, Roman Krawetz, Matthew B. Buechler, Rajiv Gandhi, Mohit Kapoor

## Abstract

**Objectives:** In this study, we employ a multi-omic approach to identify major cell types and subsets, and their transcriptomic profiles within the infrapatellar fat pad (IFP), and to determine differences in the IFP based on knee osteoarthritis (KOA), sex, and obesity status.

**Methods:** Single-nucleus RNA sequencing of 82,924 nuclei from 21 IFPs (n=6 healthy control and n=15 KOA donors), spatial transcriptomics and bioinformatic analysis were used to identify contributions of the IFP to KOA. We mapped cell subclusters from other white adipose tissues using publicly available literature. The diversity of fibroblasts within the IFP was investigated by bioinformatic analyses, comparing by KOA, sex, and obesity status. Metabolomics was used to further explore differences in fibroblasts by obesity status.

**Results:** We identified multiple subclusters of fibroblasts, macrophages, adipocytes, and endothelial cells with unique transcriptomic profiles. Using spatial transcriptomics, we resolved distributions of cell types and their transcriptomic profiles, and computationally identified putative cell-cell communication networks. Furthermore, we identified transcriptomic differences in fibroblasts from KOA versus healthy control donor IFPs, female versus male KOA-IFPs, and obese versus normal body mass index (BMI) KOA-IFPs. Finally, using metabolomics, we defined differences in metabolite levels in supernatants of naïve, profibrotic- and proinflammatory stimuli-treated fibroblasts from obese compared to normal BMI KOA-IFP.

**Conclusions:** Overall, by employing a multi-omic approach, this study provides the first comprehensive map of cellular and transcriptomic diversity of human IFP and identifies IFP fibroblasts as a key cell type contributing to transcriptomic and metabolic differences related to KOA disease, sex, or obesity.

## Introduction

Osteoarthritis (OA) is a progressive disease that diminishes the quality of life of those afflicted. It is a leading cause of years lived with disability, and the number of cases globally has increased by 132% since 1990, placing growing burden on healthcare systems^1^. Among adults aged 45 years and older, approximately 30% have detectable knee (K)OA with half experiencing chronic painful symptoms, the majority being females^2^. In addition, individuals experience stiffness and muscle weakness causing reduced mobility^2^.

Primary (non-traumatic) KOA is associated with degradation of articular cartilage, chondrocyte loss and subchondral bone remodeling^3^. As KOA progresses, mobility is reduced, accompanied by severe pain. In addition, elevated joint cytokine levels increase angiogenesis and inflammation of the synovium and fat pads^3^. There is growing interest in the contribution of soft tissues to KOA pathogenesis; however, few studies have focused on understanding the infrapatellar fat pad (IFP).

The IFP is the largest fat pad within the knee, located posterior to the inferior pole of the patella, anterior to the tibia, and between the anterior horns of the meniscus and femur^4^. Composed of white fibrous adipose tissue, the main function of the IFP is to aid with shock absorption and support joint structure^4^. The IFP is highly vascularized, becoming inflamed during KOA, leading to fibrosis and structural changes, potentially contributing to significant pain^5,6^. A recent study identified some cell types in combined IFP and synovial tissue^7^. However, to our knowledge, no study has focused on understanding the complex diversity of the cellular composition (including cell subclusters) and their transcriptomic profiles exclusively in the IFP.

Obesity [body mass index (BMI) ≥ 30 kg/m^2^] is one of the strongest risk factors for developing KOA^2^. Obese individuals demonstrate chronic, low-grade, systemic inflammation and increased mechanical stress on knee joints, consequentially increasing pro-inflammatory adipokines and cytokines that promote KOA^8^. Females are also at increased risk of KOA, and typically present with more severe pain and functional limitations, as compared to males^9^. How obesity status and sex impact the IFP during KOA pathogenesis is not fully understood.

In this study, we used single-nucleus RNA sequencing (snRNA-seq), spatial transcriptomics and advanced bioinformatic methods to identify cell types and subclusters within IFPs from patient-matched samples, creating a comprehensive cell and transcriptomic map of the IFP. We determined that IFP contains multiple cell types with specific cell subclusters, each having a unique transcriptomic profile and spatial distribution. Cell-cell communication analysis revealed putative signalling patterns across all major cell types. We demonstrated differences in the transcriptomes of fibroblasts from IFPs of KOA vs healthy controls, females vs males with KOA, and obese BMI (30-40 kg/m^2^) vs normal BMI (18.5-24.9 kg/m^2^) with KOA, suggesting putative mechanisms by which fibroblasts may contribute to KOA pathogenesis. Finally, metabolomics revealed that metabolite levels in culture supernatants were modified in fibroblasts from obese compared to normal BMI KOA-IFPs and were further altered by profibrotic and proinflammatory stimuli. Overall, we provide a comprehensive map of the cellular and transcriptomic diversity of the human IFP, with considerations for KOA, sex, and obesity status.

## Methods

Full detailed materials and methods are available in supplementary information.

## Results

### Cellular composition of the IFP

IFPs from 21 study participants, including 6 healthy control and 15 with KOA donors, were analyzed using snRNA-seq and bioinformatics analyses to identify cell types (Fig. 1A; Supplementary Fig. 1; Supplementary Table 1). A total of 82,924 nuclei were sequenced and, after filtering, 73,808 nuclei were analyzed. Eight cell types were identified based on canonical cell markers: fibroblasts, adipocytes, macrophages, endothelial cells, dendritic cells, smooth muscle cells, lymphocytes, and mast cells (Fig. 1B-D; Supplementary Fig. 2A). Major cell populations contributing to the IFP (>10% of nuclei analyzed) were fibroblasts (44.35%), macrophages (19.44%), adipocytes (16.41%), and endothelial cells (12.12) (Fig. 1C,D).

**Figure 1.**
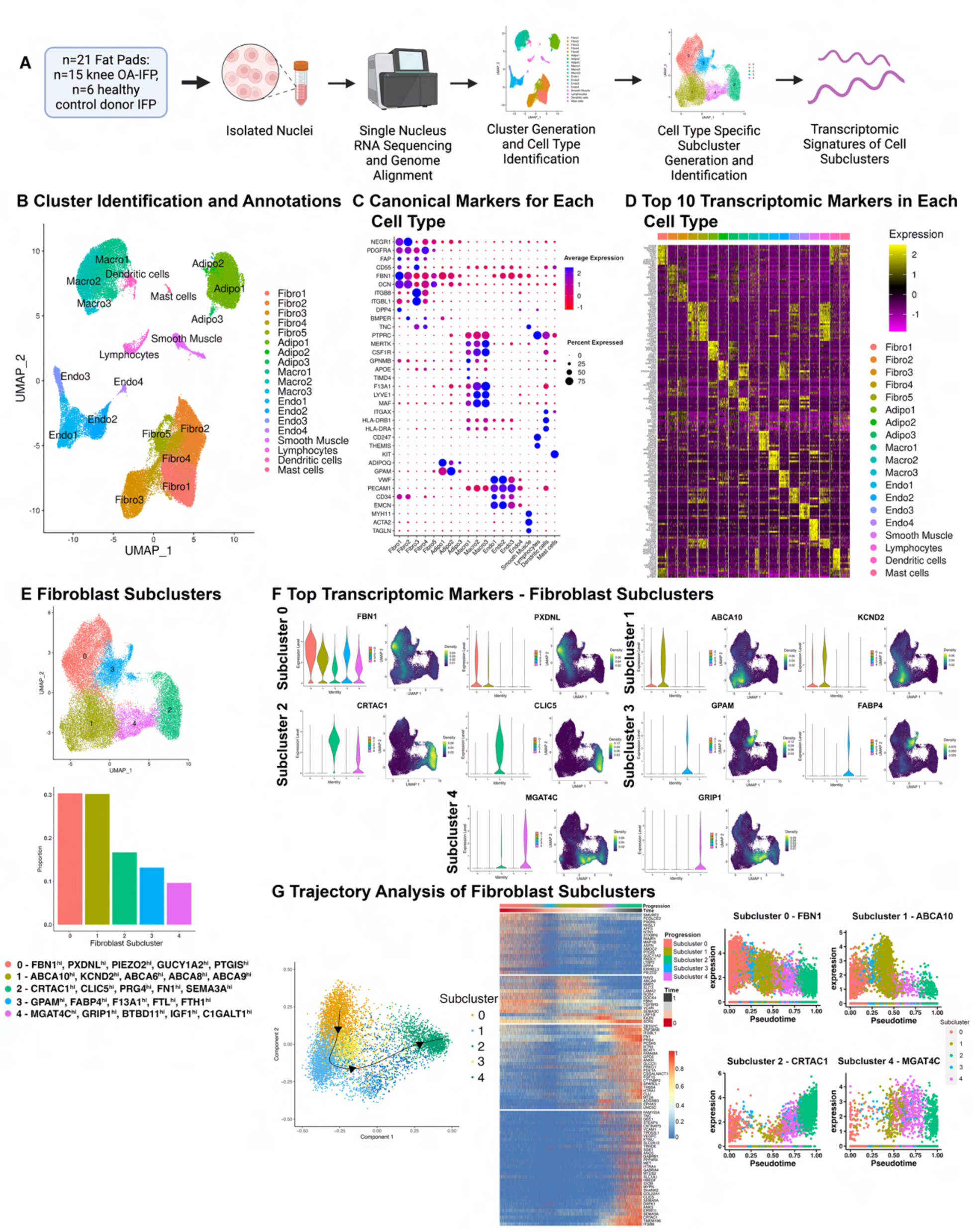
Infrapatellar Fat Pad (IFP) Cell Populations and Fibroblast Subclusters Identified by snRNA-seq (n=21). A) Schematic workflow to identify cell types and subclusters within human IFPs (Created with BioRender.com). B) Uniform manifold approximation and projection (UMAP) of clusters identified in IFPs. C) Dot plot with the average and percent population expression of canonical markers for each cell population. D) Heat map displaying expression of the top ten differentially-expressed genes (DEGs) within each cell population. E) UMAP of fibroblast subclusters and bar plot depicting fibroblast proportion of each subcluster. Top 5 unique DEGs defining each subcluster are indicated. F) Violin and nebulosa plots displaying the expression of the top two unique differentially expressed genes within each fibroblast subcluster. Nebulosa plots display expression density ranging from dark purple (low) to yellow (high). G) 2D plot showing inferred trajectory across pseudotime of integrated fibroblast subclusters using SCORPIUS and heatmap demonstrating the change in average scaled expression of the top 94 important genes defining fibroblast subcluster trajectory across pseudotime (q<0.05). Genes in the heatmap are clustered into four modules, each representing a set of genes expressed at different timepoints of the trajectory. Jitter plots depict the expression of the top marker gene for distinct subclusters across pseudotime. *Also see Supplementary Fig. 2, Supplementary Fig. 3, and Supplementary Table 2A*.

### Cell subclusters and transcriptomic profiles of the IFP

We next aimed to determine cell subclusters of each major cell type identified within IFPs (n=21). Subclustering of fibroblasts determined five subclusters within IFPs, with proportions ranging from 30.32% (subcluster 0) to 9.63% (subcluster 4) (total nuclei: 32,862, resolution 0.2) (Fig. 1E; Supplementary Fig. 2B). Unique transcriptomic profiles for each fibroblast subcluster were generated by identifying differentially expressed genes (DEGs) in one subcluster as compared to others. These unique profiles ranged from 225 genes (subcluster 2) to 43 genes (subcluster 3) [q<0.05, Log2Fold Change (FC)≥0.5, min.pct≥0.25] (Supplementary Fig. 3; Supplementary Table 2A). Top 5 genes for each fibroblast subcluster, based on decreasing Log2FC, included subcluster 0: FBN1, PXDNL, PIEZO2, GUCY1A2, and PTGIS; subcluster 1: ABCA10, KCND2, ABCA6, ABCA8, and ABCA9; subcluster 2: CRTAC1, CLIC5, PRG4, FN1, and SEMA3A; subcluster 3: GPAM, FABP4, F13A1, FTL, and FTH1; subcluster 4: MGAT4C, GRIP1, BTBD11, IGF1, and C1GALT1 (Fig. 1F; Supplementary Fig. 2C, Supplementary Table 2A). Pseudotime trajectory analysis using SCORPIUS v1.0.9^10^ showed a clear transition of fibroblast subclusters from subcluster 0 to subcluster 2, through subclusters 3, 1, and 4 (Fig. 1G). We identified genes that predicted the ordering of fibroblast subclusters along pseudotime, ranked by importance scores. There were distinct changes in module expression profiles, corresponding to transcriptomic profiles of fibroblast subclusters (Fig. 1G). Additionally, top markers of subcluster 0 (FBN1), subcluster 1 (ABCA10), subcluster 2 (CRTAC1), and subcluster 4 (MGAT4C) showed peak expression in subclusters 0, 1, 2, and 4, respectively, along pseudotime trajectory (Fig. 1G).

Using a similar bioinformatic strategy, we uncovered four subclusters of macrophages within IFPs with proportions ranging from 49.85% (subcluster 0) to 8.06% (subcluster 3) (total nuclei: 14,351, resolution 0.1) (Fig. 2A; Supplementary Fig. 4A). Each macrophage subcluster had unique transcriptomic markers spanning from 232 genes (subcluster 1) to 74 genes (subcluster 3) (Supplementary Fig. 5; Supplementary Table 2B). Top 5 genes for each macrophage subcluster based on decreasing Log2FC (q<0.05, Log2FC≥0.5, min.pct≥0.25), included subcluster 0: P2RY14, MAN1A1, FGF13, DSCAML1, and PID1; subcluster 1: FN1, TPRG1, KCNQ3, PDE3A, and FMNL2; subcluster 2: TTN, STAB1, AHNAK, RBM25, and PRRC2C; subcluster 3: NOVA1, ADH1B, DLC1, FBN1, and NEGR1 (Fig. 2B; Supplementary Fig. 4B, Supplementary Table 2B).

**Figure 2.**
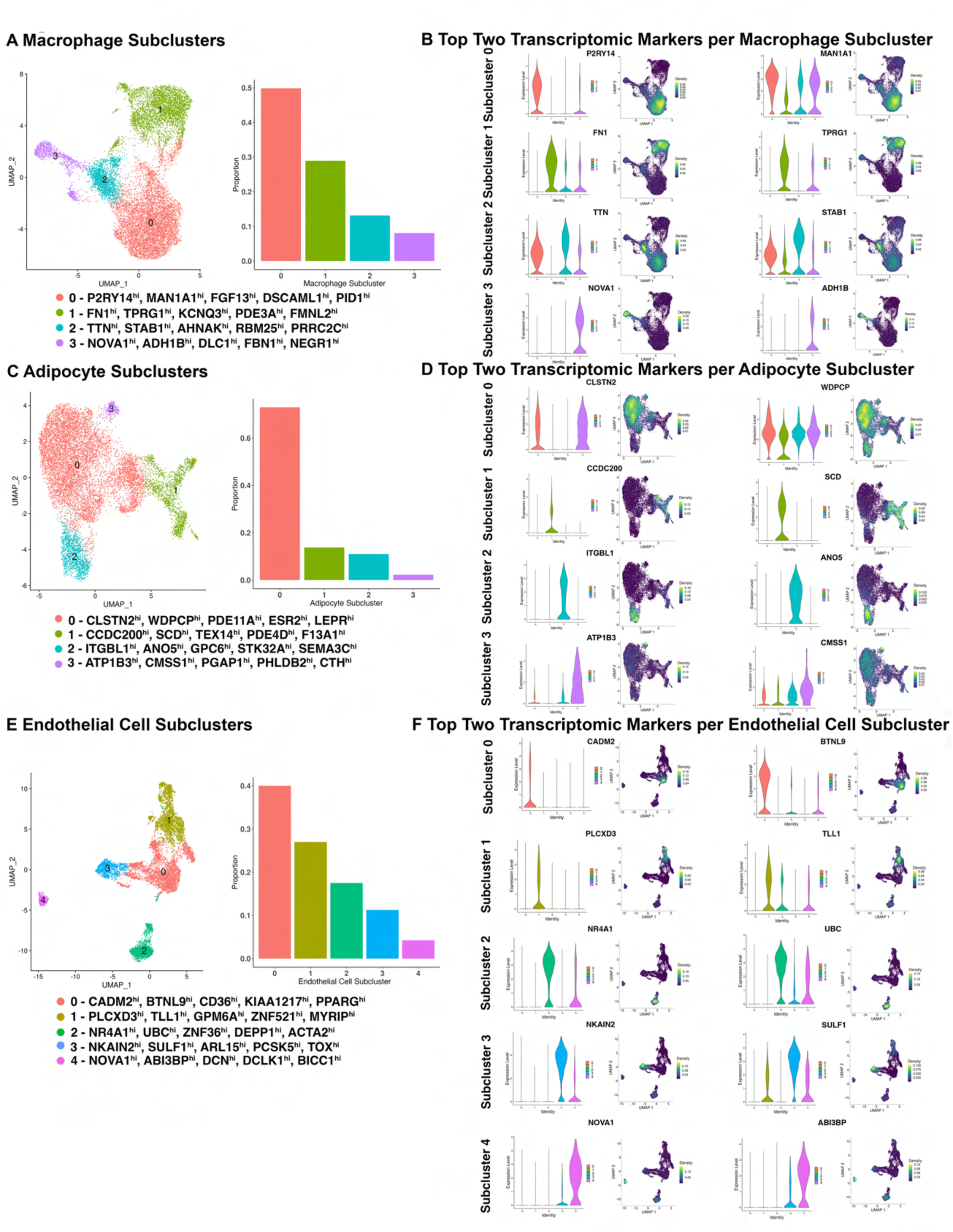
Macrophage, Adipocyte and Endothelial Cell Subclusters in Infrapatellar Fat Pad Identified by snRNA-seq (n=21). (A, C, E) UMAPs and bar plots depicting cell type subclusters and their respective proportions. Top 5 differentially expressed genes within each cluster are indicated. (B, D, F) Violin plots and nebulosa plots depicting the expression of the top two unique transcriptomic markers within each macrophage, adipocyte, and endothelial cell subcluster. Nebulosa plots present the expression density from dark purple (low) to yellow (high). *Also see Supplementary Fig. 4-9 and Supplementary Tables 2B-D*.

Subclustering of adipocytes identified four subclusters, with proportions ranging from 72.97% (subcluster 0) to 2.25% (subcluster 3) (total nuclei: 12,111, resolution 0.08) (Fig. 2C; Supplementary Fig. 6A). All adipocyte subclusters had unique transcriptomic markers, ranging from 160 genes (subcluster 2) to 89 genes (subcluster 0) (Supplementary Fig. 7; Supplementary Table 2C). Top 5 genes for each adipocyte subcluster based on decreasing Log2FC (q<0.05, Log2FC≥0.5, min.pct≥0.25), included subcluster 0: CLSTN2, WDPCP, PDE11A, ESR2, and LEPR; subcluster 1: CCDC200, SCD, TEX14, PDE4D, and F13A1; subcluster 2: ITGBL1, ANO5, GPC6, STK32A, and SEMA3C; subcluster 3: ATP1B3, CMSS1, PGAP1, PHLDB2, and CTH (Fig. 2D; Supplementary Fig. 6B, Supplementary Table 2C).

Finally, we identified five endothelial cell subclusters within IFPs, with proportions ranging from 40% (subcluster 0) to 4.22% (subcluster 4) (total nuclei: 8,944, resolution 0.05) (Fig. 2E; Supplementary Fig. 8A). Each endothelial cell subcluster had unique transcriptomic markers, ranging from 225 genes (subcluster 3) to 153 genes (subcluster 0) (Supplementary Fig. 9, Supplementary Table 2D). Top 5 genes for each endothelial cell subcluster based on decreasing Log2FC (q<0.05, Log2FC≥0.5, min.pct≥0.25), included subcluster 0: CADM2, BTNL9, CD36, KIAA1217, and PPARG; subcluster 1: PLCXD3, TLL1, GPM6A, ZNF521, and MYRIP; subcluster 2: NR4A1, UBC, ZFP36, DEPP1, and ACTA2; subcluster 3: NKAIN2, SULF1, ARL15, PCSK5, and TOX; subcluster 4: NOVA1, ABI3BP, DCN, DCLK1, and BICC1 (Fig. 2F; Supplementary Fig. 8B, Supplementary Table 2D).

Fibroblast subclusters identified within the IFP mapped to fibroblast populations identified within other adipose tissues (Supplementary Fig. 10A)^11–14^. Specifically, in combined IFP and synovial tissue, Tang et al.^7^ uncovered four fibroblast subsets, in line with fibroblast subclusters we revealed within IFP only (Supplementary Fig. 10A). Fibroblasts identified in IFP can also be referred to as preadipocytes, adipocyte stem and progenitor cells (ASPCs), and mesenchymal stromal cells (MSCs), as they mapped to populations using this nomenclature^11^. In addition, comparing our snRNA-seq data of IFP with reported white adipose tissue markers^11^, we mapped the macrophage, adipocyte, and endothelial cell subclusters we identified (Supplementary Fig. 10B-D).

### Spatial profiling of major cell types, fibroblast subclusters and transcriptomic profiles of KOA-IFP

After revealing major cell types and subtypes, and their transcriptomic profiles, within the IFP, we sought to resolve their spatial distribution using spatial transcriptomics (Fig. 3A). We analyzed n=12 KOA-IFPs patient-matched to tissues analyzed by snRNA-seq, using Visium CytAssist spatial transcriptomics technology (Supplementary Table 1). Spatial data were clustered and annotated independently from snRNA-seq data. We confirmed that fibroblasts, macrophages, adipocytes, and endothelial cells are the major cell populations within IFPs, and fibroblasts are distributed across the IFP, alongside macrophages, adipocytes, and endothelial cells (Fig. 3B; Supplementary Fig. 11A).

**Figure 3.**
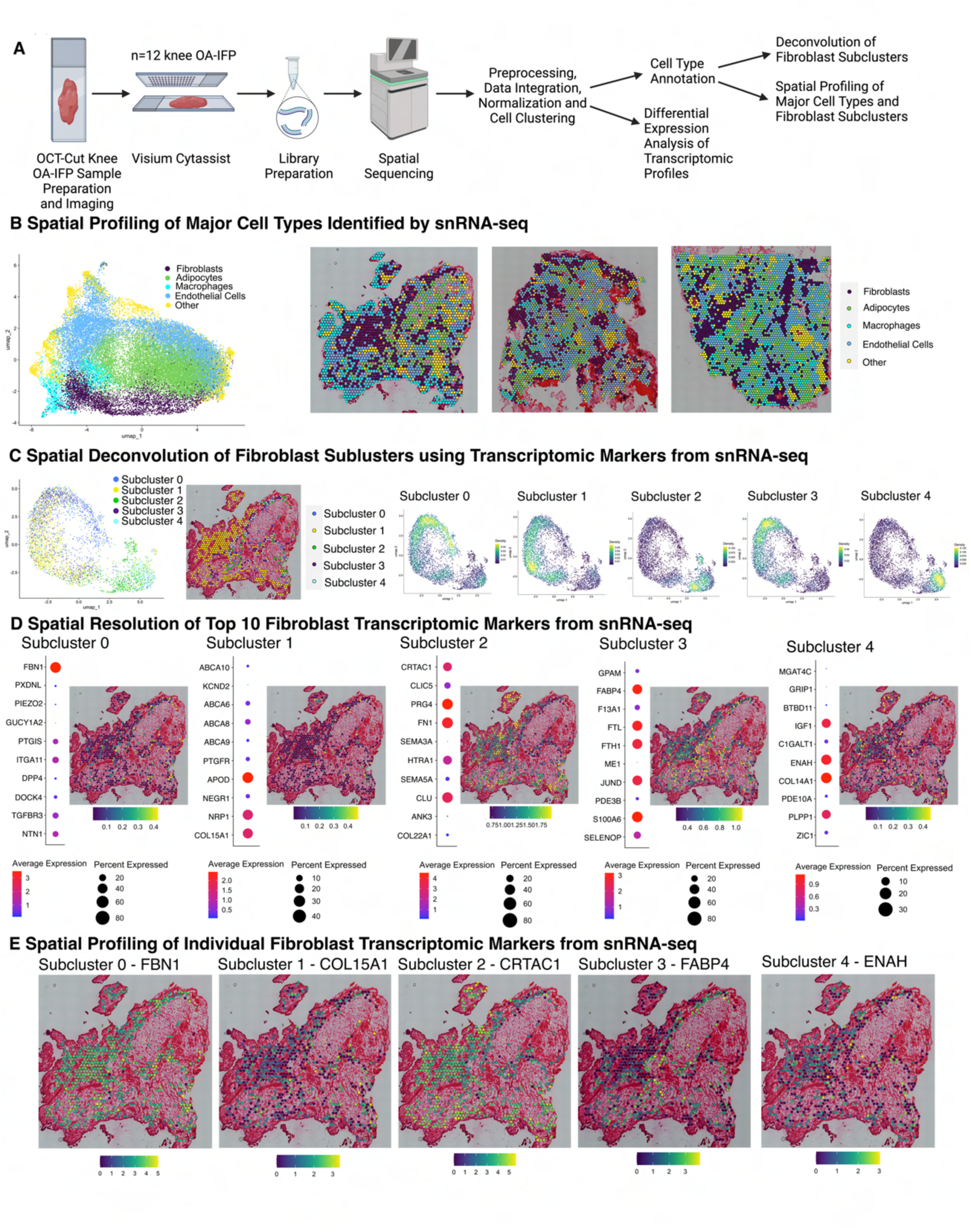
Spatial Profiling of Major Cell Types Identified by snRNA-seq Data. A) Schematic workflow to spatially resolve cell types within IFPs using spatial sequencing. Created with BioRender.com. B) UMAP (left) of spatial sequencing data of n=12 KOA-IFP. Spatial resolution of cell populations identified using spatial sequencing across multiple IFPs (right). C) UMAP (left) of spatial deconvolution of fibroblast subclusters across n=12 IFP using snRNA-seq. Spatial resolution of deconvolved fibroblast subclusters within a representative IFP sample (middle). Nebulosa plots (right) displaying the module scores of the average gene expression of the top ten transcriptomic markers of each deconvolved fibroblast subcluster. Module scores within nebulosa plots range from yellow (high) to dark purple (low). D) Dot plots show the average expression of the top 10 transcriptomic markers within each fibroblast subcluster, from blue (low average expression) to red (high average expression). Spatial plots show the resolution of the module scores of co-expression of the top ten transcriptomic markers of each fibroblast subcluster within a representative IFP, from blue (low module score) to yellow (high module score). E) Spatial resolution of individual transcriptomic markers of each fibroblast subcluster include: Subcluster 0 – FBN1 (DEG 1), Subcluster 1 – COL15A1 (DEG 10), Subcluster 2 – CRTAC1 (DEG 1), Subcluster 3 – FABP4 (DEG 2), Subcluster 4 – ENAH (DEG 5). *Also see Supplementary Fig. 11*.

Fibroblasts from spatial transcriptomic data were deconvoluted using snRNA-seq fibroblast subcluster annotations. Each fibroblast subcluster was identified within IFPs in areas independently identified as fibroblasts by spatial transcriptomics (Fig. 3C). Interestingly, fibroblast subcluster 1 tended to be found in the periphery of IFPs, while all other subclusters tended to co-locate deeper in the tissues (Fig. 3C). Subsequently, we spatially resolved transcriptomic profiles of each identified fibroblast subcluster, through co-expression of the top 10 genes of each subcluster (Fig. 3D). We also visualized one of the top DEGs of each fibroblast subcluster including: subcluster 0: FBN1; subcluster 1: COL15A1; subcluster 2: CRTAC1; subcluster 3: FABP4; subcluster 4: ENAH (Fig. 1F; Fig. 3E; Supplementary Fig. 2C; Supplementary Fig. 11B).

### Cell-cell interactions within the KOA-IFP

Since fibroblasts had the largest proportion of nuclei, and all major cell types were spatially organized proximal to fibroblasts across IFPs, we investigated putative interactions that occur between fibroblasts and all other major cell types within our snRNA-seq data. Using CellChat v2.0^14^, we identified major ligand-receptor cell signalling pathways, with macrophages, adipocytes, and endothelial cells as “sender” cells expressing ligands and fibroblasts as “receiver” cells expressing receptors. Multiple cell communications to fibroblasts were identified, with the highest proportion of ligand-receptor pairs originating from adipocytes, and the highest proportion of interactions occurring with fibroblast CD44 and ITGAV/ITGB8 receptors (Fig 4A, Supplementary Table 3).

**Figure 4.**
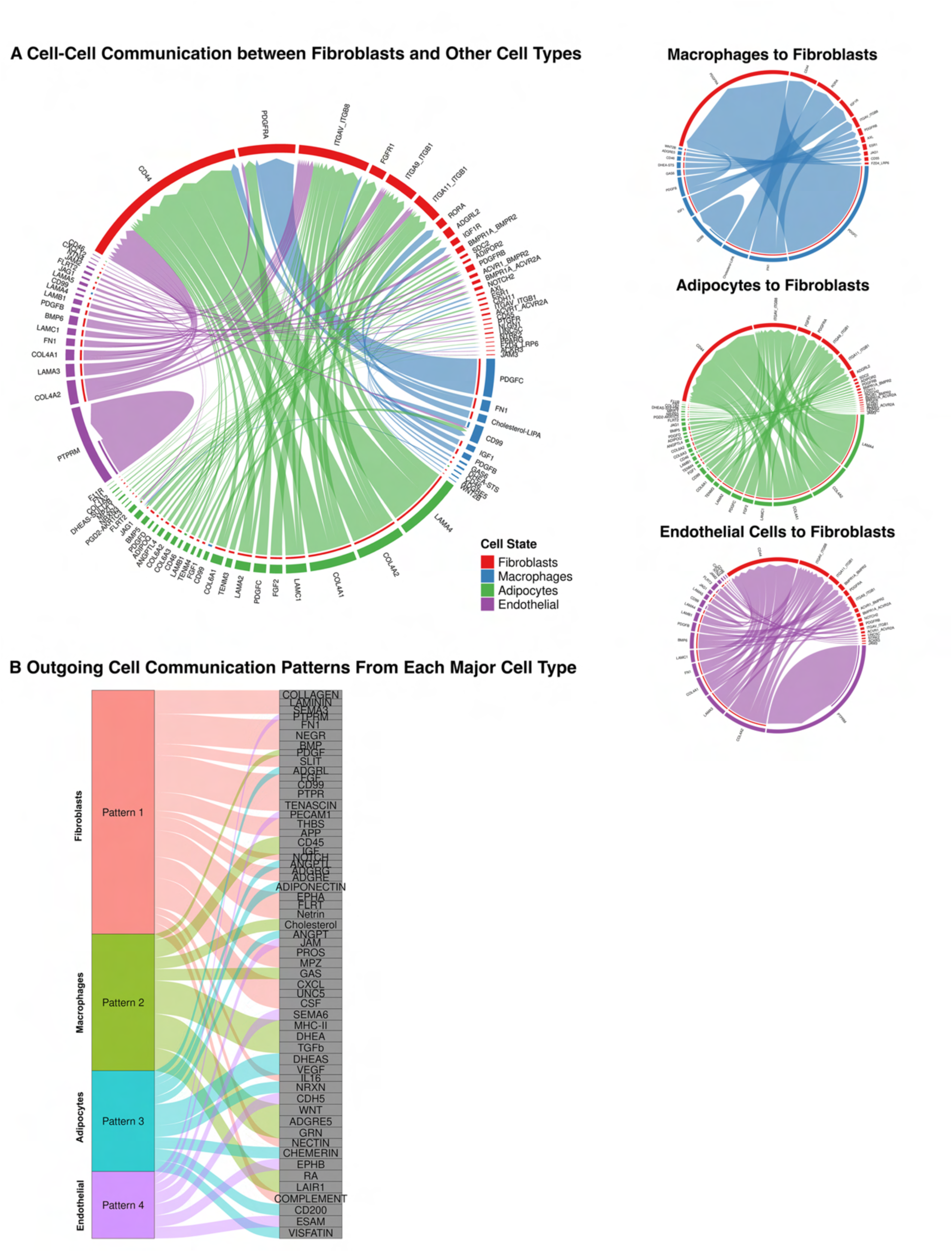
Putative Cell-Cell Communication Patterns Between Fibroblasts and Major Cell Types. A) Chord diagram (left) showing cell-cell communication between fibroblasts and other identified major cell types using snRNAseq data from n=21 infrapatellar fat pad samples. Fibroblasts are the “receiver” cell type, expressing receptors, and all other cell types are the “sender” cells, expressing ligands. Individual chord diagrams (right) showing cell-cell communication between fibroblasts and individual other cell types. B) River plot displays outgoing signalling patterns from each identified cell type. *Also see Supplementary Table 3*.

Additional investigations uncovered outgoing signalling patterns from all major cell types, without defining “receivers” or “senders”. Regardless of directionality, fibroblasts, macrophages, adipocytes, and endothelial cells had unique outgoing signalling patterns composed of multiple signaling pathways (Fig. 4B). This suggests these four cell types inter-communicate through multiple outgoing signals including collagen, PECAM1, PDGF, and ANGPTL signalling, which are known to stimulate regeneration and repair, extracellular matrix (ECM) deposition, and angiogenesis^16–19^.

### Transcriptomic differences in fibroblasts within the IFP based on KOA status

Following the identification of major cell types, subtypes, and their transcriptomic profiles, within the IFP, we investigated whether there were differences in cell composition based on KOA status. Fibroblasts, having the highest proportion of cells contributing to the IFP, were selected for subsequent analysis. Bioinformatic analysis of snRNA-seq data determined alterations in fibroblast subclusters within KOA-IFP (n=15) compared to healthy control donor IFPs (n=6) (Supplementary Table 1). We found the presence of all fibroblast subclusters across individual KOA-IFPs and healthy control donor IFPs (Fig. 5A). However, subcluster 3 had a higher proportion of nuclei contributed from healthy control donor IFPs compared to KOA-IFP (q=0.032, multiple unpaired t-tests with FDR correction) (Fig. 5B). Additionally, 38 DEGs (20 upregulated, 18 downregulated) were identified within KOA-IFPs compared to healthy control donor IFPs, with the top 5 upregulated genes: FN1, ZNF385B, PRG4, ANKH, and KAZN, and the top 5 downregulated genes: JUN, ABCA1, VIM, GSN, and SAMD4A (Fig. 5C; Supplementary Table 4A).

**Figure 5.**
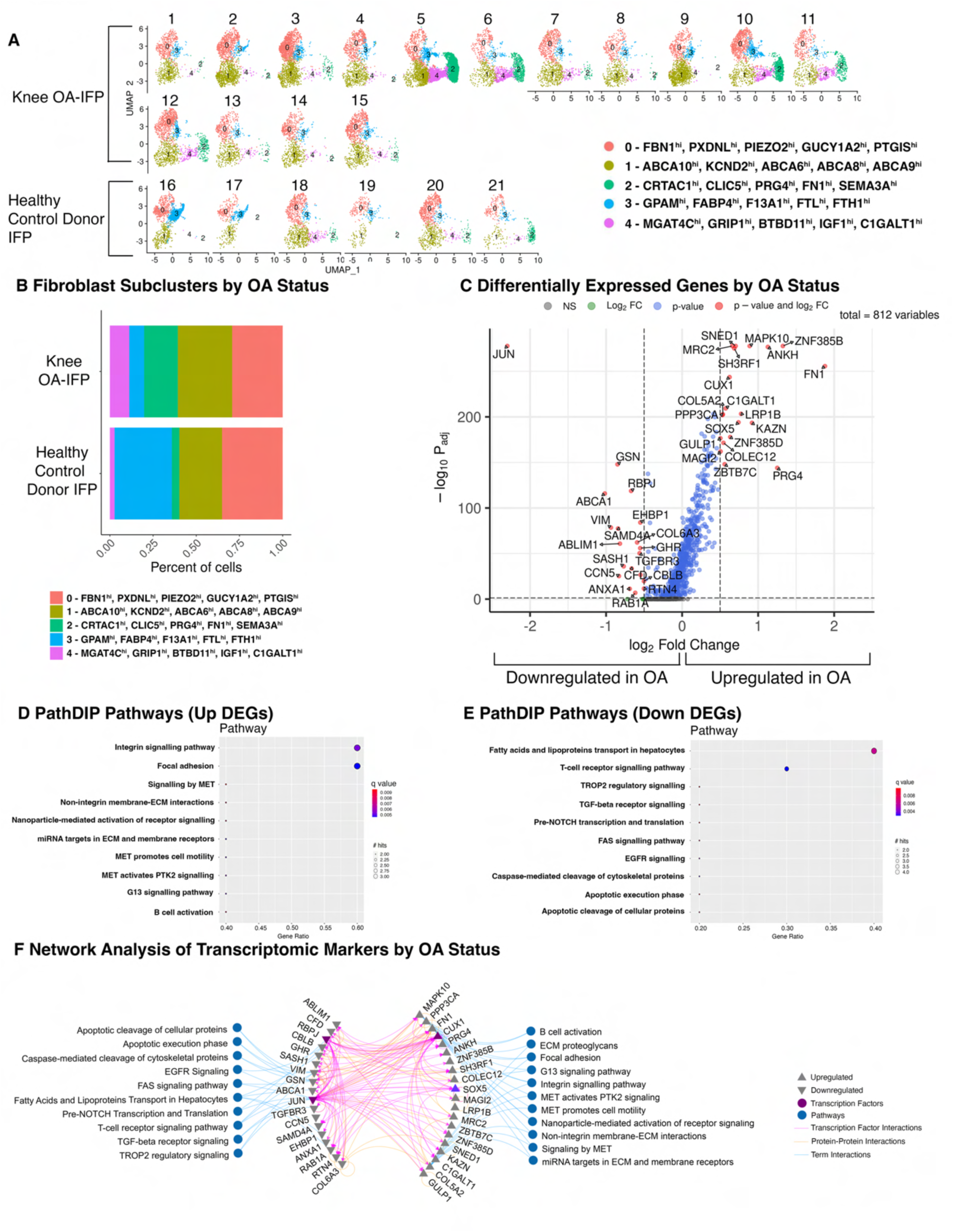
Differences in Infrapatellar Fat Pad (IFP) Fibroblast Subclusters Based on Osteoarthritis (OA) Status. A) UMAPs of fibroblast subclusters within each of the n=21 IFPs. Fifteen IFPs (top) have late-stage knee (K)OA while six (bottom) are healthy control donor IFPs. B) Stacked bar plot displaying the proportion of fibroblast subclusters within KOA-IFPs compared to healthy control donor IFPs. Fibroblast subcluster 3 was shown to have significantly more nuclei from healthy control donor IFPs compared to KOA-IFPs (q=0.032). Fibroblast subcluster proportions were analyzed by arcsin-transforming the absolute proportions across all samples and performing multiple unpaired t-tests with FDR correction using the Benjamini, Krieger and Yekutieli two stage step up method. Top 5 differentially expressed genes (DEGs) within each cluster are indicated. C) Volcano plot showing the log2 fold change (FC) of DEGs in KOA-IFPs compared to healthy control donor IFPs. DEGs were defined by a minimum of 50% of nuclei expressing a marker, with a log2FC > 0.5, q < 0.05 are upregulated while genes with a log2FC < -0.5, q < 0.05 are downregulated. D) Enriched PathDIP pathways associated with the upregulated genes from KOA-IFPs compared to healthy control donor IFPs. E) Enriched PathDIP pathways associated with the downregulated genes from KOA-IFPs compared to healthy control donor IFPs. F) Interaction network showing protein-protein interactions and transcription factor-gene interactions with enriched pathDIP pathways linked to DEGs. *Also see Supplementary Tables 4A-C*.

PathDIP analysis (pathDIP v5, https://ophid.utoronto.ca/pathDIP)^20^ identified enriched pathways associated with DEGs in KOA-IFPs compared to healthy control donor IFPs (Fig 5D,E; Supplementary Table 4B,C). Network analysis revealed connections between DEGs and enriched pathways from KOA- vs. healthy control donor-IFPs, as well as putative interactions between transcription factors (Fig. 5F). Of note, transcription factors SOX5 and CUX1 were upregulated in KOA-IFP compared to healthy control donor IFP while JUN and RBPJ were downregulated, with putative interactions between JUN and CUX1, SOX5 and JUN, and RBPJ and CUX1 (Fig. 5C,F).

### Sex-based transcriptomic differences in fibroblasts within the KOA-IFP

Female sex is a significant risk factor for developing KOA. Compared to males, post-menopausal females have a higher prevalence of KOA and report increased pain, inflammation and mobility issues associated with KOA^9^. However, how sex impacts major cell populations and subclusters within KOA-IFP remains undefined.

To understand how sex influences fibroblasts within KOA-IFPs, we performed bioinformatic analyses of snRNA-seq data of female (n=8) compared to male (n=7) KOA-IFPs (Supplementary Table 1). We observed no significant differences in the presence or proportions of fibroblast subclusters (q<0.05, multiple unpaired t-tests with FDR correction) (Fig. 6A,B). However, we identified clear differences in transcriptomic profiles of fibroblasts within female versus male KOA-IFPs (Fig. 6C). We determined 105 DEGs in female compared to male KOA-IFPs: 35 upregulated and 70 downregulated genes. The top 5 upregulated genes include CLU, HTRA1, PRG4, TMEM196, and FN1 and the top 5 downregulated genes include UTY, USP9Y, LAMA2, AFF3, and NAV3 (Fig. 6C; Supplementary Table 5A).

**Figure 6.**
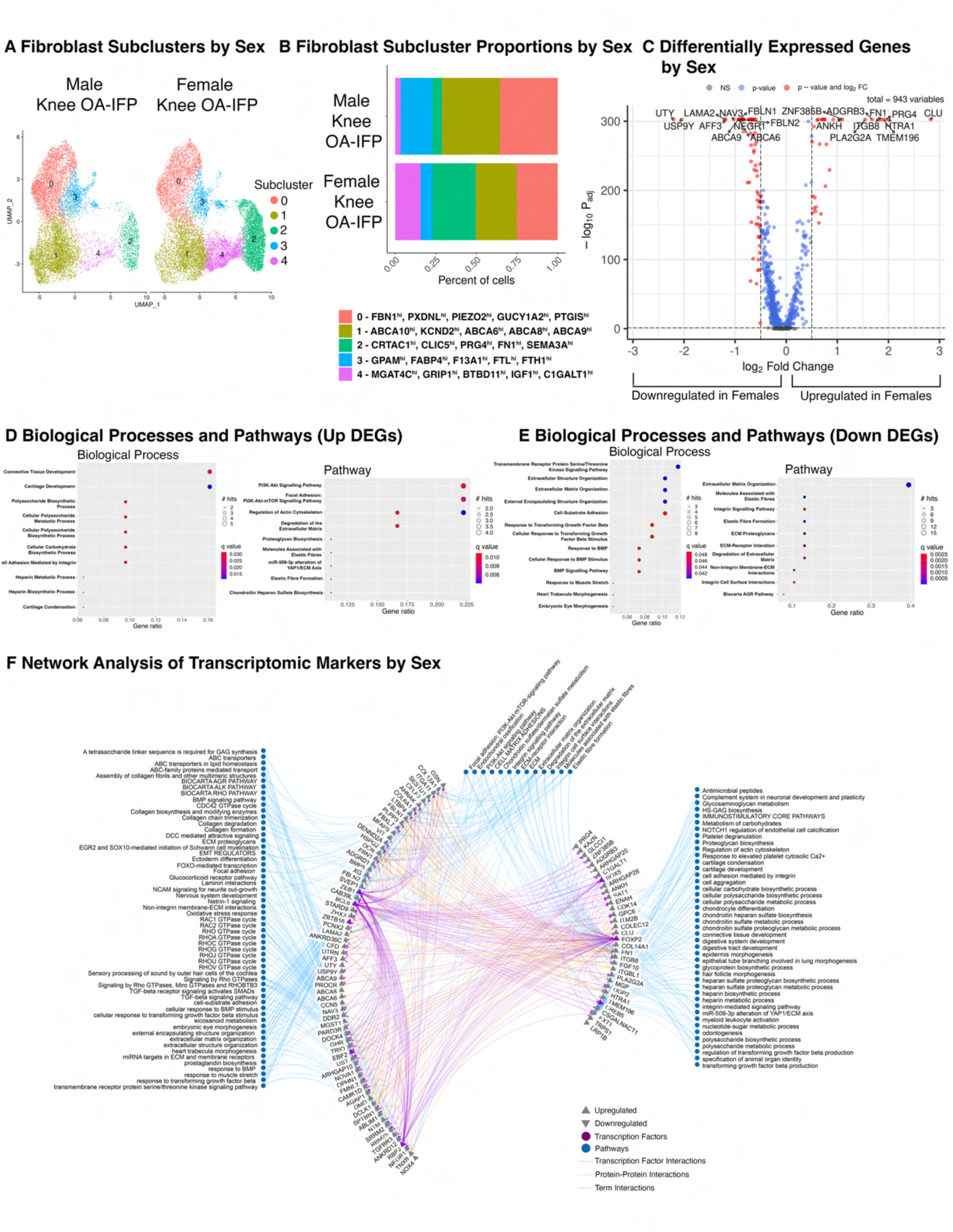
Differences in Knee Osteoarthritis (KOA)-Infrapatellar Fat Pad (IFP) Fibroblast Subclusters Based on Sex. A) UMAPs of fibroblast subclusters within n=15 KOA-IFPs split by sex (n=8 female KOA-IFP, n=7 male KOA-IFP). B) Stacked bar plot displaying the proportion of fibroblast subclusters within female KOA- IFPs compared to male KOA-IFPs. No significant differences were found in proportions of nuclei contributed to subclusters from KOA-IFPs from female versus male study participants (q<0.05). Fibroblast subcluster proportions were analyzed by arcsin-transforming the absolute proportions across all samples and performing multiple unpaired t-tests with FDR correction using the Benjamini, Krieger and Yekutieli two stage step up method. Top 5 differentially expressed genes (DEGs) within each cluster are indicated. C) Volcano plot showing the log2 fold change (FC) of DEGs in female KOA-IFP samples compared to male KOA-IFP samples. DEGs were defined by a minimum of 50% of nuclei expressing a marker, with a log2FC > 0.5, q<0.05 are upregulated while genes with a log2FC < -0.5, q<0.05 are downregulated. D) GO biological processes and pathDIP pathways enriched in the upregulated genes of female compared to male KOA- IFPs. E) GO biological processes and pathDIP pathways enriched for the downregulated genes within female compared to male KOA-IFPs. F) Interaction network showing protein-protein interactions and transcription factor-gene interactions with enriched biological processes or pathDIP pathways linked to DEGs. *Also see Supplementary Tables 5A-E*.

Gene Ontology and pathDIP analysis (pathDIP v5, https://ophid.utoronto.ca/pathDIP)^20,21^ identified enriched biological processes and pathways associated with DEGs of female compared to male KOA-IFP (Fig 6D,E; Supplementary Tables B-E). Network analysis was performed to determine connections between DEGs and associated biological processes and pathways, with putative interactions between transcription factors (Fig. 6F). Notably, transcription factors SOX5, FOXP2, and CREB5 were upregulated within female versus male KOA-IFPs, while ZEB1, BCL6, ZBTB16, EBF2, and RBPJ were downregulated, with putative interactions between transcription factor pairs ZBTB16 and BCL6, and ZEB1 and BCL6 (Fig. 6C, F).

### Transcriptomic differences of fibroblasts within KOA-IFP based on obesity status

Obesity is a prominent risk for developing KOA, however, how obesity affects fibroblasts within KOA-IFP remains to be understood. To discern how obesity status influences fibroblasts of KOA-IFPs, we performed bioinformatic analysis of snRNA-seq data comparing fibroblast subclusters within obese (n=8; BMI 30 kg/m^2^-40 kg/m^2^) versus normal BMI KOA-IFPs (n=7; BMI 18.5 kg/m^2^-24.9 kg/m^2^) (Supplementary Table 1). We found no significant differences in the presence or proportions of fibroblast subclusters between normal and obese BMI KOA-IFPs (q<0.05, multiple unpaired t-tests with FDR correction) (Fig. 7A,B). However, 21 DEGs were identified; 10 upregulated, 11 downregulated (Fig. 7C; Supplementary Table 6A). Upregulated genes from obese versus normal BMI KOA-IFPs included: HMCN1, LHFPL6, ABCA6, PRRX1, ABCA8, ITGA11, DST, STARD9, SRRM2, and FBLN1, while downregulated genes included: PRG4, CLU, FN1, ITGB8, FKBP5, UGP2, TIMP3, GPC6, ZBTB16, ITM2B, and CREB5 (Fig. 7C; Supplementary Table 6A).

**Figure 7.**
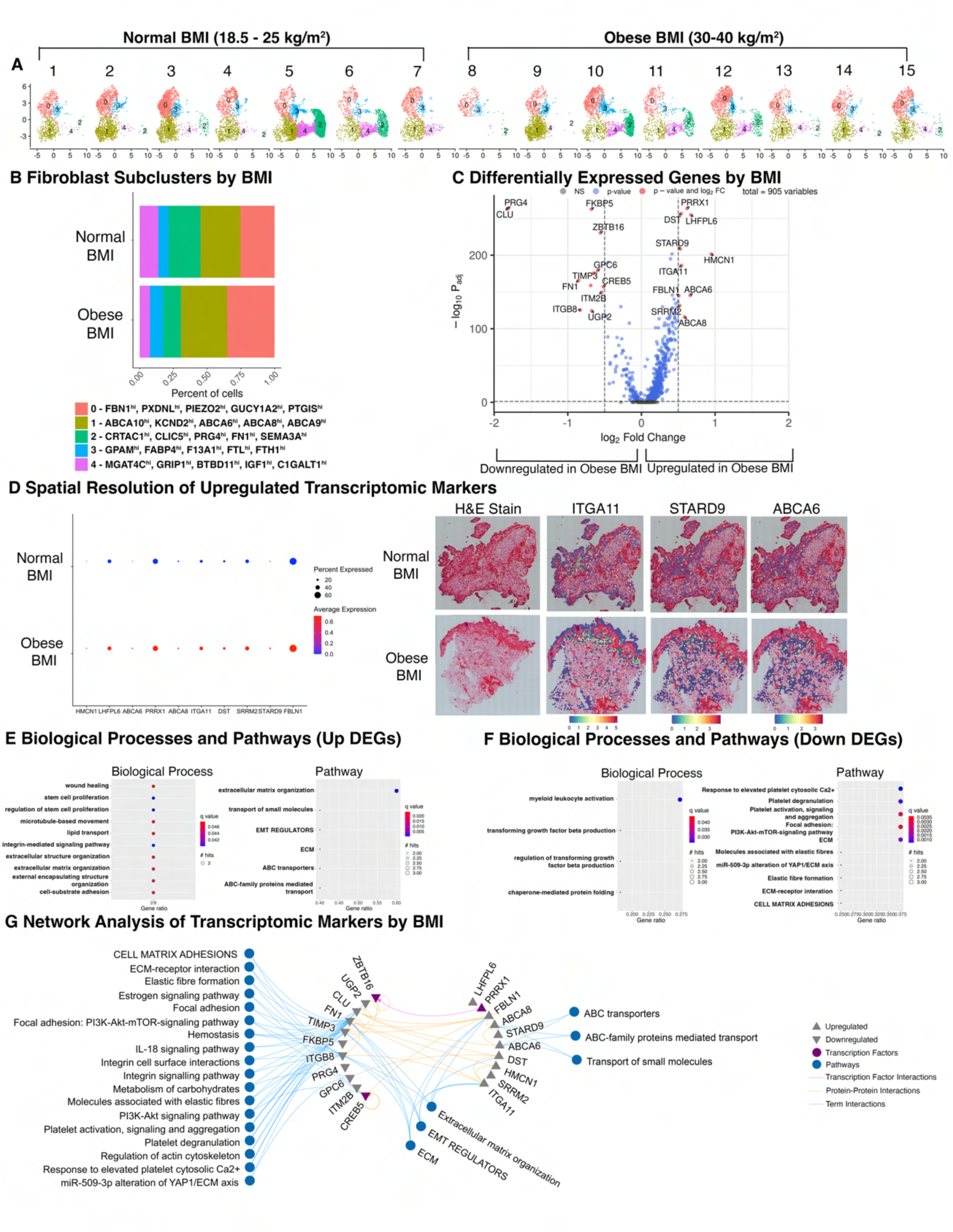
Differences in Knee Osteoarthritis (KOA)-Infrapatellar Fat Pad (IFP) Fibroblast Subclusters Based on Obesity Status. A) UMAPs of fibroblast subclusters within each of the n=15 KOA-IFPs. Seven samples (left) have normal BMI (18.5 – 25 kg/m^2^) while eight samples (right) have an obese BMI (30 – 40 kg/m^2^). B) Stacked bar plot displaying the proportion of fibroblast subclusters within obese BMI KOA-IFPs compared to normal BMI KOA-IFPs. No significant differences were found in proportions of nuclei contributed to subclusters from KOA-IFPs from obese versus normal BMI study participants (aq<0.05). Fibroblast subcluster proportions were analyzed by arcsin-transforming the absolute proportions across all samples and performing multiple unpaired t-tests with FDR correction using the Benjamini, Krieger and Yekutieli two stage step up method. Top 5 differentially expressed genes (DEGs) within each cluster are indicated. C) Volcano plot showing the log2 fold change (FC) of genes that are uniquely differentially expressed in obese BMI compared to normal BMI KOA-IFPs. DEGs were defined by a minimum of 50% of nuclei expressing a marker, genes with a log2FC > 0.5, q < 0.05 are upregulated while genes with a log2FC < -0.5, q < 0.05 are downregulated. D) Dot plot (left) with average expression and percent of population expressing each DEG based on obesity status identified in snRNA-seq data within spatial sequencing data. Spatial resolution (right) of genes significantly upregulated within obese BMI compared to normal BMI KOA-IFPs in both snRNA-seq and spatial sequencing, visualized within spatial sequencing data. E) GO biological processes and pathDIP pathways enriched for the upregulated genes from obese BMI compared to normal BMI KOA-IFP samples. F) GO biological processes and pathDIP pathways enriched for downregulated genes from obese compared to normal BMI KOA-IFP samples. G) Interaction network showing protein-protein interactions and transcription factor-gene interactions with enriched biological processes or pathDIP pathways linked to DEGs. *Also see Supplementary Tables 6A-F*.

We next spatially resolved the 10 upregulated DEGs identified in obese compared to normal BMI KOA-IFPs using spatial transcriptomics (n=12; n=6 normal and n=6 obese BMI KOA-IFP). All 10 genes upregulated in obese BMI KOA-IFPs identified in snRNA-seq data were also upregulated within spatial sequencing data based on Log2FC (Fig 7D, Supplementary Table 6B). Of the 10 upregulated genes, ITGA11, STARD9, and ABCA6, were significantly upregulated in spatial transcriptomic analyses (q<0.05; Fig. 7D, Supplementary Table 6B).

Gene Ontology and pathDIP analysis (pathDIP v5, https://ophid.utoronto.ca/pathDIP)^20,21^ identified enriched biological processes and pathways associated with DEGs in obese compared to normal BMI KOA-IFPs (Fig 7E,F; Supplementary Tables 6C-F). Network analysis determined connections between DEGs, and enriched biological processes and pathways, with putative interactions between transcription factors (Fig. 7G). Of note, the transcription factor PRRX1 was upregulated in obese versus normal BMI KOA-IFPs while ZBTB16 and CREB5 were downregulated, with a putative link between PRRX1 and ZBTB16 (Fig. 7C,G). We also identified enriched pathways related to metabolic functions, including lipid transport and carbohydrate metabolism (Fig 7G).

### Differences in metabolite levels of fibroblasts isolated from obese and normal BMI KOA-IFPs

Since metabolic deregulation is a key element of obesity and transcriptional differences in fibroblasts by obesity status had enriched pathways linked to metabolism, we sought to elucidate if alterations in supernatant metabolite levels of cultured fibroblasts from obese (n=5) compared to normal BMI KOA-IFPs (n=5) existed using targeted metabolomics (Fig. 8A). Overall, 445 metabolites were detected. From this, 5 metabolites were found at significantly different levels (p<0.05) from obese versus normal BMI KOA-IFPs, including upregulated: triglycerides (TG) (54:2) and TG (50:2), C18:2 (linoleic acid), and choline; downregulated: homoarginine (Fig. 8B, Supplementary Table 7A).

**Figure 8.**
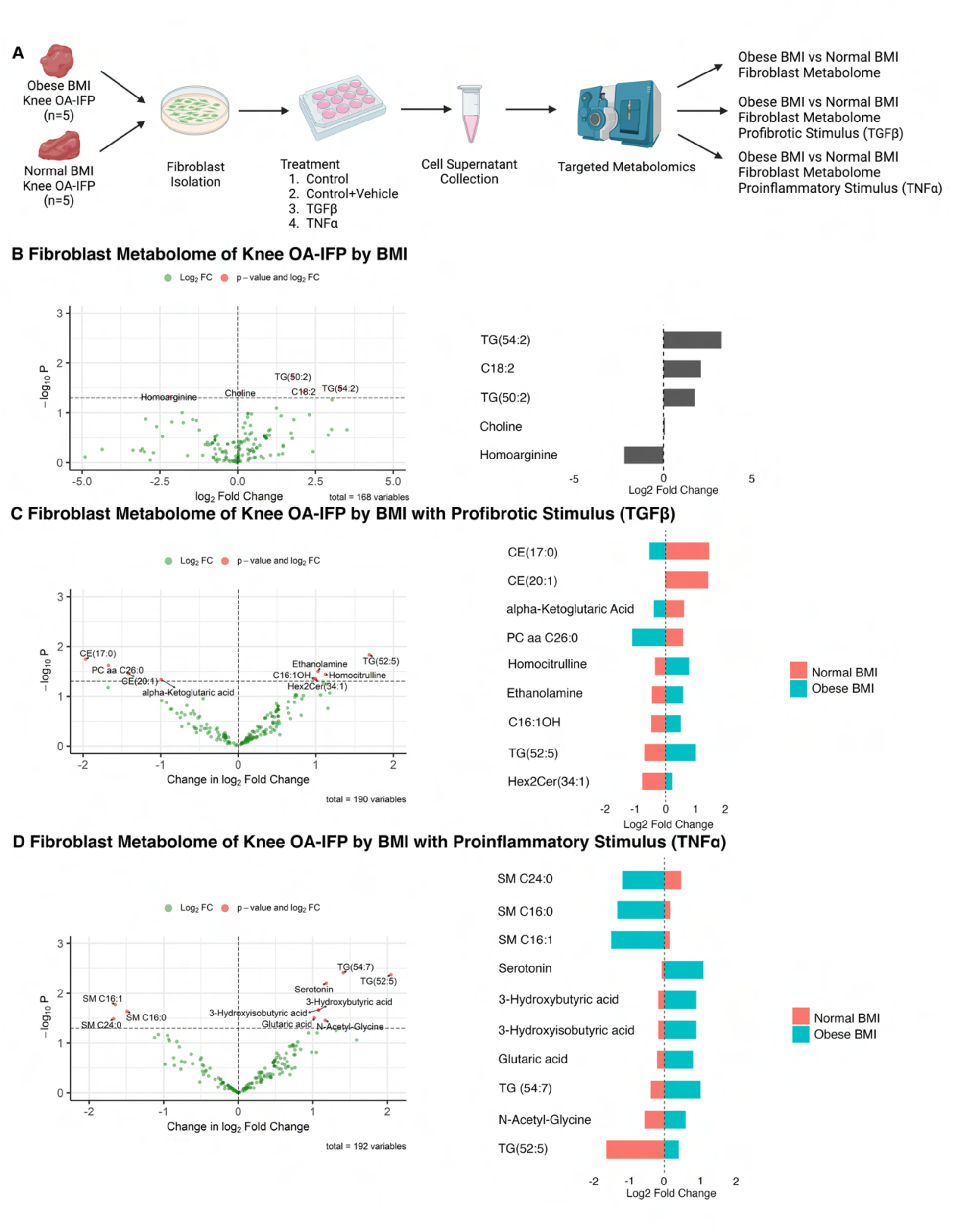
Differences in Metabolite Secretome in Fibroblasts from Knee Osteoarthritis (KOA)- Infrapatellar Fat Pad (IFP) Based on Obesity Status. A) Schematic workflow to identify differences in the metabolite levels in the supernatants of fibroblast cultures of obese compared to normal BMI KOA-IFP, stimulated with or without profibrotic or proinflammatory stimulus. Created with BioRender.com. B) Violin plot (left) and bar plot (right) showing the log2 fold change (FC) of metabolites differentially found in supernatants of fibroblast cultures from obese compared to normal BMI KOA-IFPs. Positive log2FC indicates an increase in the metabolite level within supernatants of cultures from obese compared to normal BMI KOA-IFP fibroblasts. C) Volcano plot (left) showing the change in log2FC of metabolites differentially found in supernatants of obese compared to normal BMI KOA-IFP fibroblast cultures when treated with TGFβ compared to vehicle control. Positive values indicate a greater upregulation of metabolite level in obese vs normal BMI KOA-IFP fibroblast culture supernatants when treated with TGFβ vs vehicle control. Bar plot (right) showing the log2FC of metabolites differentially found in supernatants of obese compared to normal BMI KOA-IFP fibroblast cultures when treated with TGFβ compared to vehicle control, with positive values indicating an upregulation in levels found in supernatant. D) Volcano plot (left) showing the change in log2FC of metabolites differentially found in supernatants of obese compared to normal BMI KOA-IFP fibroblast cultures when treated with TNFα compared to vehicle control. Positive values indicate a greater upregulation of metabolite level in obese vs normal BMI KOA-IFP fibroblast culture supernatants when treated with TNFα vs vehicle control. Bar plot (right) showing the log2FC of metabolites differentially found in supernatants of obese compared to normal BMI KOA-IFP fibroblast cultures treated with TNFα compared to vehicle control, with positive values indicating an upregulation in secretion. *Also see Supplementary Tables 7A-C*.

Within KOA joints, TGFβ and TNFα are found in synovial fluid bathing joint tissues, including the IFP. Since these molecules are major drivers of fibrotic and inflammatory pathologies associated with KOA^22,23^, we investigated how metabolite levels in culture supernatants of obese versus normal BMI KOA- IFP fibroblasts stimulated by TGFβ or TNFα were modified, compared to stimulation by vehicle control (PBS). When comparing fibroblast cultures from obese versus normal BMI KOA-IFPs, 9 metabolites were significantly differentially detected (p<0.05) in response to TGFβ treatment, compared to vehicle control (Fig. 8C, Supplementary Table 7B). Mean levels of homocitrulline, ethanolamine, C16:1OH (2-hydroxypalmitoleylcarnitine), TG (52:5), and Hex2Cer(34:1) (di-hexosyl ceramide) were increased by TGFβ- treated fibroblasts from obese BMI KOA-IFPs versus vehicle control, and decreased in normal BMI KOA- IFPs. In contrast, mean levels of cholesteryl esters (17:0) and (20:1), alpha-ketoglutaric acid, and phosphatidylcholine diacyl (PCaa)C26:0 were increased in TGFβ-treated fibroblast cultures from normal BMI KOA-IFP, and decreased in obese BMI KOA-IFPs (Fig. 8C; Supplementary Table 7B). Additionally, when comparing fibroblast cultures from obese versus normal BMI KOA-IFPs treated with TNFα, levels of 10 metabolites were significantly differentially detected (p<0.05) compared to vehicle control (Fig. 8D; Supplementary Table 7C). Mean levels of serotonin, 3-hydroxybutyric acid, 3-hydroxyisobutyric acid, glutaric acid, TGs (54:7) and (52:5), and N-acetyl-glycine were increased in TNFα-treated fibroblast cultures from obese BMI KOA-IFPs, and decreased in normal BMI KOA-IFPs (Fig. 8D; Supplementary Table 7C). In contrast, mean levels of sphingomyelins (SMs) C24:0, C16:0, and C16:1 were increased in TNFα-treated normal BMI KOA-IFP, and decreased in obese BMI KOA-IFPs (Fig. 8D; Supplementary Table 7C). Overall, this data shows differential metabolite levels in the supernatant from fibroblasts of obese compared to normal BMI KOA-IFPs under normal conditions and in response to profibrotic and proinflammatory stimuli.

## Discussion

Herein, we created a transcriptomic map of the IFP, identifying and spatially resolving major cell types and subtypes using multiple technologies including snRNA-seq and spatial transcriptomics, and advanced bioinformatic analysis. We also revealed that each cell type subcluster had a unique transcriptomic profile. Furthermore, we identified transcriptomic differences within IFP fibroblasts based upon KOA, sex, and obesity status, and metabolic alterations based on obesity status. To our knowledge, our study is the first comprehensive, tissue-specific, patient-matched transcriptomic map of the IFP, including transcriptomic analyses for KOA, sex, and obesity status, and using spatial transcriptomics to localize major cell populations, subsets, and transcriptomic profiles.

By employing CellChat, we determined putative communications between major cell types identified within the IFP. We found that CD44 and ITGAV/IGTB8 receptors on fibroblasts had the highest proportion of interactions with other major cell types. Interestingly, CD44 binds fibronectin and collagen while ITGAV/ITGB8 binds fibronectin^24,25^. We found fibronectin ligands were expressed by macrophages and endothelial cells, while collagen ligands were expressed by adipocytes and endothelial cells (Fig. 4A). This may indicate an overexpression of collagen and fibronectin as ligands from macrophages, adipocytes, and endothelial cells resulting in modification of fibroblast activity through CD44 and integrins.

After establishing transcriptomic profiles of fibroblast subclusters, we revealed transcriptomic differences based on KOA, sex, and obesity status. When comparing fibroblasts within KOA-IFPs to healthy control donor IFPs, we identified 38 DEGs. Notably, transcription factors SOX5 and CUX1 were upregulated and JUN and RBPJ were downregulated. These transcription factors have been linked to fibrosis, cell proliferation, differentiation, and ECM composition, mechanisms typically associated with KOA pathogenesis^26–29^. These alterations in transcriptomic profiles within KOA-IFP compared to healthy control donor tissues indicate possible changes in the function of fibroblasts, potentially contributing to KOA pathogenesis.

When investigating fibroblasts within female versus male KOA-IFP, we identified 105 DEGs, with transcription factors SOX5, FOXP2, and CREB5 upregulated and ZEB1, BCL6, ZBTB16, EBF2, and RBPJ downregulated. Upregulated transcription factors FOXP2 and SOX5 have been linked to fibrosis and cartilage degeneration, suggesting more prominent presentation of these pathologies within female KOA- IFP compared to males^30,31^. Furthermore, CREB5 impacts the yield of lubricin, important for joint lubrication and encoded by PRG4, which was also upregulated within female KOA-IFP^32^ (Fig. 6C). This suggests that fibroblasts from female IFPs may contribute more to joint lubrication as compared to fibroblasts from male IFP. Additionally, all downregulated transcription factors can be linked to inflammation, indicating alterations in pro-inflammatory responses within female IFPs^33–36^. Together, differences in the function of fibroblasts within female compared to male KOA-IFPs may modify disease pathology between sexes.

Within the fibroblasts of obese compared to normal BMI KOA-IFPs, we identified 21 DEGs. Specifically, the transcription factor PRRX1 was upregulated while ZBTB16 and CREB5 were downregulated. PRRX1 positively regulates expression of collagens^37^. Together, upregulation of PRRX1 and downregulation of ZBTB16 (as described above) may indicate increased fibrosis- and inflammation- associated signaling in obese versus normal BMI KOA-IFP, both characteristic of OA pathology. In contrast to female KOA-IFPs, CREB5 and PRG4 were downregulated within obese compared to normal BMI KOA- IFPs, suggesting lubricin production may be reduced in relation to obesity status. Overall, these observations suggest fibroblast activities are likely different in obese versus normal BMI KOA-IFPs and may contribute to accelerated KOA progression in obese BMI individuals, which requires further study.

While investigating the DEG profile of fibroblasts from obese compared to normal BMI KOA-IFP, pathways including metabolism of carbohydrates pathway, carbohydrate binding, and multiple lipid transport and ABC-family-related pathways (related to active transport of small molecules)^38^ were enriched (Fig. 7G). As metabolic dysfunction is associated with obesity and KOA, we conducted metabolomic analysis to parse out differences in metabolite levels in supernatant of IFP fibroblasts based on obesity status. We identified 5 metabolites differentially secreted from fibroblasts derived from obese versus normal BMI KOA-IFPs. Metabolites C18:2 (linoleic acid), TGs (54:2) and (50:2), and choline were upregulated while homoarginine was downregulated. Linoleic acid and choline can modify pain responses while homoarginine is an anti-inflammatory mediator^39–41^. The exact contribution of metabolite changes in obese versus normal BMI KOA-IFP fibroblasts and possible relationships with pain and inflammatory mechanisms during OA should be further explored.

We also found that the response of KOA-IFP fibroblasts to profibrotic and proinflammatory stimuli were altered based on obesity status. When KOA-IFP fibroblasts were stimulated with profibrotic TGFβ, homocitrulline, C16:1OH, and Hex2Cer 34:1 were increased while SMs C24:0, C16:0, and C16:1 were decreased in supernatants of cultures from obese versus normal BMI KOA-IFP. Homocitrulline and Hex2Cer 34:1 are associated with increased inflammation, while Hex2Cer 34:1 is linked to increased fibrosis^42,43^. C16:1OH is associated with increased oxidative stress^44^. When stimulated with proinflammatory TNFα, fibroblast cultures from obese BMI KOA-IFP patients had increased supernatant metabolite levels of serotonin, 3-hydroxybutyric acid, 3-hydroxyisobutyric acid, glutaric acid, and n-acetyl-glycine compared to cultures from normal BMI KOA-IFP. Serotonin aids in regulating inflammatory and fibrotic responses^45^. 3- Hydroxybutyric acid reduces the expression of pro-inflammatory enzymes and cytokines^46^. Furthermore, 3-hydroxyisobutyric acid, glutaric acid, and n-acetyl-glycine play roles in fatty acid metabolism, with increased levels leading to deregulation^47–49^. In contrast, TNFα stimulation decreased levels of SMs C24:0, C16:0, and C16:1, precursors for ceramide, which impacts inflammation^50^. Overall, metabolite level changes in response to profibrotic and proinflammatory stimuli indicate that fibroblasts from obese BMI KOA-IFPs may promote inflammation and fibrosis, and have deregulated fatty acid metabolism, as compared to fibroblasts from normal BMI KOA-IFP. Further investigation to identify differences in the functions of fibroblasts, and other major KOA-IFP cell types, in KOA pathogenesis based on obesity status should be considered.

This study has utilized snRNA-seq, spatial transcriptomics, and advanced bioinformatic approaches to generate comprehensive transcriptomic and spatial profiles of the human IFP. By applying these high throughput methods, we identified unique transcriptomic profiles of major cell populations and subtypes present within IFP, including fibroblasts, macrophages, adipocytes, and endothelial cells. Furthermore, we identified transcriptomic differences within fibroblasts of the IFP based on OA, sex, and obesity status, and metabolic alterations based on obesity status using metabolomics. Overall, this study provides a comprehensive map of the cellular and transcriptomic diversity of the human IFP using a multi-omic approach.

## Acknowledgements

This work is supported by the Arthritis Society of Canada Strategic Operating Grant (23-0000000271), Canada Research Chairs Program, Tony and Shari Fell Platinum Chair in Arthritis Research (University Health Network Foundation, University Health Network, Toronto. IJ was supported in part by funding from Natural Sciences Research Council (NSERC #203475), Canada Foundation for Innovation (CFI #225404, #30865), and Ontario Research Fund (RDI #34876, RE010-020). Thank you to the Schroeder Arthritis Institute Orthopaedic Research Team for consenting and collecting study participant data. We thank Edwin Speck from Princess Margaret Flow Cytometry (https://pmflow.ca), University Health Network for their assistance in FACS cell sorting. Thank you to Melanie Peralta from the Pathology Research Program within University Health Network’s Laboratory Medicine Program – Pathology Department for their assistance with the OCT- embedded samples. We would also like to thank Farzaneh Aboualizadeh from the Princess Margaret Genomics Center, University Health Network, for their assistance with spatial sequencing within this study.

## Author Contributions

HP contributed to study design, selection study participants for IFP samples, acquisition of data, interpreting all analyses, fibroblast culture experiments, and writing of the manuscript and generation of figures. PP contributed to the analysis and interpretation of single-nucleus RNA sequencing data, CellChat analysis, and generation of associated figures. JSR contributed to the analysis and interpretation of all experiments and the acquisition of funding for this study. TT contributed to the analysis and interpretation of spatial sequencing data, and generation of associated figures. CP and IJ contributed to all GO biological process and pathDIP analysis and generation of associated figures. KDS contributed to tissue preparations and the completion of single-nuclei RNA sequencing experiments and data analysis. SV has contributed to the analysis pipeline used for single-nucleus RNA sequencing data, the completion of trajectory analysis and generation of associated figures. SL contributed to study design and the experimental pipeline used for single-nucleus RNA sequencing. KP contributed to the acquisition and storage of all KOA-IFP samples and clinical data through the Schroeder Arthritis Institute Orthopaedics Biobank. NL, SHL, and VC contributed to the completion of all targeted metabolomics experiments. KH contributed to the analysis and interpretation of all targeted metabolomics experiments and generation of associated figures. PK contributed to fibroblast culture experiments. AVP and YRR contributed to collection and interpretation of data from LEAP OA clinical database. NNM and KS contributed to the collection of KOA-IFP samples and data collection. EG contributed to data interpretation. RK contributed to the collection of healthy control donor IFP samples and data analysis. MBB contributed to the study planning and interpretation of cell population mapping using previously published literature. RG contributed to the conception and acquisition of funding for this study, the acquisition of KOA-IFP, and interpretation of the data. MK contributed to the conception and acquisition of funding for this study, and the interpretation of all experiments. All authors have contributed significantly to the critical revision of the manuscript.

## Supplementary Materials and Methods

### Sample collection

Infrapatellar fat pad (IFP) tissues were obtained from study participants with knee OA undergoing total knee arthroplasty and recruited within the LEAP-OA cohort (Division of Orthopaedics, Schroeder Arthritis Institute, University Health Network, Toronto). Surgeon-identified IFPs were collected consistently from the retropatellar tendon region from discarded tissues during total knee arthroplasty, under informed consent and under research ethics board approval (REB #07-0383, #14-7592). Healthy control donor IFP biospecimens (obtained <4 hours post-mortem) from individuals without known musculoskeletal disease were obtained in collaboration with Dr. Roman Krawetz at the University of Calgary (REB #21987). Both KOA (n=15) and healthy control donor (n=6) IFP were flash frozen and stored in liquid nitrogen at -190°C in a temperature monitored vapour liquid nitrogen tank until use. Within this study, normal BMI study participants were classified as 18.5 to 24.9 kg/m^2^, while obese BMI study participants were 30 to 40 kg/m^2^. In total, n=21 IFPs were analyzed as detailed in Supplementary Table 1. Histology of KOA IFPs can be found in Supplementary Fig. 1.

For fibroblast culture studies, fresh KOA-IFPs (n=5 obese BMI, n=5 normal BMI) were obtained from subjects undergoing total knee arthroplasty and tissues were further digested and processed for cell culture (REB #14-7592).

### Single nucleus RNA sequencing (snRNA-seq)

Nuclei were isolated from frozen human IFPs (n=21, weighing approximately 50 mg) by physical dissociation using a glass dounce homogenizer followed by sorting using fluorescence-activated nuclei sorting (FANS) based on DAPI-positive fluorescence with support from Krembil Discovery Tower Flow Cytometry Facility (University Health Network) (protocol adapted from Mazutis et al., 2020^1^). Nuclei concentration was determined by DAPI staining and hemocytometer counting, with manual adjustment of the sample to 1000 nuclei/μL. SnRNA-seq libraries were generated using the 10X Genomics Chromium Next GEM Single Cell 3’ Reagent Kits (v3.1 Dual Index). Sequencing was performed on the Illumina NextSeq 550 system (2x150 bp High Output Kit) at the Arthritis Program Diagnostic and Therapeutic Innovation Centre (Schroeder Arthritis Institute).

A feature and UMI count matrix for each sample was generated from 10X Genomics 3’ Gene expression sequenced reads using CellRanger software (v7.0.0)^2^. From these files, Seurat objects were created and loaded into R (v4.3.1), where filtering of nuclei was performed based on percentage of mitochondrial content (<5%)^3^. DoubletFinder (v2.0.3) R package was used to remove doublets^4^.

Integration of all 21 samples was performed using Harmony (v0.1) R package^5^, and standard processing workflow steps were followed as follows: data were log-normalized, scaled after identifying highly variable features (n=2000), followed by PCA analysis (npcs=30) and graph-based clustering (resolution=0.5) yielding a single object of 73,808 nuclei with 31151 features. The Seurat (v4.4.0) R package was employed for the above steps to generate Uniform Manifold Approximation and Projection (UMAP) plots, where nuclei were clustered together based on similarity^3^. Each cluster identified was annotated using canonical markers (Fig. 1B).

The top four major cell types identified (> 10% of nuclei by proportion) were further explored by performing subcluster analysis individually for each cell type. Cell subsets within the major cell type populations were established based on similar highly expressed genes. For each cell type subclustering, only nuclei respective to that cell type were subsetted from the main Seurat object, followed by standard workflow steps mentioned above. Nuclei were re-integrated using Harmony R package, where the dimensionality was determined using “ElbowPlot” function (npcs=30) and optimal clustering resolution was explored using Clustree (v0.5.1) R package for each cell type^6^.

When comparing fibroblast proportions by OA, sex, or obesity status, fibroblast subcluster proportions were analyzed by arcsin-transforming absolute proportions across all samples and performing multiple unpaired t-tests with FDR correction using the Benjamini, Krieger and Yekutieli two stage step up method using GraphPad Prism v10.1.1 (GraphPad Software, Boston, Massachusetts USA, www.graphpad.com). Differences within proportions of fibroblast subclusters based on OA, sex, or obesity with adjusted p<0.05 were considered significant.

### Differentially expressed gene (DEG) signatures determination

DEGs within fibroblasts, macrophages, adipocytes, and endothelial cell subclusters were identified using the Seurat (v4.4.0) R package^3^. DEGs were filtered based on the following criteria: Log2FC > 0.5, adjusted p<0.05, calculated based on Bonferroni correction, a minimum of 25% of nuclei expressing each gene in each subcluster. From the resulting lists of genes, only known coding genes were retained, with all non-coding genes (e.g. long non-coding genes, miRNAs), mitochondrial genes, and unknown genes removed. Duplicate genes present in more than one subcluster were also filtered, generating a list of unique DEGs within each cell subcluster.

To generate a list of differentially expressed genes within fibroblasts from KOA-IFP compared to healthy control donor IFP, from female compared to male KOA-IFP, as well from obese BMI compared to normal BMI KOA-IFP, a similar process was followed. DEGs between above groups of samples were determined using Seurat (v4.4.0) and genes that had an adjusted p<0.05 and minimum 50% of nuclei expressing each gene were retained. All non-coding, mitochondrial, and unknown genes were removed, as described above.

### Trajectory analysis

Pseudotime trajectory analysis was performed using the R package SCORPIUS (v1.0.9)^7^. Trajectory inference was conducted on 2D embeddings using the “infer_trajectory” function after reducing the dimensionality of the dataset. SCORPIUS’s “gene_importances” function was then used to determine the top genes involved in ordering of the nuclei along pseudotime. Candidate marker genes with adjusted p<0.05 and scaled expression were grouped into modules. The expression of these grouped genes was visualized using a heatmap.

### Comparison of cell subclusters with published datasets

To compare similarities of identified cell subclusters within fibroblasts, macrophages, adipocytes, and endothelial cells, gene module scores were generated using “AddModuleScore” function of the Seurat (v4.4.0) R package^3^ based upon transcriptomic profiles of previously published datasets analyzing white adipose tissues. All gene module scores were plotted as violin plots and feature plots for visualization (refer to Supplementary Fig. 10). The study by Emont et al.^8^ had extensive and clear representations of identified subclusters for fibroblasts, macrophages, adipocytes, and endothelial cells within white adipose tissue. Gene module scores were generated based on top ten DEGs for each respective cell type mentioned in Supplementary Table 1 of Emont et al.^8^, after excluding non-coding genes. From Merrick et al.^9^, markers mentioned in Figure S13 from the publication were used to generate module scores. Within the Hepler et al.^10^ publication, Figure 1D displayed a heatmap of the top 20 DEGs across their fibroblast subtypes, for which gene module scores were calculated. The publication from Buechler et al.^11^ presented 20 genes as universal fibroblast markers and were used to generate module scores. Finally, using the publication from Tang et al.^12^, markers used to identify fibroblast subclusters were located within the dot plot of Figure 2B and compared to fibroblast subclusters identified in our study. Gene module scores from each independent publication were then used to compare to cell type subclusters identified in our study.

### Spatial transcriptomics

Fresh human KOA-IFPs (n=12) were embedded in Optimal Cutting Temperature (OCT) medium (GeneralData Cat# TFM-5) and frozen in liquid nitrogen. Frozen tissue blocks were sectioned (10 μM) on a cryostat (Leica CM1950 Cryostat) and mounted onto VWR Superfrost glass slides (48311-703). RNA was isolated using the Qiagen Rneasy Mini kit (Cat#74104) following manufacturer’s recommendations. RNA quality control was performed with the Agilent Bioanalyzer using the RNA 6000 Nano Kit (Cat# NC1783726). See RIN and DV200 of all the samples used for spatial sequencing in Supplementary Table 8. Tissues were fixed with methanol and Hematoxylin & Eosin (H&E) stained sections were prepared following manufacturer’s instructions (10x Genomics, Methanol Fixation, H&E Staining & Imaging for Visium Spatial Protocols, CG000160 Rev B). Tissues were imaged with Leica Dmi8 Inverted (color camera) in brightfield with 20X magnification. Sequencing libraries were constructed using 10X Genomics Visium CytAssist Spatial Gene Expression Reagent Kits (Cat# CG000495). Library quality was assessed on a high sensitivity DNA chip using an Agilent Bioanalyzer (5067-4626). Sequencing libraries were volumetrically pooled and sequenced on an Illumina NovaSeq sequencer for 150 paired end read-cycles at the Princess Margaret Genomics Cancer Centre (Toronto, ON, Canada).

Spatial transcriptomic data from n=12 KOA-IFP (normal BMI n=6, obese BMI n=6) were analyzed using Scater v1.28.0^13^ and Seurat v4.3.0^3^ for quality control and filtering, and Seurat v5.0.1^14^ for downstream processes. Scater removed barcodes with high mitochondrial content by calculating the median-absolute deviation. Further, barcodes were removed that expressed high Unique Molecular Identifiers per barcode. Filtered data was subject to normalization using “NormalizeData”, and integration using Harmony v0.1.1^5^. Clustering was accomplished using a resolution of 1.9 and 30 PCs, yielding 31 communities. Clustree R package v0.5.1^6^ was consulted to select a clustering resolution. Canonical markers were identified using the “FindAllMarkers” function, with min.pct set to 0.25 and only returning positive markers. Clusters were manually annotated by searching against a custom list of markers for cell types of interest (Fig. 1B) combined with markers from the PanglaoDB database^15^. The annotated fibroblast population was extracted from the spatial dataset and reanalyzed with the same workflow as stated above, setting the clustering resolution to 0.2. Samples were labeled according to obesity status, and differential expression analysis was accomplished using the “FindMarkers” function. Lastly, deconvolution of fibroblasts was completed using the Seurat deconvolution workflow using default parameters, except for setting the resolution parameter to “CCA” for function “FindTransferAnchors”. The query dataset was set to the fibroblast spatial population, and the reference dataset was set to the fibroblast snRNAseq population.

### Cell-cell communication analyses

To explore ligand-receptor interactions across four major cell types in our snRNA-seq data, we used CellChat v2.0 R package^16^, where fibroblasts were chosen as the “receiver” cell type and adipocytes, macrophages, and endothelial cells as the “sender” cell type. The “netVisual_chord_gene” function in CellChat was used to create a chord diagram to visualize the complex ligand-receptor interactions between source and target cell types. To identify and visualize the outgoing communication patterns of each cell type of our interest, we used the “netAnalysis_river” function to generate a river (alluvial) plot showing how each cell type coordinates to drive communication across multiple genes and signaling pathways.

### Pathway analyses

Up and downregulated DEG-set lists were used separately to perform pathway and gene ontology enrichment analysis. Pathway enrichment analysis was performed using pathDIP v5 (https://ophid.utoronto.ca/pathDIP)^17^ API in R 4.3.0, excluding Disease and Drugs & Vitamins pathways, and using literature curated associations. Statistically significant pathways (adjusted p < 0.05) were further considered. Gene Ontology Biological Processes enrichment analysis was performed using clusterProfiler v4.8.3^18^ in R, retaining terms with adjusted p < 0.05. Catrin database v.1 (unpublished; https://ophid.utoronto.ca/Catrin) was used to identify transcription factors (TF) targeting and part of the deregulated genes. A network of protein-protein interactions and term – gene – TF interactions was built using NAViGaTOR v3.0.16^19^. Interactions among genes were retrieved using IID v. 2020-11^20^. All other figures were plotted using ggplot2 3.4.2 in R^21^.

### Fibroblast cell culture

Fresh KOA-IFPs (n=5 obese BMI KOA-IFP, n=5 normal BMI KOA-IFP) were gathered from discarded tissues after total knee replacement and immediately mechanically digested with scissors. Tissues were enzymatically digested using trypsin (1mg/mL) (Sigma) for 30 minutes and collagenase type 1 (2mg/mL) (Sigma) for 3 hours, each diluted in DMEM (Gibco)+1%PenStrep (Wisent). The fatty layer was removed, and the cell pellet was washed twice and resuspended in DMEM+1%PenStrep+10%FBS (Gibco). Cell suspensions were plated into 20mL cell culture flask and incubated at 37°C until 80% confluent. Fibroblasts were then lifted from the cell culture flask using trypsin (0.05%) (ThermoFisher) and frozen at passage 0 in FBS+10%DMSO (Fisher Scientific) with 1 million cells per cryovial. Frozen fibroblasts were stored in liquid nitrogen until use.

Frozen fibroblasts were gently thawed within a 37°C water bath for 10 seconds and placed into warmed DMEM+1%PenStrep+10%FBS. Thawed fibroblasts were plated into a 20mL cell culture flask and incubated at 37°C for 48 hours until 100% confluent. Confluent fibroblasts were then lifted from the cell culture flask using trypsin (0.05%) (ThermoFisher), passaged to a 12 well plate with 80,000 cells per well with DMEM+1%PenStrep+10%FBS, and incubated at 37°C for 24 hours. At 100% confluency, fibroblasts were then starved for 24 hours with DMEM+1% PenStrep+1%FBS. After starvation, fibroblasts were treated with either Control (DMEM+1%PenStrep+1%FBS), Control+Vehicle [DMEM+1%PenStrep+1%FBS+1%PBS (Gibco)], TGFβ [DMEM+1% PenStrep+1%FBS+TGFβ(20ng/mL, Biotechne)], or TNFα [DMEM+1% PenStrep+1%FBS+TNFα(10ng/mL, Biotechne)], and incubated for 48 hours. Supernatant was collected, frozen and stored at -80°C until use.

### Metabolomics

Samples were processed at the Schroeder Arthritis Institute’s Metabolomics Facility. A targeted quantitative metabolomics approach was applied to analyze supernatant samples using a combination of direct injection (DI) mass spectrometry (MS) with a reverse-phase liquid chromatography (LC)-MS/MS using a custom MTX MEGA Assay (MetabolomiX, Edmonton, Canada). This custom assay, in combination with a QE orbitrap (ThermoFisher Scientific, Waltham, MA) mass spectrometer, was used for targeted identification and quantification of 646 endogenous metabolites including amino acids or amino acid-related metabolites, biogenic amines, ceramides, cholesterol esters, diacylglycerols, acylcarnitines, glycerophospholipids, sphingomyelins, triacylglycerols, organic acids and nucleotide/nucleosides. The method combines the derivatization and extraction of analytes, and the selective MS detection using multiple reaction monitoring (MRM) pairs. Isotope-labeled internal standards and other internal standards are used for metabolite quantification. For all metabolites except organic acids, samples were thawed on ice and vortexed and centrifuged at 13,000 x g. Samples were loaded onto a filter spot within an upper chamber of a 96-well plate and dried in a stream of nitrogen. Subsequently, phenyl- isothiocyanate was added for derivatization. After a 20 min incubation, filter spots were dried again using an evaporator. Extraction of the metabolites was achieved by adding 300 µL of extraction solvent. The extracts were obtained by centrifugation into the lower 96-deep well plate, followed by a dilution step with MS running solvent. For organic acid analysis, 150 µL of ice-cold methanol and 10 µL of isotope-labeled internal standard mixture was added to samples for overnight protein precipitation, followed by centrifugation at 13000x g for 20 minutes. Next, 50 µL of supernatant was loaded into the center of a 96- deep well plate, followed by the addition of 3-nitrophenylhydrazine (NPH) reagent. After incubation for 2 hours, butylated hydroxytoluene stabilizer (BHT) and water were added before LC-MS injection. MS analysis was performed on a QE Orbitrap spectrometry instrument equipped with a Vanquish Flex UHPLC system (ThermoFisher Scientific, Waltham, MA). The samples were delivered to the MS by a LC method followed by a DI method.

Metaboanalyst (v6.0)^22^ was used to process and analyze metabolomic data, using one-factor and metadata statistical analysis modules. All metabolites with missing values, those with values below the level of detection, or zero, were replaced with 1/10^th^ of the lowest detected concentration for that metabolite. All metabolites that had zero variance across all samples were removed. The metabolites hexose, C0, C12, C2 and C14:1 were also removed due to unreliable high values or extreme outliers impacting normalization. Samples were then normalized by sum and each metabolite was log_10_ transformed. Metabolites were then scaled using the Pareto approach. To compare metabolite levels in supernatant from fibroblast cultures from obese compared to normal BMI KOA-IFP, FCs and p-values were calculated using unpaired equal variance t-tests.

We then analyzed the interaction between obesity status groups (obese vs normal BMI KOA-IFP) and treatment (TGFβ or TNFα vs Control + Vehicle), to test whether response to treatment differed in the supernatants of obese compared to normal BMI KOA-IFP fibroblast cultures. Interaction analysis was conducted in Metaboanalyst using the “limma” R package^23^; a linear model was fit regressing metabolite expression on study subject, treatment, and the interaction between treatment and obesity status. Significant p-values (p<0.05) on the interaction term, or “change in log2FC”, indicate that the change in metabolite expression from control-vehicle to treatment was different within the obese BMI KOA-IFP group compared to the normal BMI KOA-IFP group. A positive change in log2FC indicates that the metabolite level within the obese BMI KOA-IFP group increased more than within the normal BMI knee-OA IFP group when treated.

### Data Availability

Raw sequencing and processed data were submitted to GEO data repository with a Superseries GEO accession ID: GSE253200. There are two sub-series where GEO accession ID: GSE253198 consists of 21 single nucleus RNA-seq IFP samples and GEO accession ID: GSE253199 consists of 12 spatial transcriptomics IFP samples. Note that all 12 spatial transcriptomics IFP are patient-matched (refer to Supplementary Table 1).

### Patient and Public Involvement Statement

Patients or the public were not involved in the planning, analysis or completion of this study.

---

**Supplementary Figure 1.**
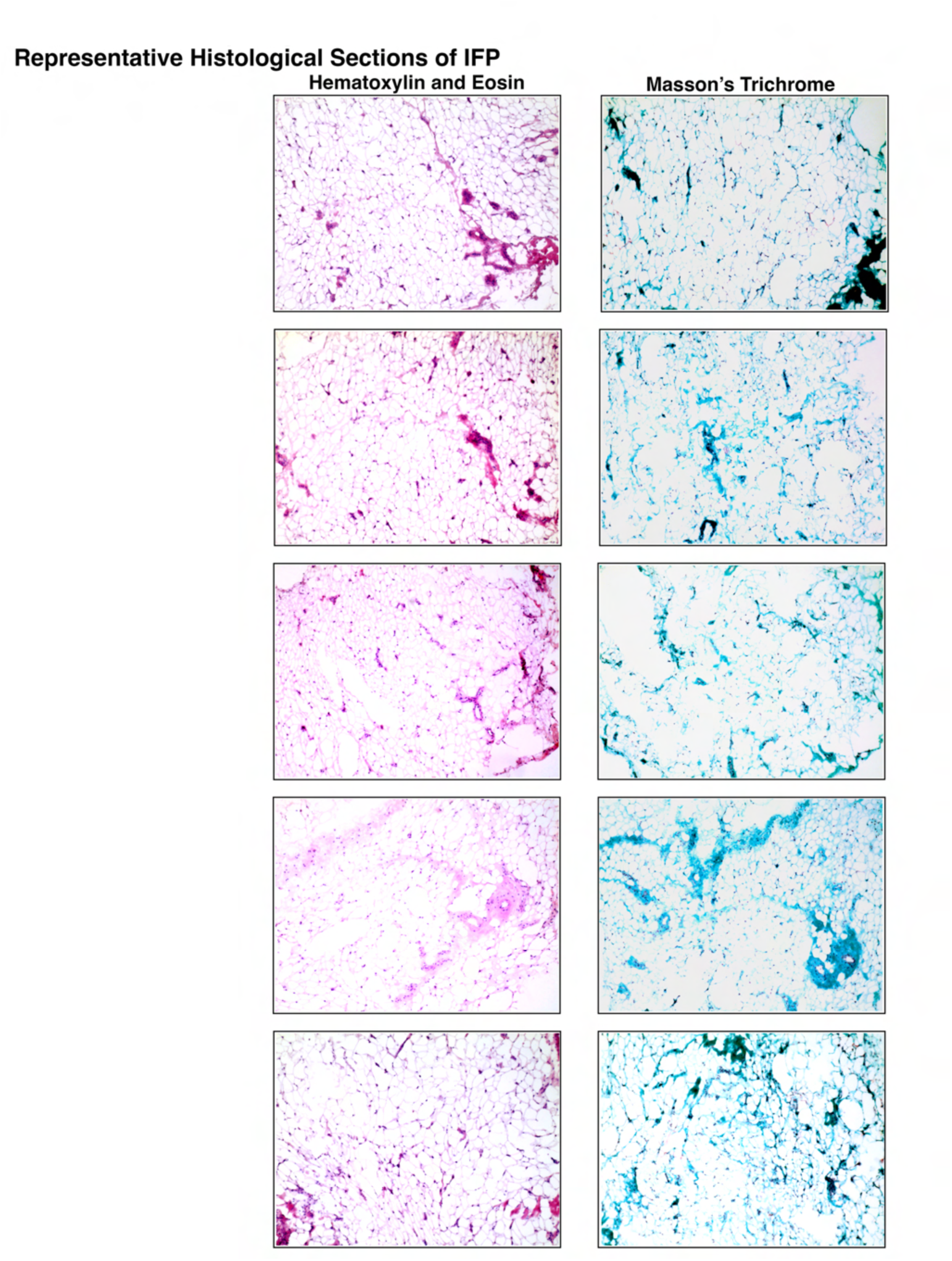
Infrapatellar Fat Pad (IFP) Histology. Representative histological sections of the knee OA IFP from five separate knee OA study participants stained with Hematoxylin & Eosin (left) and Masson’s Trichrome (right) stains.

**Supplementary Figure 2.**
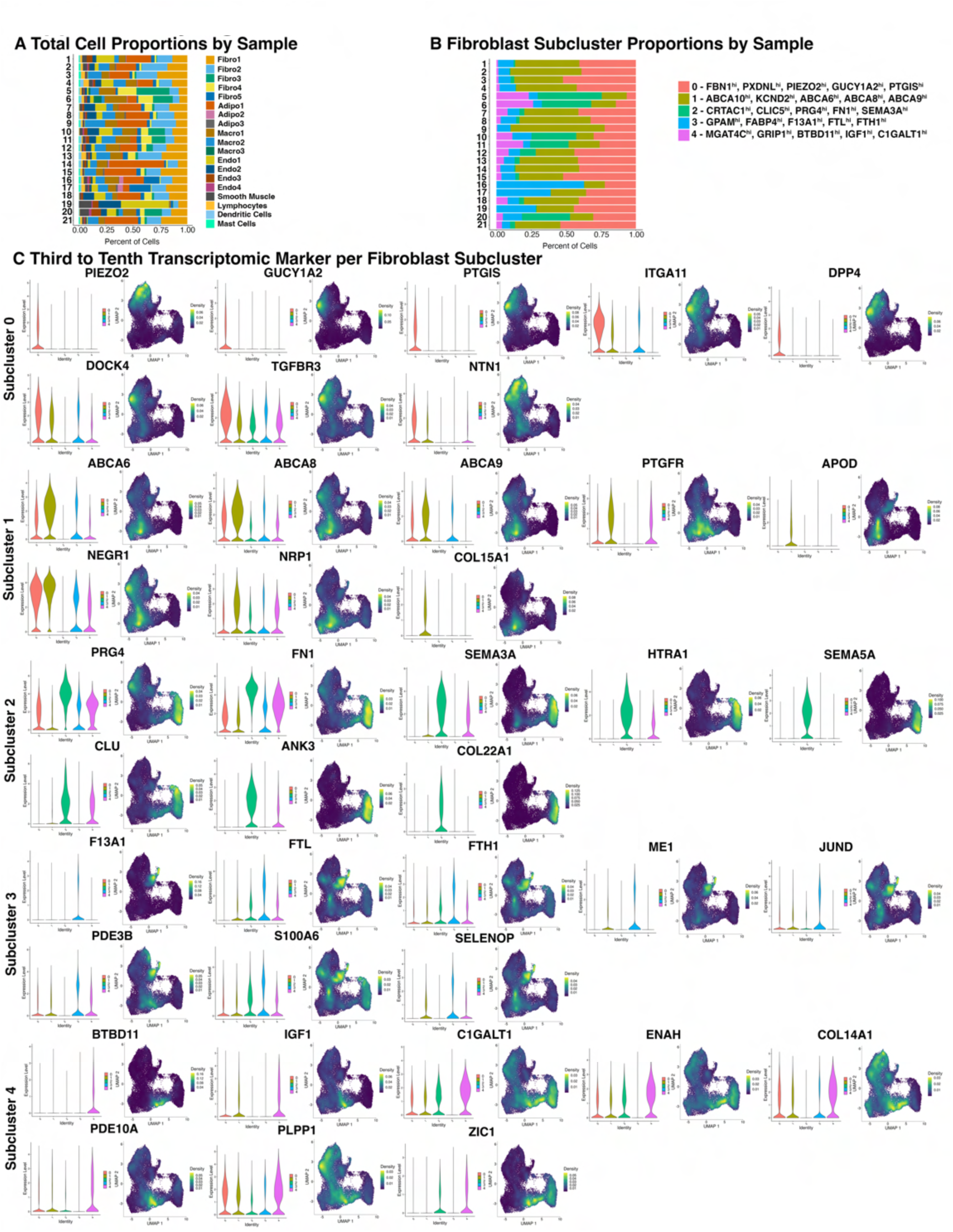
Fibroblast Subcluster Proportions and Differentially Expressed Genes (DEGs). A) Stacked bar graph showing the proportion of each cell type identified from snRNA seq analysis for n=21 IFPs, split by sample. B) Stacked bar graph showing the proportion of each fibroblast subcluster for n=21 IFP samples. Top 5 unique DEGs defining each subcluster are indicated. C) Violin and nebulosa plots showing the expression of the third to tenth top unique DEGs within each fibroblast subcluster. Nebulosa plots demonstrate expression density from dark purple (low) to yellow (high). *Also see Fig. 1, Supplementary Fig. 3, and Supplementary Table 2A*.

**Supplementary Figure 3.**
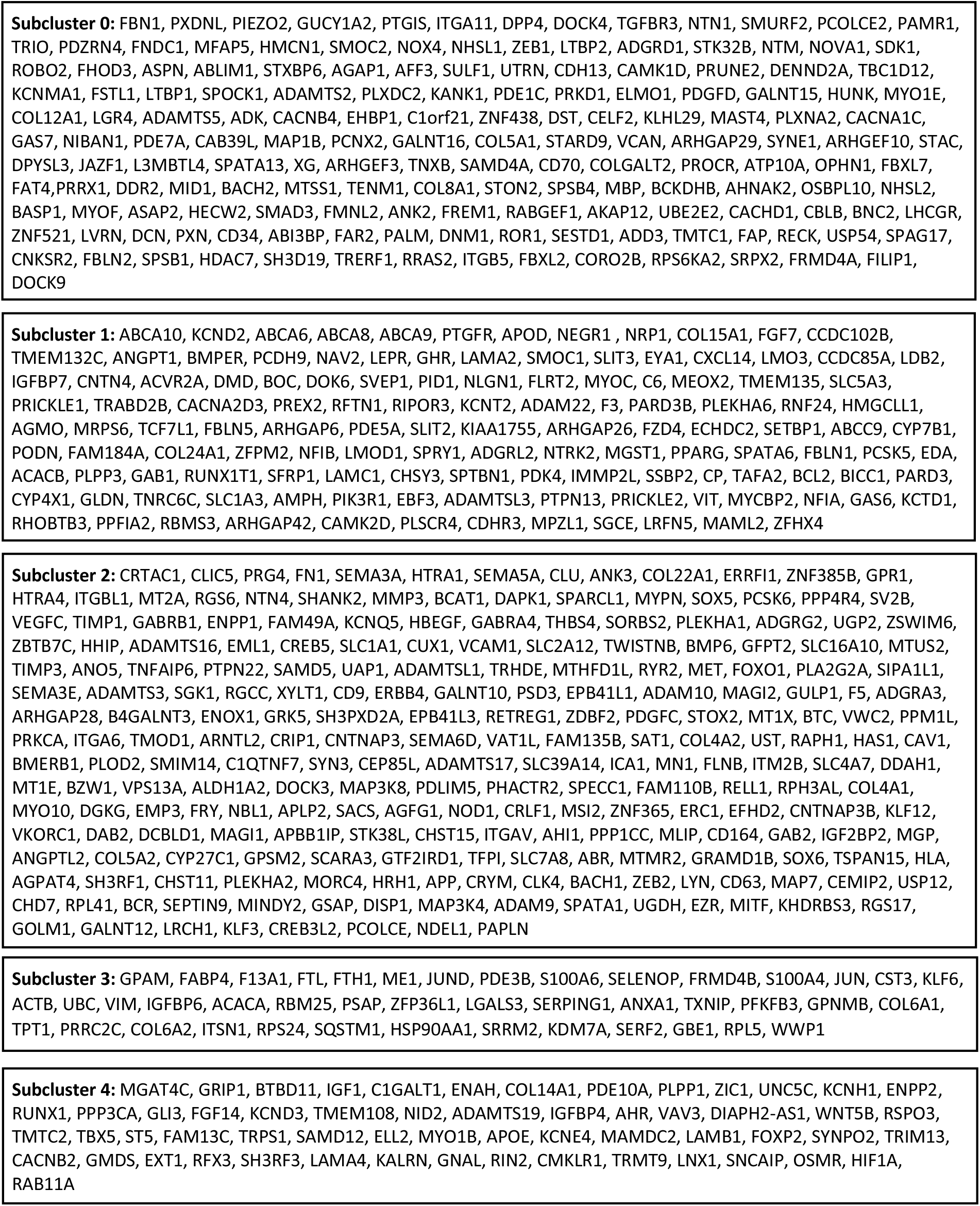
List of 613 Unique Differentially Expressed Genes Within Each Subcluster of Fibroblasts Identified by Bioinformatic Analysis of n=21 Infrapatellar Fat Pad Samples from snRNA-seq. Genes are ordered by decreasing average log2 fold change. Unknown and non-coding genes, and genes that were duplicated across subclusters, were removed. *Also see Fig. 1, Supplementary Fig. 2, and Supplementary Table 2A*.

**Supplementary Figure 4.**
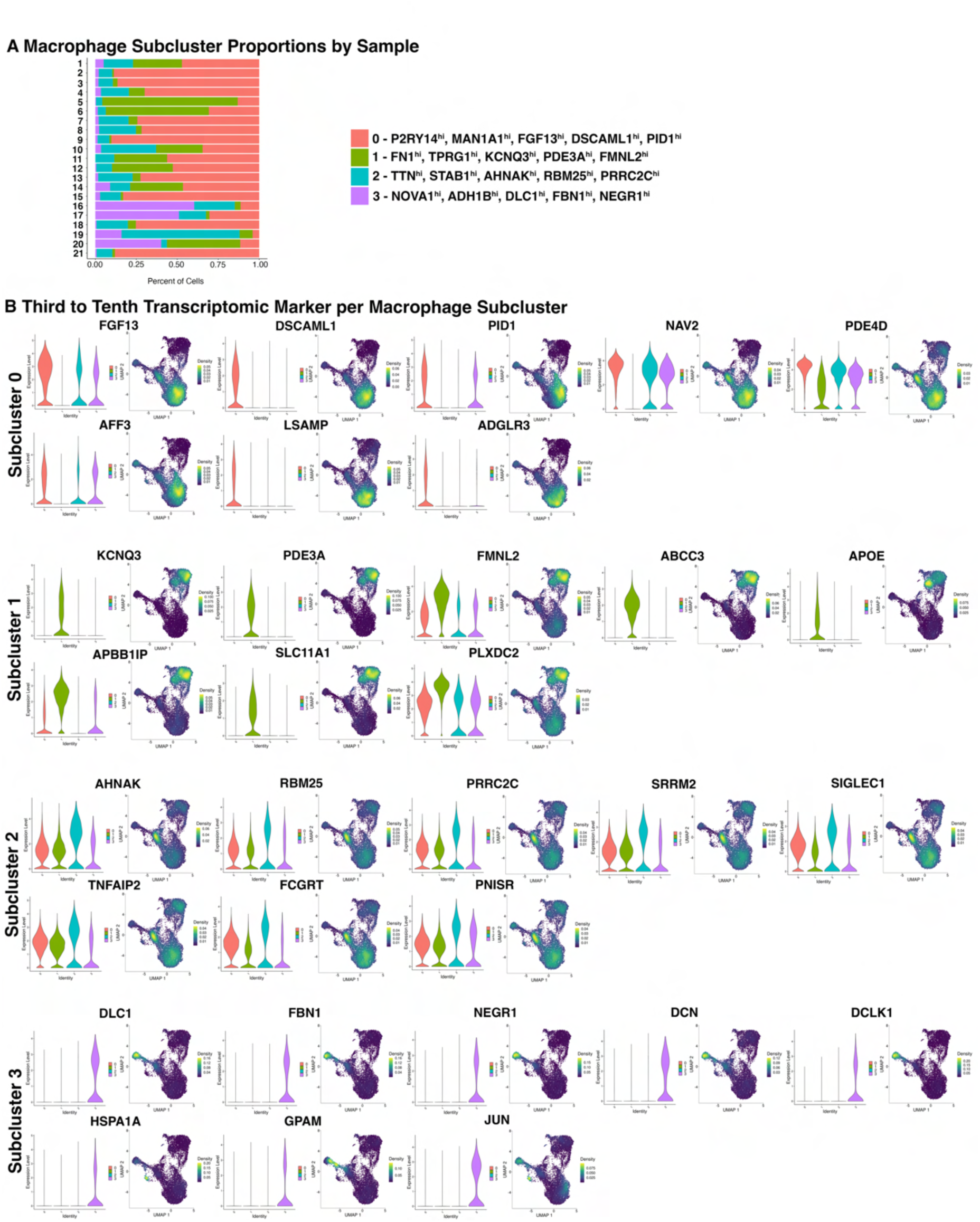
Macrophage Subcluster Proportions and Differentially Expressed Genes (DEGs). A) Stacked bar plot showing the proportion of each macrophage subcluster identified from snRNA seq analysis for n=21 IFP samples. Top 5 unique DEGs defining each subcluster are indicated. B) Violin plots and nebulosa plots showing the expression of the third to tenth top unique DEGs within each macrophage subcluster. Nebulosa plots demonstrate expression density from dark purple (low) to yellow (high). *Also see Fig. 2, Supplementary Fig. 5, and Supplementary Table 2B*.

**Supplementary Figure 5.**
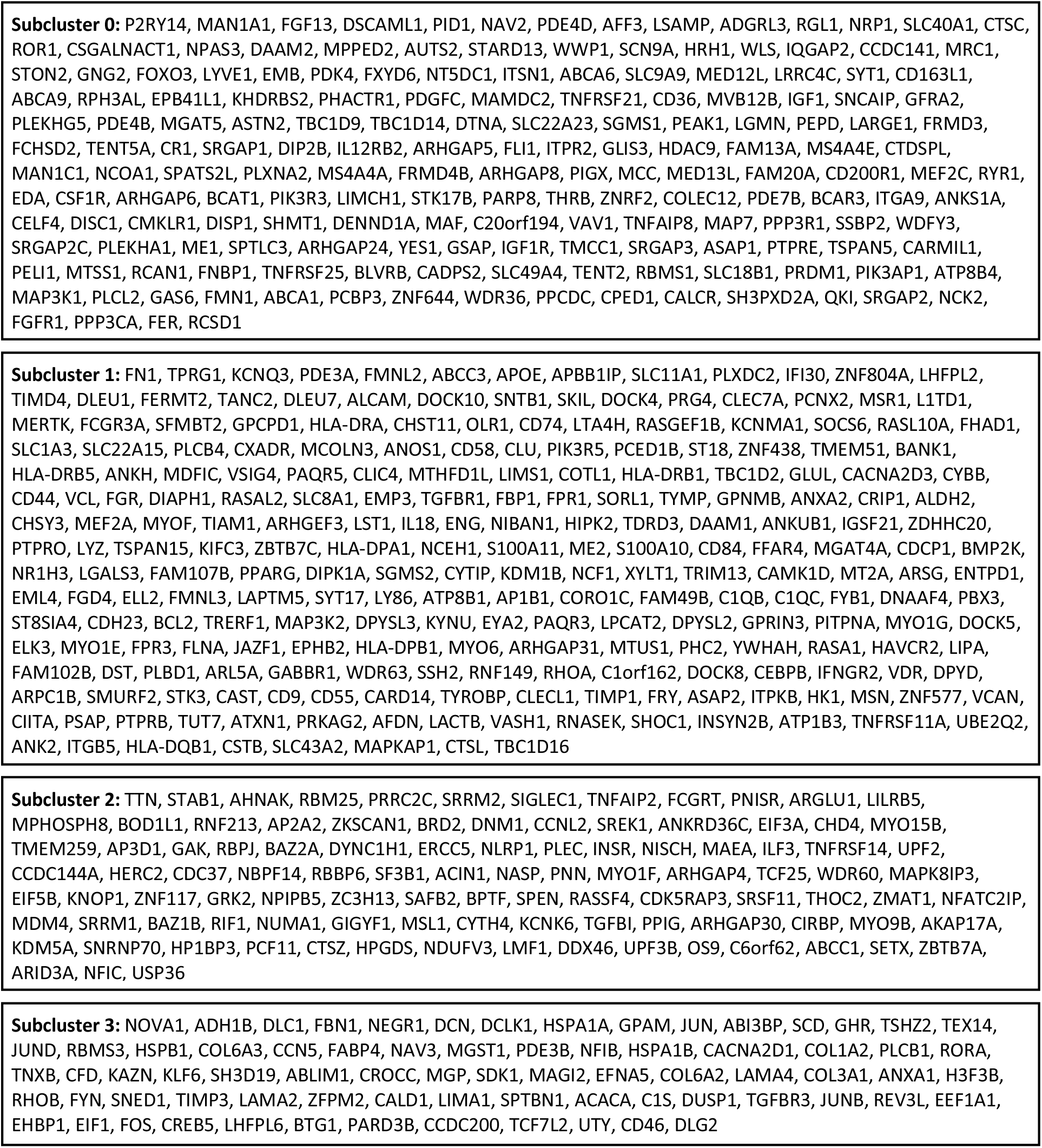
List of 582 Unique Differentially Expressed Genes Within Each Subcluster of Macrophages Identified by Bioinformatic Analysis of n=21 Infrapatellar Fat Pad Samples from snRNA-seq. Genes are ordered based on decreasing average log2 fold change. Unknown and non-coding genes, and genes that were duplicated across subclusters, were removed. *Also see Fig. 2, Supplementary Fig. 3, and Supplementary Table 2B*.

**Supplementary Figure 6.**
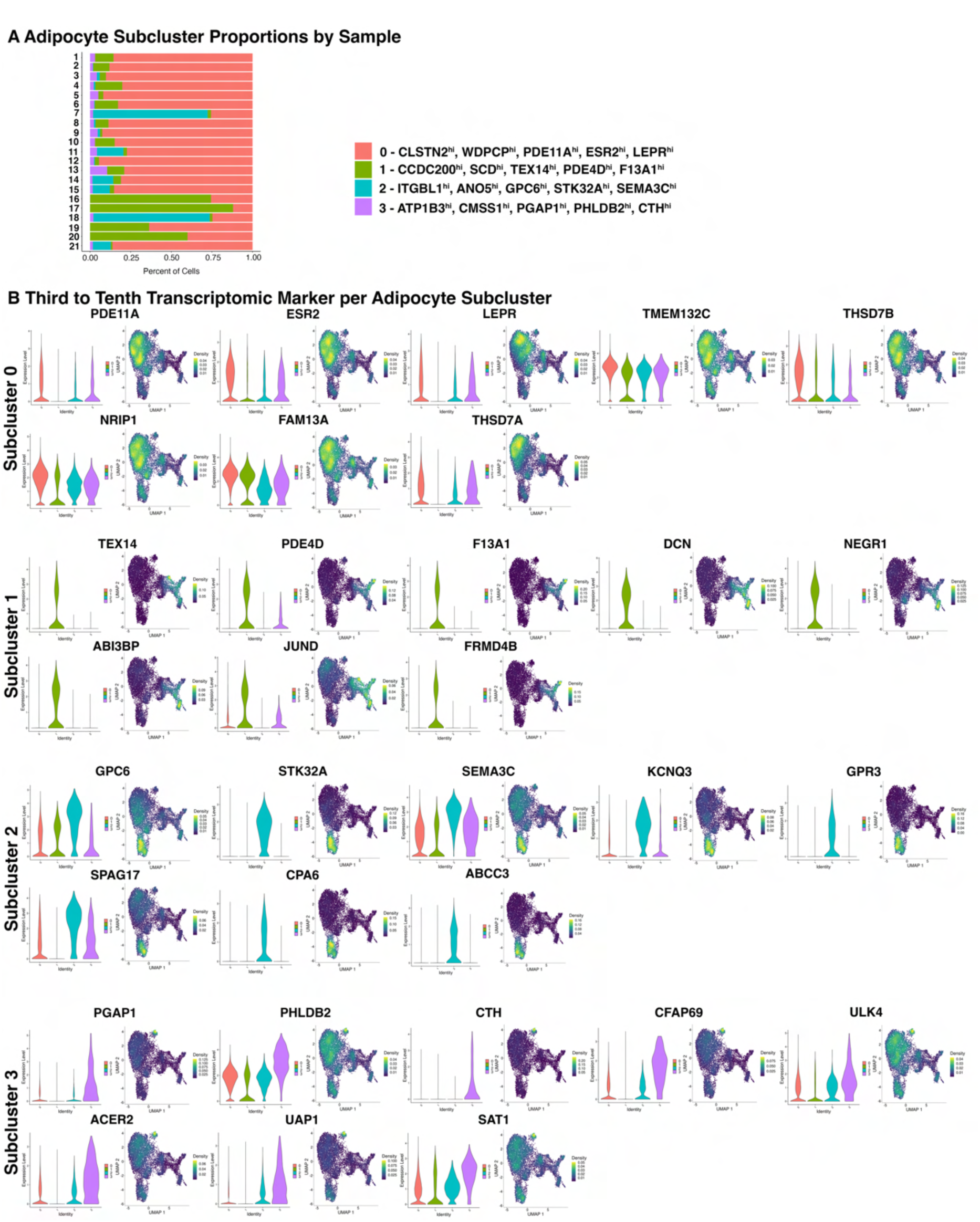
Adipocyte Subcluster Proportions and Differentially Expressed Genes (DEGs). A) Stacked bar plot showing the proportion of each adipocyte subcluster identified from snRNA seq analysis for n=21 IFP samples. Top 5 unique DEGs defining each subcluster are indicated. B) Violin plots and nebulosa plots showing the expression of the third to tenth top unique DEGs within each adipocyte subcluster. Nebulosa plots demonstrate expression density from dark purple (low) to yellow (high). *Also see Fig. 2, Supplementary Fig. 7, and Supplementary Table 2C*.

**Supplementary Figure 7.**
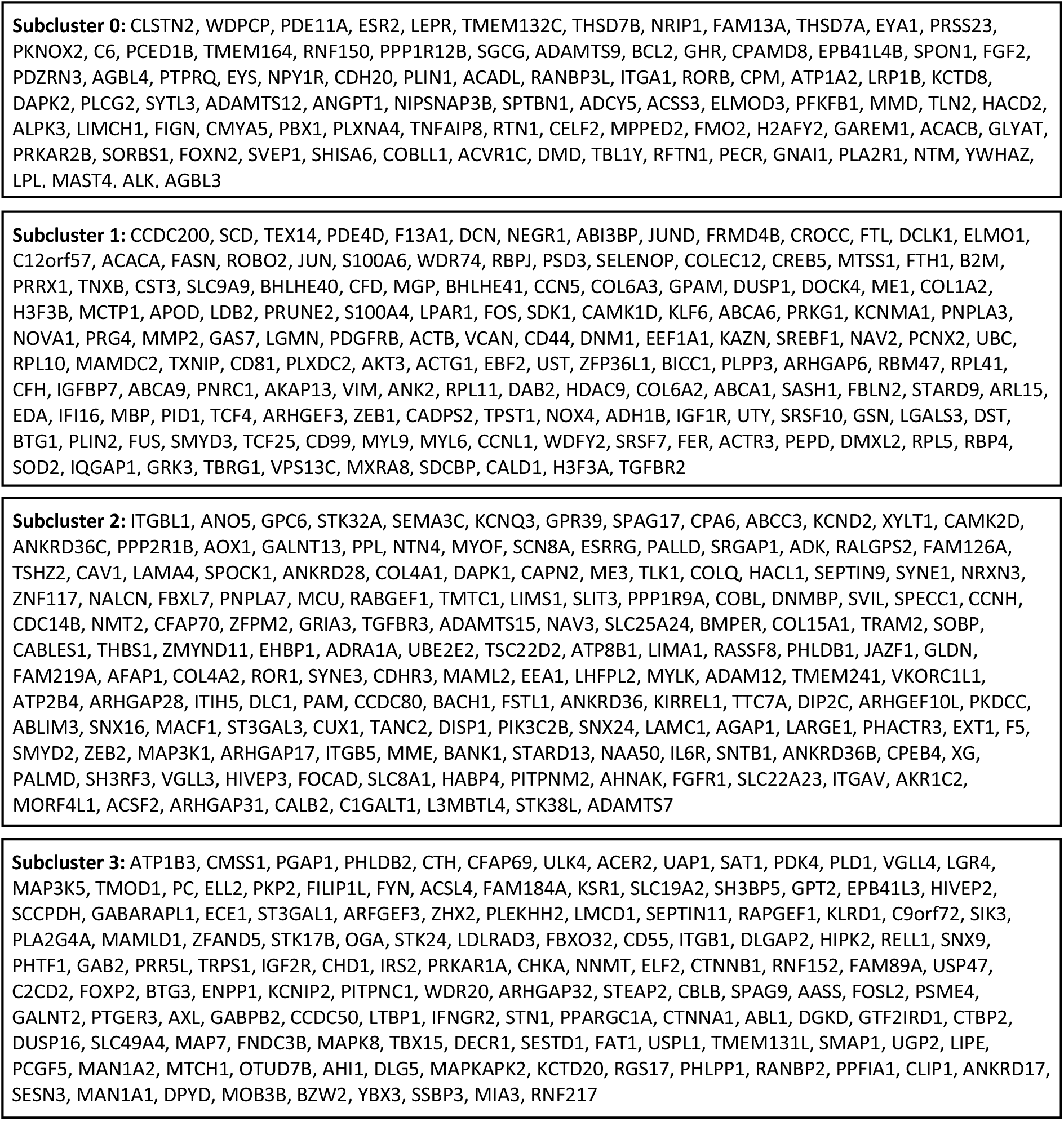
List of 535 Unique Differentially Expressed Genes Within Each Subcluster of Adipocytes Identified by Bioinformatic Analysis of n=21 Infrapatellar Fat Pad Samples from snRNA-seq. Genes are ordered based on decreasing average log2 fold change. Unknown, non-coding genes and genes that were duplicated across subclusters were removed. *Also see Fig. 2, Supplementary Fig. 6, and Supplementary Table 2C*.

**Supplementary Figure 8:**
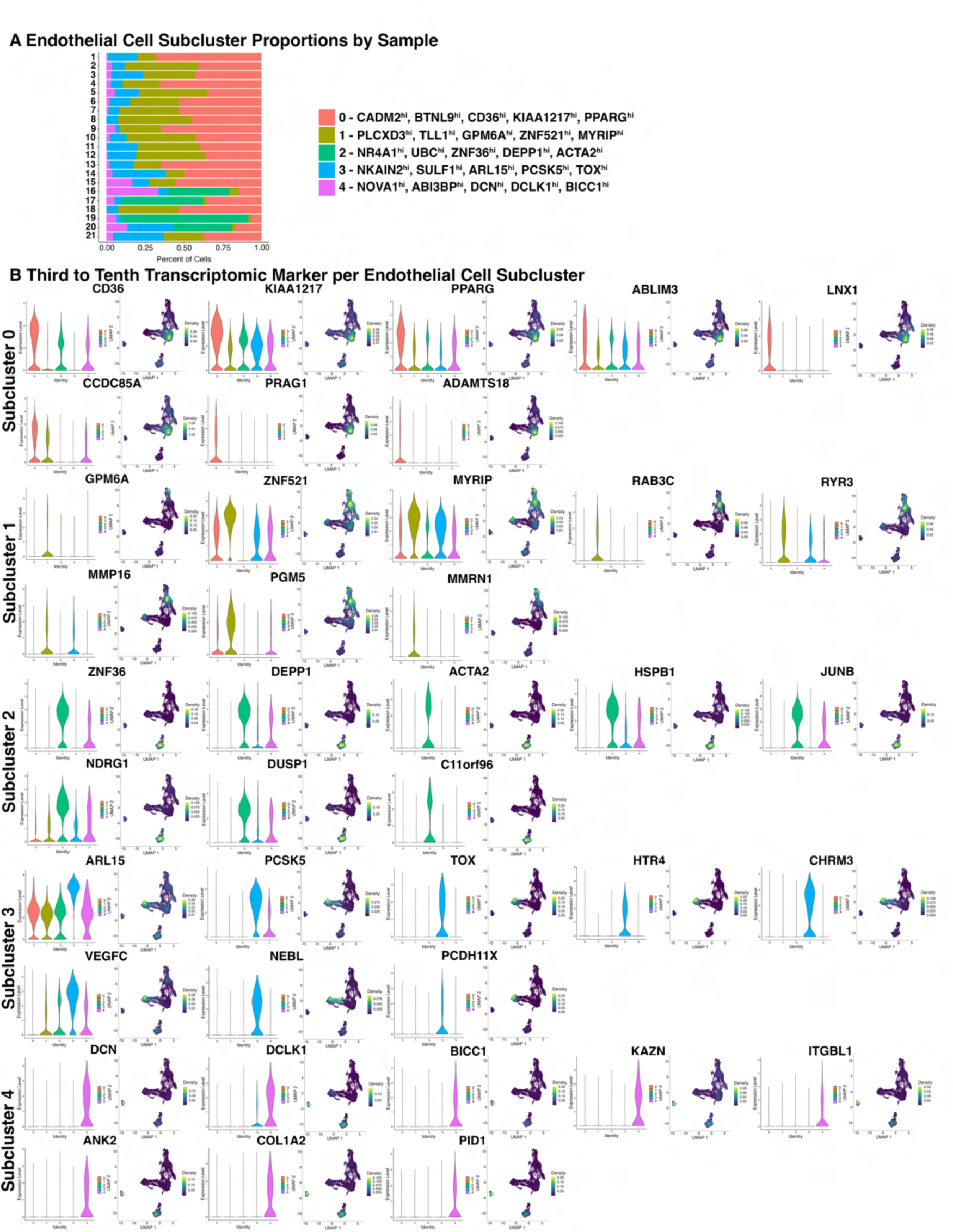
Endothelial Cell Subcluster Proportions and Differentially Expressed Genes (DEGs). A) Bar graph showing the proportion of each endothelial cell subcluster identified from snRNA seq for n=21 IFP samples. Top 5 unique DEGs defining each subcluster are indicated. B) Violin plots and nebulosa plots showing the expression of the third to tenth top unique differentially expressed genes within each endothelial cell subcluster. Nebulosa plots demonstrate expression density from dark purple (low) to yellow (high). *Also see Fig. 2, Supplementary Fig. 9, and Supplementary Table 2D*.

**Supplementary Figure 9.**
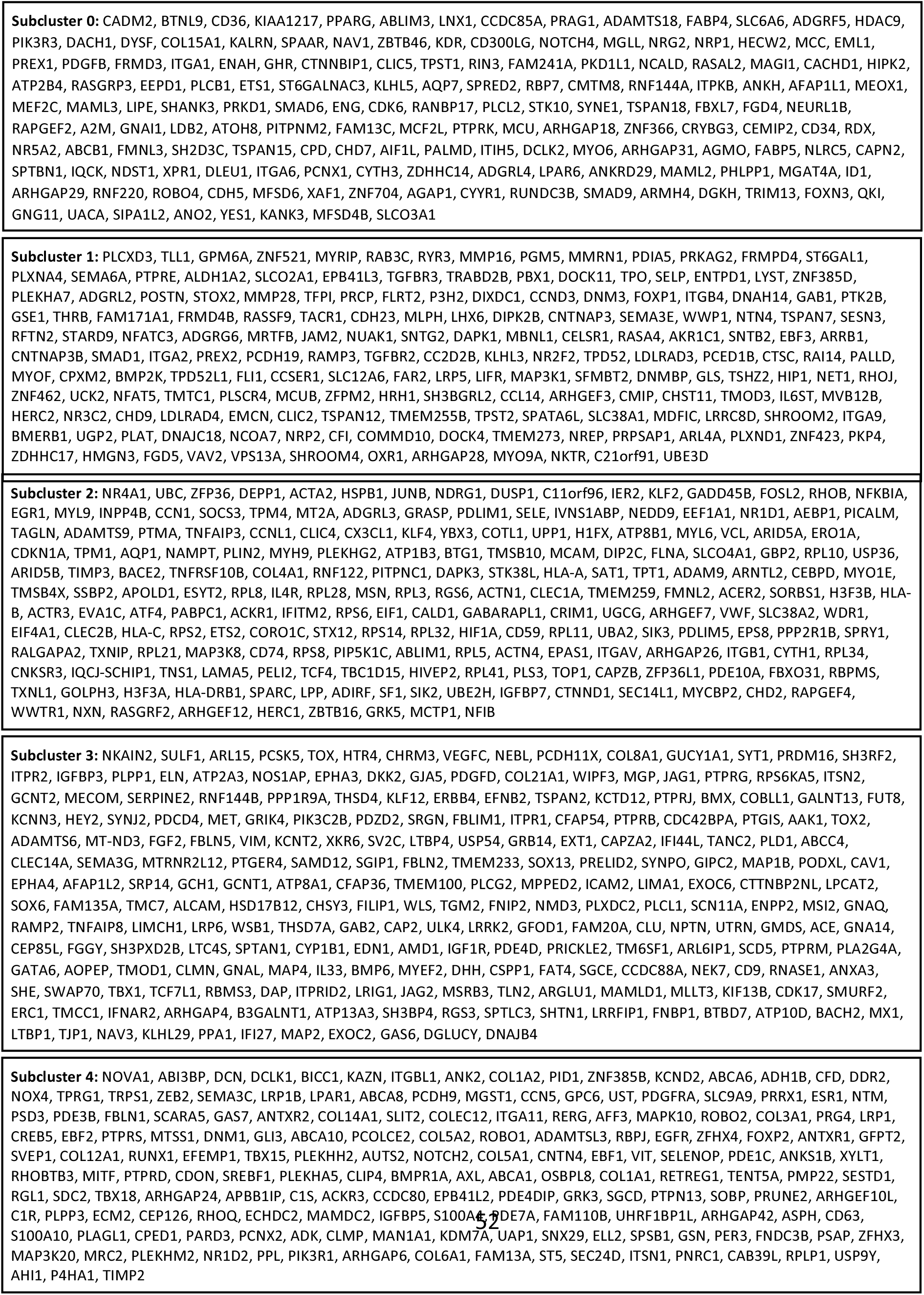
List of 909 Unique Differentially Expressed Genes Within Each Subcluster of Endothelial Cells Identified Using Bioinformatic Analysis of n=21 Infrapatellar Fat Pad Samples from snRNA-seq. Genes are ordered based on decreasing average log2 fold change. Unknown, non-coding genes and genes duplicated across subclusters were removed. *Also see Fig. 2, Supplementary Fig. 8, and Supplementary Table 2D*.

**Supplementary Figure 10.**
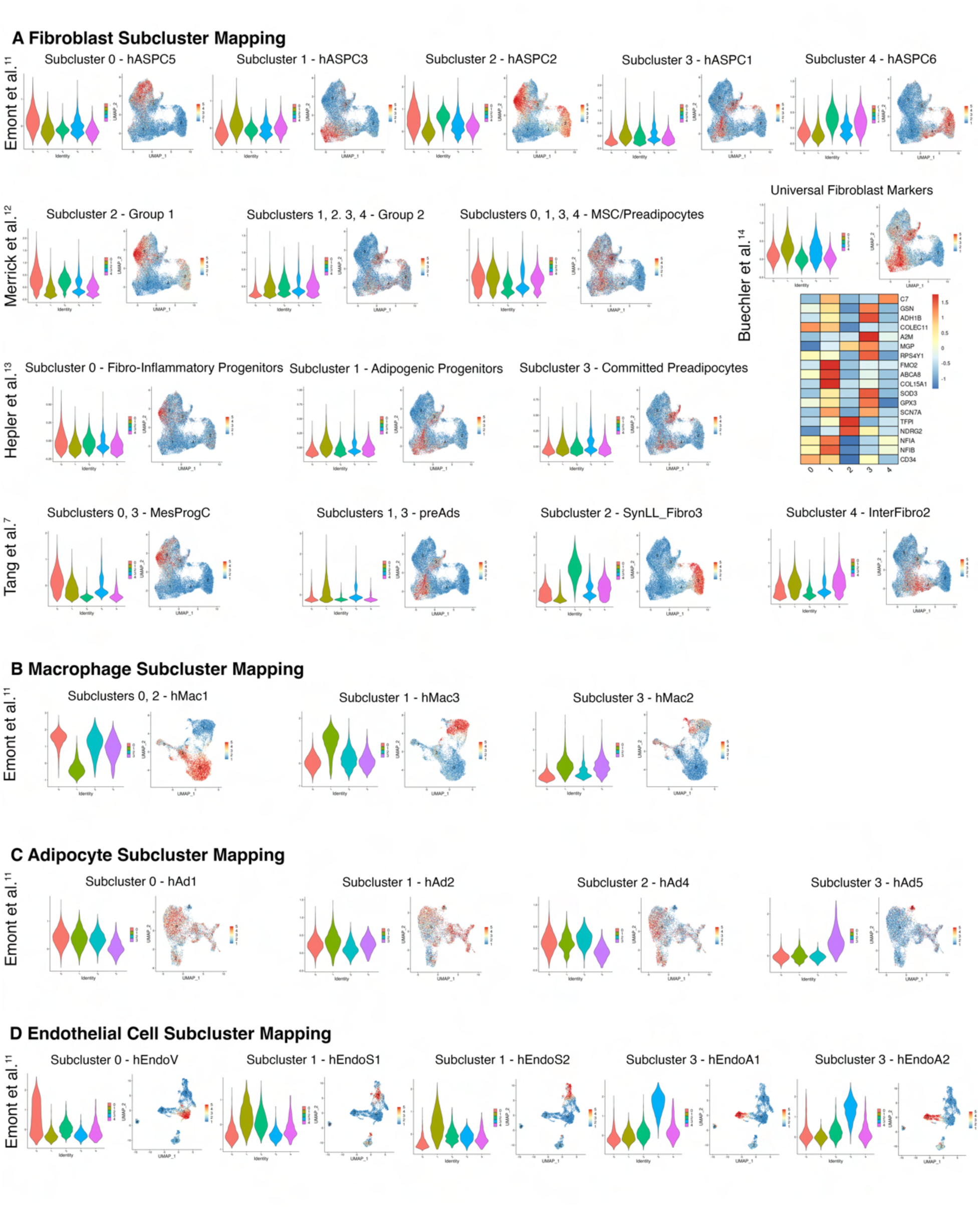
Comparison of Major Cell Population Subclusters in snRNA-seq Data with Published Literature. A) Violin plots and feature plots depicting gene module scores of fibroblast subclusters identified in white adipose tissue by Emont et al.^11^, Merrick et al.^12^, Hepler et al.^13^, and Tang et al.^7^ based on the top differentially expressed genes (DEGs). Mapped populations include: Emont et al.^11^ Subcluster 0 – hASPC5; Subcluster 1 – hASPC3; Subcluster 2 – hASPC2; Subcluster 3 – hASPC1; Subcluster 4 – hASPC6: Merrick et al.^12^ Subcluster 2 – Group 2; Subclusters 1, 2, 3, 4, Group 2; Subclusters 0, 1, 3, 4 – MSC/Preadipocytes: Hepler et al.^13^ Subcluster 0 – Fibro-Inflammatory Progenitors; Subcluster 1 – Adipogenic Progenitors; Subcluster 3 – Committed Preadipocytes: Tang et al.^7^ Subclusters 0, 3 – MesProgCs, Subclusters 1, 3 – preADs, Subcluster 2 – SynLL_Fibro3, Subcluster 4 – InterFibro2. In addition, a violin plot, feature plot and heat map of the expression of universal fibroblast markers identified by Buechler et al^14^ within the snRNA-seq data of the IFP are presented. (B-D) Violin plots and feature plots displaying the gene module scores of (B) macrophage, (C) adipocyte, and (D) endothelial cell subclusters based on DEGs for the respective cell subtypes within white adipose tissue identified by Emont et al.^11^. Mapped populations include: Macrophage Subcluster 0, 2 - hMac1; Subcluster 1 - hMac3; Subcluster 3 - hMac2: Adipocyte Subcluster 0 - hAd1; Subcluster 1 - hAd2; Subcluster 2 - hAd4; Subcluster 3 - hAd5: Endothelial Cell Subcluster 0 - hEndoV (venular); Subcluster 1 - hEndoS1 and hEndoS2 (stalk); Subcluster 3 - hEndoA1 and hEndoA2 (arteriolar).

**Supplementary Figure 11.**
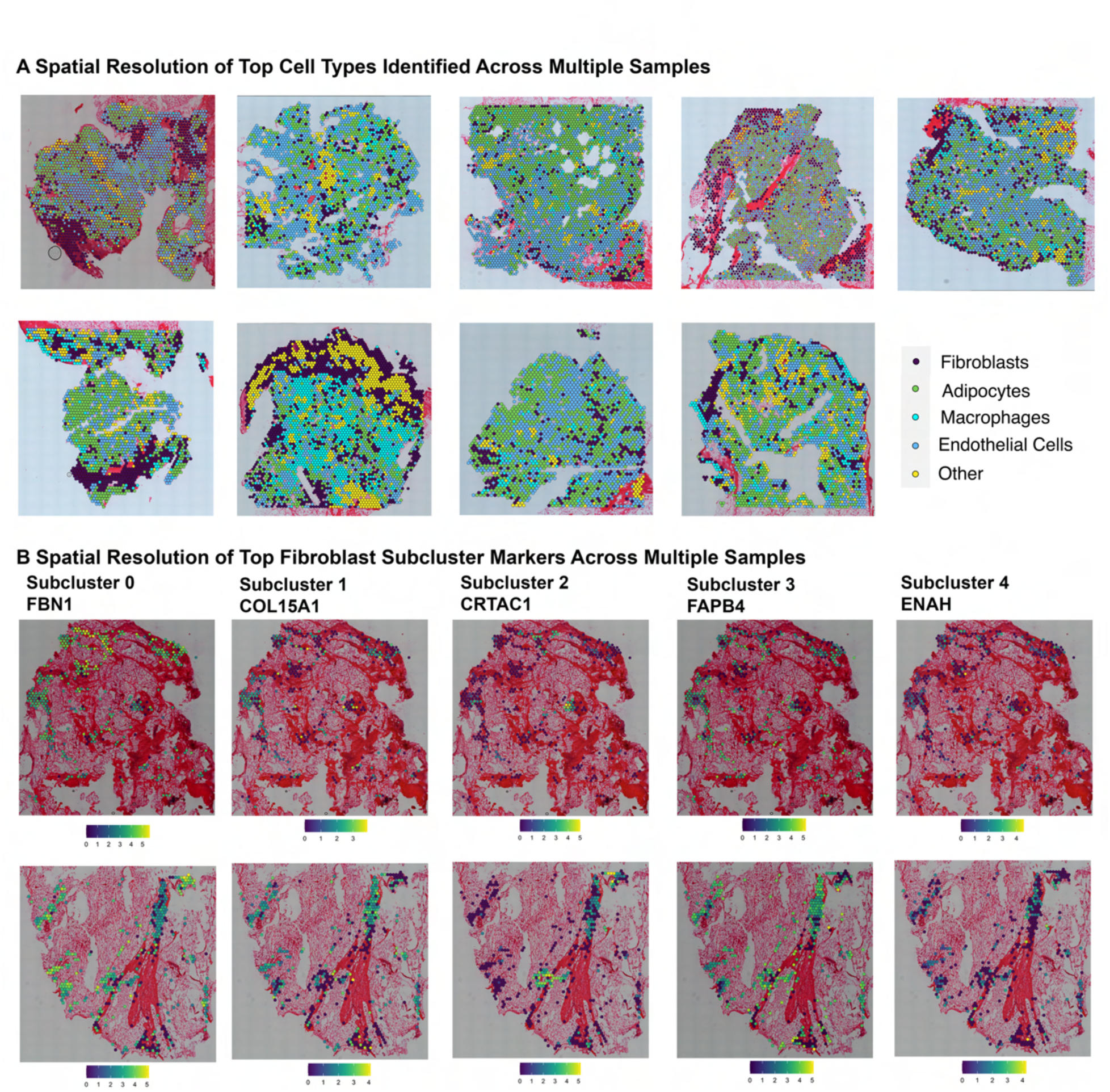
Spatial Profiling of Major Cell Populations Identified in Knee Osteoarthritis Infrapatellar Fat Pads (IFPs). A) Spatial resolution of all major cell types using spatial sequencing across remaining 9 IFP samples. B) Spatial resolution of top transcriptomic markers of each fibroblast subcluster across two independent samples including: Subcluster 0 – FBN1 (DEG 1), Subcluster 1 – COL15A1 (DEG 10), Subcluster 2 – CRTAC1 (DEG 1), Subcluster 3 – FABP4 (DEG 2), Subcluster 4 – ENAH (DEG 5). *Also see Fig. 3*

