## Supplementary Table 1 for "Cell and Transcriptomic Diversity of Infrapatellar Fat Pad during Knee Osteoarthritis"

**Supplementary Table 1: Patient demographics and anthropometrics.** Knee OA infrapatellar fat pad (KOA-IFP) samples were collected from participants in the LEAP OA end stage knee cohort (Schroeder Arthritis Insititute). Each KOA-IFP sample was taken from discarded tissues after total knee arthroplasty. Sex and body mass index (calculated from height and weight) were derived from patient reported measures within study questionnaires. Healthy control donor samples were collected post-mortem <4 hours after death with no history of musculoskeletal disease. Check marks indicate that the sample was used for the indicated experiment.

| Sample ID | Age | Sex | Body Mass Index ($kg/m^2$) | BMI Category ($kg/m^2$) | OA Status | Single Nucleus RNA Sequencing | Spatial Sequencing | Metabolomics |
|---|---|---|---|---|---|---|---|---|
| 1 | 61 | M | 20.96 | Normal BMI 18.5-25 | Knee OA | ✓ | ✓ | |
| 2 | 61 | M | 24.79 | Normal BMI 18.5-25 | Knee OA | ✓ | ✓ | |
| 3 | 55 | M | 24.01 | Normal BMI 18.5-25 | Knee OA | ✓ | ✓ | |
| 4 | 54 | M | 23.6 | Normal BMI 18.5-25 | Knee OA | ✓ | ✓ | |
| 5 | 56 | F | 21.66 | Normal BMI 18.5-25 | Knee OA | ✓ | ✓ | |
| 6 | 58 | F | 21.02 | Normal BMI 18.5-25 | Knee OA | ✓ | | |
| 7 | 61 | F | 23.92 | Normal BMI 18.5-25 | Knee OA | ✓ | ✓ | |
| 8 | 58 | M | 38.48 | Obese BMI 30-40 | Knee OA | ✓ | ✓ | |
| 9 | 57 | M | 34.54 | Obese BMI 30-40 | Knee OA | ✓ | ✓ | |
| 10 | 55 | M | 33.79 | Obese BMI 30-40 | Knee OA | ✓ | ✓ | |
| 11 | 61 | F | 34.6 | Obese BMI 30-40 | Knee OA | ✓ | | |
| 12 | 60 | F | 38.51 | Obese BMI 30-40 | Knee OA | ✓ | ✓ | |
| 13 | 61 | F | 35.9 | Obese BMI 30-40 | Knee OA | ✓ | ✓ | |
| 14 | 57 | F | 39.2 | Obese BMI 30-40 | Knee OA | ✓ | ✓ | |
| 15 | 61 | F | 37.8 | Obese BMI 30-40 | Knee OA | ✓ | | |
| 16 | 49 | M | 22.3 | Normal BMI 18.5-25 | Healthy Control Donor | ✓ | | |
| 17 | 71 | M | 21.1 | Normal BMI 18.5-25 | Healthy Control Donor | ✓ | | |
| 18 | 61 | M | 19.9 | Normal BMI 18.5-25 | Healthy Control Donor | ✓ | | |
| 19 | 45 | M | 25.5 | Overweight BMI 25-30 | Healthy Control Donor | ✓ | | |
| 20 | 66 | F | 28.8 | Overweight BMI 25-30 | Healthy Control Donor | ✓ | | |
| 21 | 59 | F | 26.6 | Overweight BMI 25-30 | Healthy Control Donor | ✓ | | |
| 22 | 66 | F | 23.42 | Normal BMI 18.5-25 | Knee OA | | | ✓ |
| 23 | 70 | F | 21.88 | Normal BMI 18.5-25 | Knee OA | | | ✓ |
| 24 | 72 | M | 22.08 | Normal BMI 18.5-25 | Knee OA | | | ✓ |
| 25 | 66 | F | 21 | Normal BMI 18.5-25 | Knee OA | | | ✓ |
| 26 | 57 | F | 24.8 | Normal BMI 18.5-25 | Knee OA | | | ✓ |
| 27 | 74 | M | 33.3 | Obese BMI 30-40 | Knee OA | | | ✓ |
| 28 | 65 | M | 35.49 | Obese BMI 30-40 | Knee OA | | | ✓ |
| 29 | 70 | F | 32.39 | Obese BMI 30-40 | Knee OA | | | ✓ |
| 30 | 53 | F | 35.5 | Obese BMI 30-40 | Knee OA | | | ✓ |
| 31 | 78 | F | 36.1 | Obese BMI 30-40 | Knee OA | | | ✓ |
