## Supplementary Table 2 for "Cell and Transcriptomic Diversity of Infrapatellar Fat Pad during Knee Osteoarthritis"

**Supplementary Table 2A: Fibroblast subcluster transcriptomic profiles.** Top 30 unique transcriptomic markers of identified fibroblast subclusters from n=21 IFPs. Genes are organized by decreasing average Log2 fold change (FC) filtered by adjusted p < 0.05, log2FC ≥ 0.5, and min.pct ≥ 0.25. Genes duplicated across multiple subclusters, unknown genes, and non-coding genes were not included.

| gene | p_val | avg_log2FC | pct.1 | pct.2 | p_val_adj | cluster |
|---|---|---|---|---|---|---|
| FBN1 | 0 | 1.734785273 | 0.937 | 0.752 | 0 | 0 |
| PXDNL | 0 | 1.683954325 | 0.589 | 0.21 | 0 | 0 |
| PIEZO2 | 0 | 1.572391285 | 0.44 | 0.082 | 0 | 0 |
| GUCY1A2 | 0 | 1.471070056 | 0.381 | 0.059 | 0 | 0 |
| PTGIS | 0 | 1.470975538 | 0.538 | 0.113 | 0 | 0 |
| ITGA11 | 0 | 1.436356409 | 0.815 | 0.312 | 0 | 0 |
| DPP4 | 0 | 1.422835663 | 0.491 | 0.062 | 0 | 0 |
| DOCK4 | 0 | 1.383776483 | 0.72 | 0.423 | 0 | 0 |
| TGFBR3 | 0 | 1.381785783 | 0.836 | 0.606 | 0 | 0 |
| NTN1 | 0 | 1.371787541 | 0.644 | 0.246 | 0 | 0 |
| SMURF2 | 0 | 1.334990735 | 0.73 | 0.443 | 0 | 0 |
| PCOLCE2 | 0 | 1.331439713 | 0.697 | 0.419 | 0 | 0 |
| PAMR1 | 0 | 1.32565553 | 0.438 | 0.083 | 0 | 0 |
| TRIO | 0 | 1.247345887 | 0.882 | 0.622 | 0 | 0 |
| PDZRN4 | 0 | 1.223451735 | 0.405 | 0.164 | 0 | 0 |
| FNDC1 | 0 | 1.198954196 | 0.396 | 0.111 | 0 | 0 |
| MFAP5 | 0 | 1.186781675 | 0.653 | 0.278 | 0 | 0 |
| HMCN1 | 0 | 1.176926003 | 0.661 | 0.4 | 0 | 0 |
| SMOC2 | 0 | 1.171368476 | 0.448 | 0.149 | 0 | 0 |
| NOX4 | 0 | 1.148684876 | 0.748 | 0.473 | 0 | 0 |
| NHSL1 | 0 | 1.147104902 | 0.561 | 0.213 | 0 | 0 |
| ZEB1 | 0 | 1.140687171 | 0.961 | 0.819 | 0 | 0 |
| LTBP2 | 0 | 1.113126552 | 0.465 | 0.147 | 0 | 0 |
| ADGRD1 | 0 | 1.07915696 | 0.627 | 0.278 | 0 | 0 |
| STK32B | 0 | 1.054613096 | 0.282 | 0.093 | 0 | 0 |
| NTM | 0 | 1.041349918 | 0.729 | 0.355 | 0 | 0 |
| NOVA1 | 0 | 1.035560835 | 0.958 | 0.851 | 0 | 0 |
| SDK1 | 0 | 1.011179898 | 0.865 | 0.745 | 0 | 0 |
| ROBO2 | 7.55E-217 | 1.00896225 | 0.388 | 0.23 | 2.35E-212 | 0 |
| FHOD3 | 0 | 0.984999384 | 0.375 | 0.139 | 0 | 0 |
| ABCA10 | 0 | 2.429033726 | 0.72 | 0.249 | 0 | 1 |
| KCND2 | 0 | 1.849184701 | 0.591 | 0.18 | 0 | 1 |
| ABCA6 | 0 | 1.655734353 | 0.841 | 0.371 | 0 | 1 |
| ABCA8 | 0 | 1.454307244 | 0.845 | 0.496 | 0 | 1 |
| ABCA9 | 0 | 1.403400228 | 0.751 | 0.301 | 0 | 1 |
| PTGFR | 0 | 1.369929098 | 0.702 | 0.269 | 0 | 1 |
| APOD | 0 | 1.355469151 | 0.422 | 0.104 | 0 | 1 |
| NEGR1 | 0 | 1.326814166 | 0.951 | 0.608 | 0 | 1 |
| NRP1 | 0 | 1.321139479 | 0.719 | 0.394 | 0 | 1 |
| COL15A1 | 0 | 1.305196651 | 0.504 | 0.155 | 0 | 1 |
| FGF7 | 0 | 1.254095192 | 0.61 | 0.224 | 0 | 1 |
| CCDC102B | 0 | 1.246711844 | 0.396 | 0.167 | 0 | 1 |
| TMEM132C | 0 | 1.245190103 | 0.414 | 0.077 | 0 | 1 |
| ANGPT1 | 0 | 1.244135565 | 0.553 | 0.208 | 0 | 1 |
| BMPER | 0 | 1.223671408 | 0.419 | 0.126 | 0 | 1 |
| PCDH9 | 0 | 1.222974703 | 0.686 | 0.384 | 0 | 1 |
| NAV2 | 0 | 1.190919947 | 0.794 | 0.425 | 0 | 1 |
| LEPR | 0 | 1.162723812 | 0.495 | 0.228 | 0 | 1 |
| GHR | 0 | 1.14154993 | 0.806 | 0.47 | 0 | 1 |
| LAMA2 | 0 | 1.126830497 | 0.841 | 0.563 | 0 | 1 |
| SMOC1 | 0 | 1.107928303 | 0.403 | 0.096 | 0 | 1 |
| SLIT3 | 0 | 1.048724208 | 0.689 | 0.305 | 0 | 1 |
| EYA1 | 0 | 1.04538102 | 0.259 | 0.035 | 0 | 1 |
| CXCL14 | 0 | 1.040081074 | 0.261 | 0.036 | 0 | 1 |
| LMO3 | 0 | 1.039617363 | 0.445 | 0.141 | 0 | 1 |
| CCDC85A | 0 | 1.014094076 | 0.556 | 0.255 | 0 | 1 |
| LDB2 | 0 | 1.012008623 | 0.783 | 0.462 | 0 | 1 |
| IGFBP7 | 0 | 0.986730578 | 0.444 | 0.178 | 0 | 1 |
| CNTN4 | 4.27E-158 | 0.981340393 | 0.281 | 0.163 | 1.33E-153 | 1 |
| ACVR2A | 0 | 0.968786053 | 0.579 | 0.273 | 0 | 1 |
| CRTAC1 | 0 | 3.513972339 | 0.974 | 0.165 | 0 | 2 |
| CLIC5 | 0 | 3.419483797 | 0.786 | 0.037 | 0 | 2 |
| PRG4 | 0 | 3.334914791 | 0.973 | 0.545 | 0 | 2 |
| FN1 | 0 | 3.314091069 | 0.994 | 0.656 | 0 | 2 |
| SEMA3A | 0 | 3.122961449 | 0.794 | 0.198 | 0 | 2 |
| HTRA1 | 0 | 3.052520609 | 0.836 | 0.238 | 0 | 2 |
| SEMA5A | 0 | 2.911772497 | 0.784 | 0.069 | 0 | 2 |
| CLU | 0 | 2.838984231 | 0.764 | 0.272 | 0 | 2 |
| ANK3 | 0 | 2.76300859 | 0.8 | 0.088 | 0 | 2 |
| COL22A1 | 0 | 2.73563487 | 0.592 | 0.016 | 0 | 2 |
| ERRFI1 | 0 | 2.700706687 | 0.823 | 0.2 | 0 | 2 |
| ZNF385B | 0 | 2.691728596 | 0.983 | 0.57 | 0 | 2 |
| GPR1 | 0 | 2.529331577 | 0.555 | 0.087 | 0 | 2 |
| HTRA4 | 0 | 2.422607741 | 0.56 | 0.045 | 0 | 2 |
| ITGBL1 | 0 | 2.417292805 | 0.931 | 0.44 | 0 | 2 |
| MT2A | 0 | 2.382742184 | 0.688 | 0.178 | 0 | 2 |
| RGS6 | 0 | 2.332412789 | 0.498 | 0.054 | 0 | 2 |
| NTN4 | 0 | 2.167388728 | 0.839 | 0.265 | 0 | 2 |
| SHANK2 | 0 | 2.142788268 | 0.624 | 0.023 | 0 | 2 |
| MMP3 | 0 | 2.088978662 | 0.309 | 0.038 | 0 | 2 |
| BCAT1 | 0 | 2.042667307 | 0.81 | 0.228 | 0 | 2 |
| DAPK1 | 0 | 2.031579384 | 0.768 | 0.106 | 0 | 2 |
| SPARCL1 | 0 | 2.02035861 | 0.765 | 0.238 | 0 | 2 |
| MYPN | 0 | 1.995943822 | 0.576 | 0.02 | 0 | 2 |
| SOX5 | 0 | 1.911208983 | 0.996 | 0.92 | 0 | 2 |
| PCSK6 | 0 | 1.844911507 | 0.871 | 0.319 | 0 | 2 |
| PPP4R4 | 0 | 1.785779022 | 0.455 | 0.046 | 0 | 2 |
| SV2B | 0 | 1.783252337 | 0.598 | 0.031 | 0 | 2 |
| VEGFC | 0 | 1.777840068 | 0.516 | 0.057 | 0 | 2 |
| TIMP1 | 0 | 1.7745506 | 0.48 | 0.157 | 0 | 2 |
| GPAM | 0 | 2.341088926 | 0.273 | 0.096 | 0 | 3 |
| F13A1 | 0 | 2.16299891 | 0.251 | 0.051 | 0 | 3 |
| FABP4 | 5.21E-271 | 2.319047946 | 0.305 | 0.132 | 1.62E-266 | 3 |
| FTL | 4.10E-148 | 2.023678465 | 0.418 | 0.319 | 1.28E-143 | 3 |
| FTH1 | 1.43E-59 | 1.681705105 | 0.36 | 0.327 | 4.46E-55 | 3 |
| ME1 | 5.23E-61 | 1.57967291 | 0.264 | 0.199 | 1.63E-56 | 3 |
| JUND | 6.42E-57 | 1.573507259 | 0.326 | 0.279 | 2.00E-52 | 3 |
| PDE3B | 1.71E-95 | 1.546246059 | 0.37 | 0.294 | 5.32E-91 | 3 |
| S100A6 | 1.90E-63 | 1.545294434 | 0.425 | 0.414 | 5.92E-59 | 3 |
| SELENOP | 5.91E-29 | 1.497371508 | 0.297 | 0.286 | 1.84E-24 | 3 |
| FRMD4B | 2.68E-78 | 1.464725819 | 0.262 | 0.179 | 8.33E-74 | 3 |
| S100A4 | 2.92E-16 | 1.328295762 | 0.377 | 0.438 | 9.09E-12 | 3 |
| JUN | 6.50E-77 | 1.318373116 | 0.333 | 0.256 | 2.03E-72 | 3 |
| CST3 | 1.53E-18 | 1.306795433 | 0.304 | 0.311 | 4.75E-14 | 3 |
| KLF6 | 6.53E-08 | 1.24340144 | 0.243 | 0.256 | 0.0020329 | 3 |
| ACTB | 9.11E-11 | 1.212901448 | 0.31 | 0.344 | 2.84E-06 | 3 |
| UBC | 3.26E-16 | 1.199045389 | 0.254 | 0.247 | 1.02E-11 | 3 |
| VIM | 1.41E-38 | 1.108225519 | 0.548 | 0.698 | 4.39E-34 | 3 |
| IGFBP6 | 4.47E-09 | 1.095197625 | 0.246 | 0.251 | 0.0001392 | 3 |
| ACACA | 1.21E-12 | 0.8087 | 0.301 | 0.474 | 3.76E-08 | 3 |
| RBM25 | 1.98E-23 | 0.783143825 | 0.398 | 0.698 | 6.18E-19 | 3 |
| PSAP | 3.06E-12 | 0.778420804 | 0.259 | 0.397 | 9.54E-08 | 3 |
| ZFP36L1 | 5.60E-08 | 0.765787271 | 0.309 | 0.464 | 0.0017459 | 3 |
| LGALS3 | 1.87E-25 | 0.732692581 | 0.257 | 0.434 | 5.83E-21 | 3 |
| SERPING1 | 1.78E-19 | 0.715316499 | 0.217 | 0.349 | 5.54E-15 | 3 |
| ANXA1 | 1.47E-10 | 0.708740389 | 0.348 | 0.546 | 4.59E-06 | 3 |
| TXNIP | 3.09E-11 | 0.700524014 | 0.274 | 0.414 | 9.64E-07 | 3 |
| PFKFB3 | 1.20E-21 | 0.67477932 | 0.169 | 0.278 | 3.73E-17 | 3 |
| GPNMB | 5.33E-25 | 0.63667076 | 0.175 | 0.294 | 1.66E-20 | 3 |
| COL6A1 | 4.83E-23 | 0.60886209 | 0.371 | 0.634 | 1.50E-18 | 3 |
| MGAT4C | 0 | 2.236845534 | 0.552 | 0.118 | 0 | 4 |
| GRIP1 | 0 | 1.378623759 | 0.365 | 0.078 | 0 | 4 |
| BTBD11 | 0 | 1.325020382 | 0.328 | 0.032 | 0 | 4 |
| IGF1 | 1.42E-172 | 1.264360283 | 0.444 | 0.238 | 4.43E-168 | 4 |
| C1GALT1 | 0 | 1.183228707 | 0.777 | 0.461 | 0 | 4 |
| ENAH | 0 | 1.170289073 | 0.752 | 0.423 | 0 | 4 |
| COL14A1 | 0 | 1.113026574 | 0.741 | 0.407 | 0 | 4 |
| PDE10A | 1.42E-178 | 1.089691855 | 0.459 | 0.248 | 4.43E-174 | 4 |
| PLPP1 | 1.00E-201 | 1.089498501 | 0.748 | 0.53 | 3.12E-197 | 4 |
| ZIC1 | 0 | 1.060503445 | 0.439 | 0.124 | 0 | 4 |
| UNC5C | 0 | 1.043761736 | 0.602 | 0.23 | 0 | 4 |
| KCNH1 | 0 | 1.038605593 | 0.298 | 0.049 | 0 | 4 |
| ENPP2 | 0 | 1.006571646 | 0.401 | 0.13 | 0 | 4 |
| RUNX1 | 2.91E-164 | 1.004993015 | 0.702 | 0.507 | 9.06E-160 | 4 |
| PPP3CA | 0 | 1.002188396 | 0.939 | 0.748 | 0 | 4 |
| GLI3 | 8.40E-242 | 0.91317705 | 0.871 | 0.687 | 2.62E-237 | 4 |
| FGF14 | 7.53E-194 | 0.905879977 | 0.647 | 0.398 | 2.34E-189 | 4 |
| KCND3 | 8.67E-304 | 0.860089036 | 0.345 | 0.111 | 2.70E-299 | 4 |
| TMEM108 | 2.71E-168 | 0.855491426 | 0.48 | 0.257 | 8.45E-164 | 4 |
| NID2 | 3.29E-216 | 0.845077788 | 0.412 | 0.192 | 1.03E-211 | 4 |
| ADAMTS19 | 6.28E-181 | 0.825121235 | 0.27 | 0.1 | 1.96E-176 | 4 |
| IGFBP4 | 2.51E-209 | 0.810284418 | 0.533 | 0.293 | 7.81E-205 | 4 |
| AHR | 6.68E-179 | 0.795404897 | 0.63 | 0.416 | 2.08E-174 | 4 |
| VAV3 | 1.43E-278 | 0.786639012 | 0.329 | 0.108 | 4.44E-274 | 4 |
| DIAPH2-AS1 | 1.44E-133 | 0.775904984 | 0.331 | 0.165 | 4.48E-129 | 4 |
| WNT5B | 6.41E-217 | 0.77070089 | 0.408 | 0.178 | 2.00E-212 | 4 |
| RSPO3 | 6.62E-272 | 0.753506529 | 0.342 | 0.117 | 2.06E-267 | 4 |
| TMTC2 | 1.46E-124 | 0.747131831 | 0.308 | 0.148 | 4.56E-120 | 4 |
| TBX5 | 1.01E-284 | 0.745392027 | 0.361 | 0.128 | 3.16E-280 | 4 |
| ST5 | 7.75E-197 | 0.73823137 | 0.732 | 0.52 | 2.41E-192 | 4 |

**Supplementary Table 2B: Macrophage subcluster transcriptomic profiles.** Top 30 unique transcriptomic markers of identified macrophage subclusters from n=21 IFPs. Genes are organized by decreasing average Log2 fold change (FC) filtered by adjusted p < 0.05, log2FC ≥ 0.5, and min.pct ≥ 0.25. Genes duplicated across multiple subclusters, unknown genes, and non-coding genes were not included.

| gene | p_val | avg_log2FC | pct.1 | pct.2 | p_val_adj | cluster |
|---|---|---|---|---|---|---|
| P2RY14 | 0 | 2.103037548 | 0.768 | 0.123 | 0 | 0 |
| MAN1A1 | 0 | 1.785470283 | 0.938 | 0.537 | 0 | 0 |
| FGF13 | 0 | 1.750181744 | 0.804 | 0.256 | 0 | 0 |
| DSCAML1 | 0 | 1.724764307 | 0.677 | 0.116 | 0 | 0 |
| PID1 | 0 | 1.696207899 | 0.619 | 0.161 | 0 | 0 |
| NAV2 | 0 | 1.680329873 | 0.954 | 0.436 | 0 | 0 |
| PDE4D | 0 | 1.533801412 | 0.984 | 0.792 | 0 | 0 |
| AFF3 | 0 | 1.500223725 | 0.612 | 0.161 | 0 | 0 |
| LSAMP | 0 | 1.429391453 | 0.49 | 0.149 | 0 | 0 |
| ADGRL3 | 0 | 1.396131373 | 0.471 | 0.087 | 0 | 0 |
| RGL1 | 0 | 1.392270393 | 0.94 | 0.553 | 0 | 0 |
| NRP1 | 0 | 1.362635661 | 0.964 | 0.568 | 0 | 0 |
| SLC40A1 | 0 | 1.342207289 | 0.744 | 0.24 | 0 | 0 |
| CTSC | 0 | 1.3398606 | 0.733 | 0.271 | 0 | 0 |
| ROR1 | 0 | 1.300318291 | 0.498 | 0.09 | 0 | 0 |
| CSGALNACT1 | 0 | 1.294592136 | 0.585 | 0.132 | 0 | 0 |
| NPAS3 | 0 | 1.290076353 | 0.612 | 0.268 | 0 | 0 |
| DAAM2 | 0 | 1.287543013 | 0.54 | 0.066 | 0 | 0 |
| MPPED2 | 0 | 1.277263341 | 0.69 | 0.199 | 0 | 0 |
| AUTS2 | 0 | 1.274215649 | 0.964 | 0.695 | 0 | 0 |
| STARD13 | 0 | 1.271071319 | 0.96 | 0.631 | 0 | 0 |
| WWP1 | 0 | 1.226785409 | 0.961 | 0.649 | 0 | 0 |
| SCN9A | 0 | 1.22490961 | 0.943 | 0.433 | 0 | 0 |
| HRH1 | 0 | 1.216640276 | 0.878 | 0.407 | 0 | 0 |
| WLS | 0 | 1.191570892 | 0.839 | 0.305 | 0 | 0 |
| IQGAP2 | 0 | 1.178544092 | 0.965 | 0.601 | 0 | 0 |
| CCDC141 | 0 | 1.167142958 | 0.724 | 0.156 | 0 | 0 |
| MRC1 | 0 | 1.15774462 | 0.98 | 0.636 | 0 | 0 |
| STON2 | 0 | 1.156172628 | 0.724 | 0.154 | 0 | 0 |
| GNG2 | 0 | 1.098893752 | 0.814 | 0.336 | 0 | 0 |
| FN1 | 0 | 3.026191061 | 0.81 | 0.241 | 0 | 1 |
| TPRG1 | 0 | 2.974092641 | 0.746 | 0.105 | 0 | 1 |
| KCNQ3 | 0 | 2.544930482 | 0.595 | 0.045 | 0 | 1 |
| PDE3A | 0 | 2.440180397 | 0.662 | 0.052 | 0 | 1 |
| FMNL2 | 0 | 2.296649198 | 0.94 | 0.615 | 0 | 1 |
| ABCC3 | 0 | 2.283442708 | 0.888 | 0.129 | 0 | 1 |
| APOE | 0 | 2.198205764 | 0.566 | 0.099 | 0 | 1 |
| APBB1IP | 0 | 2.149370365 | 0.931 | 0.355 | 0 | 1 |
| SLC11A1 | 0 | 1.756047176 | 0.687 | 0.136 | 0 | 1 |
| PLXDC2 | 0 | 1.751039402 | 0.986 | 0.815 | 0 | 1 |
| IFI30 | 0 | 1.741122004 | 0.719 | 0.161 | 0 | 1 |
| ZNF804A | 0 | 1.724536807 | 0.589 | 0.07 | 0 | 1 |
| LHFPL2 | 0 | 1.683564935 | 0.788 | 0.279 | 0 | 1 |
| TIMD4 | 0 | 1.65198548 | 0.57 | 0.073 | 0 | 1 |
| DLEU1 | 0 | 1.649548681 | 0.895 | 0.517 | 0 | 1 |
| FERMT2 | 0 | 1.570781566 | 0.543 | 0.059 | 0 | 1 |
| TANC2 | 0 | 1.493822767 | 0.867 | 0.466 | 0 | 1 |
| DLEU7 | 0 | 1.475609551 | 0.522 | 0.094 | 0 | 1 |
| ALCAM | 0 | 1.461606497 | 0.518 | 0.182 | 0 | 1 |
| DOCK10 | 0 | 1.447607763 | 0.762 | 0.244 | 0 | 1 |
| SNTB1 | 0 | 1.385066835 | 0.573 | 0.145 | 0 | 1 |
| SKIL | 0 | 1.382599644 | 0.693 | 0.174 | 0 | 1 |
| DOCK4 | 0 | 1.366971412 | 0.957 | 0.693 | 0 | 1 |
| PRG4 | 0 | 1.342525101 | 0.672 | 0.244 | 0 | 1 |
| CLEC7A | 0 | 1.338944611 | 0.651 | 0.11 | 0 | 1 |
| PCNX2 | 0 | 1.334879539 | 0.615 | 0.087 | 0 | 1 |
| MSR1 | 0 | 1.326546893 | 0.927 | 0.613 | 0 | 1 |
| L1TD1 | 0 | 1.32277115 | 0.427 | 0.015 | 0 | 1 |
| MERTK | 0 | 1.298830531 | 0.82 | 0.57 | 0 | 1 |
| FCGR3A | 0 | 1.263663722 | 0.52 | 0.052 | 0 | 1 |
| TTN | 2.23E-103 | 2.007419259 | 0.629 | 0.575 | 6.96E-99 | 2 |
| STAB1 | 1.15E-161 | 1.887167001 | 0.688 | 0.713 | 3.59E-157 | 2 |
| AHNAK | 7.25E-131 | 1.830382602 | 0.687 | 0.773 | 2.26E-126 | 2 |
| RBM25 | 2.17E-17 | 1.321463243 | 0.502 | 0.682 | 6.75E-13 | 2 |
| PRRC2C | 3.47E-18 | 1.195198287 | 0.519 | 0.723 | 1.08E-13 | 2 |
| SRRM2 | 1.15E-25 | 1.184018633 | 0.562 | 0.779 | 3.59E-21 | 2 |
| SIGLEC1 | 7.87E-37 | 1.08737796 | 0.587 | 0.75 | 2.45E-32 | 2 |
| TNFAIP2 | 9.59E-33 | 1.038742651 | 0.629 | 0.857 | 2.99E-28 | 2 |
| FCGRT | 2.40E-13 | 1.026093195 | 0.524 | 0.733 | 7.46E-09 | 2 |
| PNISR | 9.26E-14 | 0.982540165 | 0.549 | 0.801 | 2.89E-09 | 2 |
| ARGLU1 | 1.57E-09 | 0.960437136 | 0.54 | 0.803 | 4.90E-05 | 2 |
| LILRB5 | 6.70E-14 | 0.909859483 | 0.56 | 0.804 | 2.09E-09 | 2 |
| MPHOSPH8 | 8.77E-12 | 0.901840944 | 0.311 | 0.555 | 2.73E-07 | 2 |
| BOD1L1 | 1.20E-08 | 0.8973923 | 0.32 | 0.548 | 0.0003737 | 2 |
| RNF213 | 7.92E-14 | 0.866322766 | 0.579 | 0.831 | 2.47E-09 | 2 |
| AP2A2 | 1.69E-30 | 0.85977164 | 0.646 | 0.838 | 5.26E-26 | 2 |
| ZKSCAN1 | 3.69E-11 | 0.84534867 | 0.234 | 0.402 | 1.15E-06 | 2 |
| BRD2 | 1.94E-07 | 0.836805976 | 0.175 | 0.278 | 0.0060547 | 2 |
| DNM1 | 4.19E-18 | 0.812731278 | 0.603 | 0.822 | 1.31E-13 | 2 |
| CCNL2 | 1.70E-10 | 0.798934468 | 0.2 | 0.338 | 5.30E-06 | 2 |
| SREK1 | 7.65E-19 | 0.785176659 | 0.261 | 0.493 | 2.38E-14 | 2 |
| ANKRD36C | 3.26E-17 | 0.768455111 | 0.273 | 0.506 | 1.01E-12 | 2 |
| EIF3A | 5.32E-17 | 0.753749616 | 0.231 | 0.427 | 1.66E-12 | 2 |
| CHD4 | 1.11E-18 | 0.724435788 | 0.187 | 0.351 | 3.47E-14 | 2 |
| MYO15B | 6.82E-23 | 0.722113644 | 0.199 | 0.391 | 2.13E-18 | 2 |
| TMEM259 | 3.44E-19 | 0.714543834 | 0.186 | 0.354 | 1.07E-14 | 2 |
| AP3D1 | 1.15E-15 | 0.712259753 | 0.174 | 0.317 | 3.59E-11 | 2 |
| GAK | 2.04E-19 | 0.709812038 | 0.222 | 0.419 | 6.35E-15 | 2 |
| RBPJ | 1.33E-35 | 0.704514196 | 0.863 | 0.962 | 4.15E-31 | 2 |
| BAZ2A | 1.39E-16 | 0.703579995 | 0.191 | 0.352 | 4.32E-12 | 2 |
| NOVA1 | 0 | 3.234602809 | 0.6 | 0.088 | 0 | 3 |
| ADH1B | 0 | 2.715267617 | 0.411 | 0.056 | 0 | 3 |
| DLC1 | 0 | 2.634805057 | 0.532 | 0.118 | 0 | 3 |
| FBN1 | 2.40E-299 | 2.481451183 | 0.477 | 0.116 | 7.47E-295 | 3 |
| NEGR1 | 0 | 2.477191266 | 0.421 | 0.076 | 0 | 3 |
| DCN | 1.06E-302 | 2.475150699 | 0.571 | 0.176 | 3.31E-298 | 3 |
| DCLK1 | 0 | 2.433346956 | 0.379 | 0.044 | 0 | 3 |
| HSPA1A | 5.56E-199 | 2.394871767 | 0.333 | 0.077 | 1.73E-194 | 3 |
| GPAM | 2.77E-179 | 2.381951745 | 0.316 | 0.076 | 8.64E-175 | 3 |
| JUN | 1.69E-270 | 2.324899605 | 0.578 | 0.2 | 5.26E-266 | 3 |
| ABI3BP | 1.32E-263 | 2.323305455 | 0.42 | 0.1 | 4.12E-259 | 3 |
| SCD | 0 | 2.255881849 | 0.339 | 0.049 | 0 | 3 |
| GHR | 3.81E-235 | 2.23648112 | 0.378 | 0.087 | 1.19E-230 | 3 |
| TSHZ2 | 2.38E-281 | 2.148803894 | 0.381 | 0.074 | 7.42E-277 | 3 |
| TEX14 | 2.40E-302 | 2.086396633 | 0.304 | 0.04 | 7.46E-298 | 3 |
| JUND | 2.90E-143 | 2.081022145 | 0.496 | 0.236 | 9.04E-139 | 3 |
| RBMS3 | 9.40E-187 | 2.015781064 | 0.548 | 0.242 | 2.93E-182 | 3 |
| HSPB1 | 3.87E-139 | 2.002174662 | 0.304 | 0.088 | 1.21E-134 | 3 |
| COL6A3 | 6.35E-203 | 1.998795225 | 0.358 | 0.088 | 1.98E-198 | 3 |
| CCN5 | 1.49E-258 | 1.905814355 | 0.289 | 0.042 | 4.64E-254 | 3 |
| FABP4 | 1.88E-126 | 1.895917058 | 0.389 | 0.151 | 5.86E-122 | 3 |
| NAV3 | 2.07E-165 | 1.873020848 | 0.37 | 0.111 | 6.46E-161 | 3 |
| MGST1 | 1.76E-233 | 1.870122242 | 0.291 | 0.048 | 5.48E-229 | 3 |
| PDE3B | 1.66E-66 | 1.861634782 | 0.457 | 0.3 | 5.17E-62 | 3 |
| NFIB | 1.90E-205 | 1.824991415 | 0.322 | 0.069 | 5.92E-201 | 3 |
| HSPA1B | 7.91E-207 | 1.820034092 | 0.284 | 0.052 | 2.46E-202 | 3 |
| CACNA2D1 | 6.37E-203 | 1.778093938 | 0.285 | 0.054 | 1.98E-198 | 3 |
| COL1A2 | 4.57E-146 | 1.737716072 | 0.326 | 0.096 | 1.42E-141 | 3 |
| PLCB1 | 1.79E-166 | 1.721078019 | 0.298 | 0.071 | 5.58E-162 | 3 |
| RORA | 1.09E-90 | 1.716012299 | 0.534 | 0.359 | 3.39E-86 | 3 |

**Supplementary Table 2C: Adipocyte subcluster transcriptomic profiles.** Top 30 unique transcriptomic markers of identified adipocyte subclusters from n=21 IFPs. Genes are organized by decreasing average Log2 fold change (FC) filtered by adjusted p < 0.05, log2FC ≥ 0.5, and min.pct ≥ 0.25. Genes duplicated across multiple subclusters, unknown genes, and non-coding genes were not included.

| gene | p_val | avg_log2FC | pct.1 | pct.2 | p_val_adj | cluster |
|---|---|---|---|---|---|---|
| CLSTN2 | 0 | 1.58623549 | 0.671 | 0.258 | 0 | 0 |
| WDPCP | 5.53E-138 | 0.98251398 | 0.954 | 0.817 | 1.72E-133 | 0 |
| PDE11A | 4.80E-225 | 0.97573316 | 0.559 | 0.245 | 1.50E-220 | 0 |
| ESR2 | 0 | 0.93806998 | 0.745 | 0.345 | 0 | 0 |
| LEPR | 5.76E-144 | 0.93316347 | 0.58 | 0.344 | 1.79E-139 | 0 |
| TMEM132C | 0 | 0.8547564 | 0.965 | 0.776 | 0 | 0 |
| THSD7B | 2.61E-276 | 0.83684416 | 0.757 | 0.379 | 8.14E-272 | 0 |
| NRIP1 | 4.25E-270 | 0.8108642 | 0.915 | 0.695 | 1.32E-265 | 0 |
| FAM13A | 2.23E-302 | 0.80138638 | 0.956 | 0.808 | 6.94E-298 | 0 |
| THSD7A | 4.61E-225 | 0.77355951 | 0.697 | 0.383 | 1.44E-220 | 0 |
| EYA1 | 9.23E-225 | 0.76953558 | 0.689 | 0.346 | 2.88E-220 | 0 |
| PRSS23 | 1.29E-217 | 0.76479628 | 0.795 | 0.531 | 4.03E-213 | 0 |
| PKNOX2 | 3.87E-215 | 0.76039822 | 0.539 | 0.228 | 1.20E-210 | 0 |
| C6 | 6.50E-177 | 0.7590641 | 0.541 | 0.256 | 2.02E-172 | 0 |
| PCED1B | 4.32E-243 | 0.75858556 | 0.849 | 0.548 | 1.35E-238 | 0 |
| TMEM164 | 5.75E-147 | 0.7550311 | 0.797 | 0.57 | 1.79E-142 | 0 |
| RNF150 | 1.41E-194 | 0.75372577 | 0.881 | 0.651 | 4.38E-190 | 0 |
| PPP1R12B | 2.16E-305 | 0.75064723 | 0.936 | 0.739 | 6.73E-301 | 0 |
| SGCG | 5.27E-250 | 0.74626529 | 0.577 | 0.225 | 1.64E-245 | 0 |
| ADAMTS9 | 3.89E-225 | 0.73667054 | 0.631 | 0.311 | 1.21E-220 | 0 |
| BCL2 | 3.91E-276 | 0.7363708 | 0.952 | 0.772 | 1.22E-271 | 0 |
| GHR | 0 | 0.72910126 | 0.984 | 0.935 | 0 | 0 |
| CPAMD8 | 5.33E-122 | 0.72855546 | 0.344 | 0.13 | 1.66E-117 | 0 |
| EPB41L4B | 4.36E-267 | 0.71621088 | 0.772 | 0.449 | 1.36E-262 | 0 |
| SPON1 | 1.88E-209 | 0.71512116 | 0.64 | 0.36 | 5.87E-205 | 0 |
| FGF2 | 1.80E-205 | 0.71124915 | 0.766 | 0.508 | 5.60E-201 | 0 |
| PDZRN3 | 4.32E-166 | 0.70604478 | 0.845 | 0.597 | 1.34E-161 | 0 |
| AGBL4 | 2.22E-125 | 0.69089767 | 0.753 | 0.551 | 6.93E-121 | 0 |
| PTPRQ | 2.75E-197 | 0.69049477 | 0.482 | 0.173 | 8.56E-193 | 0 |
| EYS | 5.85E-255 | 0.68564677 | 0.843 | 0.553 | 1.82E-250 | 0 |
| CCDC200 | 2.64E-294 | 2.77969171 | 0.342 | 0.065 | 8.22E-290 | 1 |
| SCD | 0 | 2.68502657 | 0.59 | 0.162 | 0 | 1 |
| TEX14 | 0 | 2.65644527 | 0.422 | 0.07 | 0 | 1 |
| PDE4D | 0 | 2.58621697 | 0.547 | 0.177 | 0 | 1 |
| F13A1 | 0 | 2.53756941 | 0.432 | 0.069 | 0 | 1 |
| DCN | 0 | 2.4477494 | 0.642 | 0.223 | 0 | 1 |
| NEGR1 | 0 | 2.30897987 | 0.591 | 0.222 | 0 | 1 |
| ABI3BP | 0 | 2.30004688 | 0.596 | 0.232 | 0 | 1 |
| JUND | 7.45E-176 | 2.24080122 | 0.577 | 0.35 | 2.32E-171 | 1 |
| FRMD4B | 0 | 2.22633503 | 0.414 | 0.086 | 0 | 1 |
| CROCC | 5.92E-50 | 2.18417001 | 0.388 | 0.282 | 1.84E-45 | 1 |
| FTL | 7.20E-174 | 2.10502392 | 0.542 | 0.302 | 2.24E-169 | 1 |
| DCLK1 | 0 | 2.09490494 | 0.507 | 0.157 | 0 | 1 |
| ELMO1 | 0 | 2.07879105 | 0.472 | 0.068 | 0 | 1 |
| C12orf57 | 3.57E-254 | 2.01837306 | 0.254 | 0.037 | 1.11E-249 | 1 |
| ACACA | 3.44E-119 | 1.90538334 | 0.69 | 0.648 | 1.07E-114 | 1 |
| FASN | 5.87E-55 | 1.87103336 | 0.407 | 0.304 | 1.83E-50 | 1 |
| ROBO2 | 0 | 1.85564454 | 0.322 | 0.05 | 0 | 1 |
| JUN | 1.04E-178 | 1.84570321 | 0.613 | 0.374 | 3.25E-174 | 1 |
| S100A6 | 3.07E-81 | 1.83768505 | 0.448 | 0.306 | 9.55E-77 | 1 |
| WDR74 | 9.12E-134 | 1.80896977 | 0.266 | 0.082 | 2.84E-129 | 1 |
| RBPJ | 1.72E-114 | 1.78974465 | 0.652 | 0.645 | 5.35E-110 | 1 |
| PSD3 | 2.39E-180 | 1.76628188 | 0.546 | 0.294 | 7.46E-176 | 1 |
| SELENOP | 7.34E-68 | 1.75216878 | 0.513 | 0.429 | 2.29E-63 | 1 |
| COLEC12 | 2.82E-284 | 1.73292213 | 0.343 | 0.068 | 8.77E-280 | 1 |
| CREB5 | 1.40E-279 | 1.68846883 | 0.342 | 0.068 | 4.38E-275 | 1 |
| MTSS1 | 2.02E-103 | 1.67134867 | 0.463 | 0.284 | 6.29E-99 | 1 |
| FTH1 | 1.01E-55 | 1.67061834 | 0.466 | 0.38 | 3.15E-51 | 1 |
| B2M | 3.31E-100 | 1.6623125 | 0.411 | 0.228 | 1.03E-95 | 1 |
| PRRX1 | 0 | 1.64368872 | 0.356 | 0.048 | 0 | 1 |
| ITGBL1 | 0 | 2.68888223 | 0.656 | 0.071 | 0 | 2 |
| ANO5 | 0 | 2.41921128 | 0.806 | 0.209 | 0 | 2 |
| GPC6 | 0 | 2.23701055 | 0.892 | 0.592 | 0 | 2 |
| STK32A | 0 | 2.19418432 | 0.866 | 0.167 | 0 | 2 |
| SEMA3C | 0 | 1.96550475 | 0.993 | 0.738 | 0 | 2 |
| KCNQ3 | 0 | 1.9267872 | 0.899 | 0.292 | 0 | 2 |
| GPR39 | 0 | 1.86498215 | 0.634 | 0.061 | 0 | 2 |
| SPAG17 | 0 | 1.8345231 | 0.905 | 0.52 | 0 | 2 |
| CPA6 | 0 | 1.81064697 | 0.591 | 0.067 | 0 | 2 |
| ABCC3 | 0 | 1.72743325 | 0.659 | 0.08 | 0 | 2 |
| KCND2 | 0 | 1.69262381 | 0.626 | 0.172 | 0 | 2 |
| XYLT1 | 0 | 1.68299627 | 0.967 | 0.602 | 0 | 2 |
| CAMK2D | 9.41E-305 | 1.6646019 | 0.961 | 0.741 | 2.93E-300 | 2 |
| ANKRD36C | 0 | 1.6389676 | 0.955 | 0.624 | 0 | 2 |
| PPP2R1B | 0 | 1.53211042 | 0.973 | 0.682 | 0 | 2 |
| AOX1 | 0 | 1.50551156 | 0.712 | 0.185 | 0 | 2 |
| GALNT13 | 0 | 1.48287303 | 0.596 | 0.135 | 0 | 2 |
| PPL | 0 | 1.39619491 | 0.687 | 0.102 | 0 | 2 |
| NTN4 | 0 | 1.38792962 | 0.497 | 0.082 | 0 | 2 |
| MYOF | 0 | 1.38364848 | 0.893 | 0.379 | 0 | 2 |
| SCN8A | 0 | 1.28226832 | 0.518 | 0.038 | 0 | 2 |
| ESRRG | 6.57E-210 | 1.23972649 | 0.799 | 0.425 | 2.05E-205 | 2 |
| PALLD | 1.67E-232 | 1.19403442 | 0.667 | 0.286 | 5.22E-228 | 2 |
| SRGAP1 | 2.61E-245 | 1.14888568 | 0.882 | 0.528 | 8.12E-241 | 2 |
| ADK | 2.66E-224 | 1.14563588 | 0.995 | 0.842 | 8.29E-220 | 2 |
| RALGPS2 | 0 | 1.13786661 | 0.631 | 0.131 | 0 | 2 |
| FAM126A | 3.95E-291 | 1.11725626 | 0.909 | 0.534 | 1.23E-286 | 2 |
| TSHZ2 | 3.69E-179 | 1.11225545 | 0.914 | 0.668 | 1.15E-174 | 2 |
| CAV1 | 7.32E-224 | 1.09693578 | 0.966 | 0.706 | 2.28E-219 | 2 |
| LAMA4 | 0 | 1.08789252 | 1 | 0.928 | 0 | 2 |
| ATP1B3 | 1.26E-100 | 3.3007941 | 0.779 | 0.316 | 3.94E-96 | 3 |
| CMSS1 | 5.42E-48 | 2.82881156 | 0.868 | 0.583 | 1.69E-43 | 3 |
| PGAP1 | 3.55E-72 | 2.76690303 | 0.68 | 0.279 | 1.11E-67 | 3 |
| PHLDB2 | 1.02E-91 | 2.26280553 | 0.978 | 0.862 | 3.18E-87 | 3 |
| CTH | 2.99E-104 | 2.17497161 | 0.515 | 0.11 | 9.31E-100 | 3 |
| CFAP69 | 8.94E-117 | 2.0015443 | 0.915 | 0.447 | 2.79E-112 | 3 |
| ULK4 | 2.21E-34 | 1.71837864 | 0.864 | 0.608 | 6.90E-30 | 3 |
| ACER2 | 2.48E-61 | 1.61662851 | 0.79 | 0.425 | 7.74E-57 | 3 |
| UAP1 | 6.64E-81 | 1.58307622 | 0.724 | 0.276 | 2.07E-76 | 3 |
| SAT1 | 2.25E-75 | 1.5091085 | 0.941 | 0.695 | 7.01E-71 | 3 |
| PDK4 | 3.02E-41 | 1.49809866 | 0.871 | 0.627 | 9.42E-37 | 3 |
| PLD1 | 1.26E-40 | 1.49799888 | 0.897 | 0.731 | 3.94E-36 | 3 |
| VGLL4 | 3.84E-57 | 1.43862881 | 0.897 | 0.667 | 1.20E-52 | 3 |
| LGR4 | 2.81E-31 | 1.3315722 | 0.879 | 0.741 | 8.77E-27 | 3 |
| MAP3K5 | 1.70E-53 | 1.32368114 | 0.824 | 0.511 | 5.31E-49 | 3 |
| TMOD1 | 5.13E-76 | 1.31022488 | 0.688 | 0.273 | 1.60E-71 | 3 |
| PC | 1.79E-65 | 1.29472007 | 0.949 | 0.665 | 5.58E-61 | 3 |
| ELL2 | 2.89E-26 | 1.29254722 | 0.945 | 0.797 | 8.99E-22 | 3 |
| PKP2 | 5.07E-41 | 1.29031911 | 0.614 | 0.316 | 1.58E-36 | 3 |
| FILIP1L | 5.59E-25 | 1.27568302 | 0.835 | 0.633 | 1.74E-20 | 3 |
| FYN | 1.05E-50 | 1.27082848 | 0.879 | 0.606 | 3.28E-46 | 3 |
| ACSL4 | 2.24E-28 | 1.24987464 | 0.702 | 0.43 | 6.97E-24 | 3 |
| FAM184A | 7.28E-48 | 1.235134 | 0.651 | 0.318 | 2.27E-43 | 3 |
| KSR1 | 1.21E-69 | 1.23384552 | 0.562 | 0.179 | 3.75E-65 | 3 |
| SLC19A2 | 8.57E-69 | 1.17261436 | 0.776 | 0.372 | 2.67E-64 | 3 |
| SH3BP5 | 1.08E-40 | 1.16348075 | 0.809 | 0.538 | 3.35E-36 | 3 |
| GPT2 | 7.21E-41 | 1.12927196 | 0.827 | 0.606 | 2.25E-36 | 3 |
| EPB41L3 | 3.08E-21 | 1.11933704 | 0.691 | 0.484 | 9.59E-17 | 3 |
| HIVEP2 | 6.09E-42 | 1.11447337 | 0.86 | 0.615 | 1.90E-37 | 3 |
| SCCPDH | 4.31E-51 | 1.11025304 | 0.739 | 0.402 | 1.34E-46 | 3 |

**Supplementary Table 2D: Endothelial cell subcluster transcriptomic profiles.** Top 30 unique transcriptomic markers of identified endothelial cell subclusters from n=21 IFPs. Genes are organized by decreasing average Log2 fold change (FC) filtered by adjusted p < 0.05, log2FC ≥ 0.5, and min.pct ≥ 0.25. Genes duplicated across multiple subclusters, unknown genes, and non-coding genes were not included.

| gene | p_val | avg_log2FC | pct.1 | pct.2 | p_val_adj | cluster |
|---|---|---|---|---|---|---|
| CADM2 | 2.00E-293 | 2.968101895 | 0.322 | 0.044 | 6.23E-289 | 0 |
| BTNL9 | 0 | 2.883778551 | 0.734 | 0.157 | 0 | 0 |
| CD36 | 0 | 2.534451945 | 0.703 | 0.314 | 0 | 0 |
| KIAA1217 | 0 | 1.782021244 | 0.812 | 0.612 | 0 | 0 |
| PPARG | 0 | 1.74017807 | 0.713 | 0.392 | 0 | 0 |
| ABLIM3 | 4.50E-98 | 1.705366492 | 0.593 | 0.466 | 1.40E-93 | 0 |
| LNX1 | 0 | 1.618130926 | 0.516 | 0.134 | 0 | 0 |
| CCDC85A | 1.41E-266 | 1.596114117 | 0.622 | 0.308 | 4.40E-262 | 0 |
| PRAG1 | 0 | 1.578891618 | 0.399 | 0.07 | 0 | 0 |
| ADAMTS18 | 2.23E-114 | 1.575969651 | 0.29 | 0.112 | 6.95E-110 | 0 |
| FABP4 | 8.00E-287 | 1.537617674 | 0.582 | 0.222 | 2.49E-282 | 0 |
| SLC6A6 | 1.97E-238 | 1.527600972 | 0.372 | 0.095 | 6.13E-234 | 0 |
| ADGRF5 | 0 | 1.363627465 | 0.684 | 0.271 | 0 | 0 |
| HDAC9 | 3.65E-169 | 1.329191832 | 0.524 | 0.26 | 1.14E-164 | 0 |
| PIK3R3 | 9.70E-159 | 1.316237622 | 0.679 | 0.468 | 3.02E-154 | 0 |
| DACH1 | 4.14E-232 | 1.304208041 | 0.698 | 0.389 | 1.29E-227 | 0 |
| DYSF | 2.33E-281 | 1.298959813 | 0.508 | 0.168 | 7.26E-277 | 0 |
| COL15A1 | 2.81E-237 | 1.272653239 | 0.639 | 0.309 | 8.74E-233 | 0 |
| KALRN | 1.66E-180 | 1.253628986 | 0.744 | 0.595 | 5.16E-176 | 0 |
| SPAAR | 0 | 1.25174832 | 0.391 | 0.048 | 0 | 0 |
| NAV1 | 2.65E-295 | 1.247301387 | 0.723 | 0.383 | 8.25E-291 | 0 |
| ZBTB46 | 3.17E-258 | 1.211635037 | 0.659 | 0.34 | 9.88E-254 | 0 |
| KDR | 6.35E-255 | 1.197827736 | 0.469 | 0.151 | 1.98E-250 | 0 |
| CD300LG | 0 | 1.196086839 | 0.392 | 0.022 | 0 | 0 |
| NOTCH4 | 0 | 1.195819564 | 0.639 | 0.236 | 0 | 0 |
| MGLL | 2.80E-177 | 1.187613334 | 0.61 | 0.343 | 8.74E-173 | 0 |
| NRG2 | 1.82E-261 | 1.183892151 | 0.31 | 0.045 | 5.68E-257 | 0 |
| NRP1 | 2.79E-243 | 1.170648799 | 0.785 | 0.554 | 8.69E-239 | 0 |
| HECW2 | 1.56E-175 | 1.162470355 | 0.4 | 0.144 | 4.87E-171 | 0 |
| MCC | 1.90E-223 | 1.149841402 | 0.605 | 0.299 | 5.93E-219 | 0 |
| PLCXD3 | 0 | 2.372562145 | 0.451 | 0.045 | 0 | 1 |
| TLL1 | 0 | 2.064909716 | 0.625 | 0.171 | 0 | 1 |
| GPM6A | 9.81E-206 | 1.957455921 | 0.28 | 0.054 | 3.06E-201 | 1 |
| ZNF521 | 0 | 1.776351902 | 0.932 | 0.525 | 0 | 1 |
| MYRIP | 0 | 1.640901215 | 0.86 | 0.461 | 0 | 1 |
| RAB3C | 0 | 1.587496951 | 0.368 | 0.037 | 0 | 1 |
| RYR3 | 3.07E-283 | 1.536638542 | 0.578 | 0.201 | 9.55E-279 | 1 |
| MMP16 | 6.74E-287 | 1.524259936 | 0.38 | 0.073 | 2.10E-282 | 1 |
| PGM5 | 0 | 1.519000023 | 0.709 | 0.283 | 0 | 1 |
| MMRN1 | 0 | 1.494615346 | 0.359 | 0.052 | 0 | 1 |
| PDIA5 | 0 | 1.433681736 | 0.678 | 0.183 | 0 | 1 |
| PRKAG2 | 0 | 1.410954425 | 0.786 | 0.3 | 0 | 1 |
| FRMPD4 | 3.48E-244 | 1.381938783 | 0.328 | 0.06 | 1.09E-239 | 1 |
| ST6GAL1 | 2.86E-272 | 1.299530462 | 0.781 | 0.436 | 8.90E-268 | 1 |
| PLXNA4 | 8.80E-263 | 1.296648153 | 0.374 | 0.076 | 2.74E-258 | 1 |
| SEMA6A | 5.32E-292 | 1.292845392 | 0.689 | 0.309 | 1.66E-287 | 1 |
| PTPRE | 0 | 1.283630333 | 0.774 | 0.374 | 0 | 1 |
| ALDH1A2 | 3.17E-262 | 1.24806462 | 0.446 | 0.115 | 9.86E-258 | 1 |
| SLCO2A1 | 4.99E-260 | 1.235690042 | 0.563 | 0.192 | 1.56E-255 | 1 |
| EPB41L3 | 4.80E-296 | 1.23160182 | 0.647 | 0.231 | 1.50E-291 | 1 |
| TGFBR3 | 1.15E-281 | 1.221764403 | 0.767 | 0.372 | 3.57E-277 | 1 |
| TRABD2B | 8.29E-292 | 1.203185102 | 0.315 | 0.039 | 2.58E-287 | 1 |
| PBX1 | 2.69E-244 | 1.178583831 | 0.731 | 0.371 | 8.37E-240 | 1 |
| DOCK11 | 0 | 1.161425454 | 0.623 | 0.194 | 0 | 1 |
| TPO | 3.29E-75 | 1.130209341 | 0.56 | 0.392 | 1.03E-70 | 1 |
| SELP | 0 | 1.126854452 | 0.418 | 0.038 | 0 | 1 |
| ENTPD1 | 2.06E-277 | 1.123838526 | 0.743 | 0.343 | 6.41E-273 | 1 |
| LYST | 1.49E-249 | 1.110249255 | 0.864 | 0.631 | 4.63E-245 | 1 |
| ZNF385D | 2.93E-170 | 1.106248784 | 0.718 | 0.424 | 9.11E-166 | 1 |
| PLEKHA7 | 2.68E-221 | 1.105975925 | 0.613 | 0.276 | 8.34E-217 | 1 |
| NR4A1 | 0 | 3.495952546 | 0.788 | 0.088 | 0 | 2 |
| UBC | 0 | 3.191393266 | 0.826 | 0.195 | 0 | 2 |
| ZFP36 | 0 | 3.155462999 | 0.779 | 0.108 | 0 | 2 |
| DEPP1 | 0 | 3.018886428 | 0.788 | 0.134 | 0 | 2 |
| ACTA2 | 0 | 2.824600061 | 0.497 | 0.044 | 0 | 2 |
| HSPB1 | 0 | 2.760145419 | 0.755 | 0.217 | 0 | 2 |
| JUNB | 0 | 2.714031033 | 0.69 | 0.124 | 0 | 2 |
| NDRG1 | 0 | 2.705911582 | 0.778 | 0.363 | 0 | 2 |
| DUSP1 | 0 | 2.687725019 | 0.754 | 0.149 | 0 | 2 |
| C11orf96 | 0 | 2.535184154 | 0.437 | 0.026 | 0 | 2 |
| IER2 | 0 | 2.337632886 | 0.587 | 0.171 | 0 | 2 |
| KLF2 | 0 | 2.289599382 | 0.798 | 0.382 | 0 | 2 |
| GADD45B | 0 | 2.278020171 | 0.446 | 0.046 | 0 | 2 |
| FOSL2 | 4.23E-306 | 2.202493836 | 0.523 | 0.152 | 1.32E-301 | 2 |
| RHOB | 0 | 2.184342732 | 0.556 | 0.15 | 0 | 2 |

| gene | p_val | avg_log2FC | pct.1 | pct.2 | p_val_adj | cluster |
|---|---|---|---|---|---|---|
| NFKBIA | 3.17E-252 | 2.169967716 | 0.527 | 0.19 | 9.88E-248 | 2 |
| EGR1 | 0 | 2.122236164 | 0.359 | 0.033 | 0 | 2 |
| MYL9 | 1.17E-251 | 2.055554108 | 0.457 | 0.13 | 3.63E-247 | 2 |
| INPP4B | 7.30E-212 | 2.021760165 | 0.406 | 0.117 | 2.27E-207 | 2 |
| CCN1 | 0 | 2.007912728 | 0.436 | 0.083 | 0 | 2 |
| SOCS3 | 0 | 2.002701574 | 0.357 | 0.036 | 0 | 2 |
| TPM4 | 3.53E-239 | 1.989796241 | 0.618 | 0.312 | 1.10E-234 | 2 |
| MT2A | 1.33E-277 | 1.977703296 | 0.411 | 0.085 | 4.13E-273 | 2 |
| ADGRL3 | 2.23E-301 | 1.969733219 | 0.296 | 0.029 | 6.93E-297 | 2 |
| GRASP | 2.35E-195 | 1.951705329 | 0.546 | 0.264 | 7.33E-191 | 2 |
| PDLIM1 | 2.27E-209 | 1.939688064 | 0.58 | 0.301 | 7.06E-205 | 2 |
| SELE | 0 | 1.937781407 | 0.316 | 0.033 | 0 | 2 |
| IVNS1ABP | 1.83E-173 | 1.932191616 | 0.458 | 0.189 | 5.71E-169 | 2 |
| NEDD9 | 4.71E-287 | 1.905115386 | 0.709 | 0.386 | 1.47E-282 | 2 |
| EEF1A1 | 3.33E-244 | 1.869294571 | 0.642 | 0.315 | 1.04E-239 | 2 |
| NKAIN2 | 0 | 4.372038723 | 0.858 | 0.099 | 0 | 3 |
| SULF1 | 0 | 3.030508537 | 0.929 | 0.25 | 0 | 3 |
| ARL15 | 0 | 3.007707776 | 0.994 | 0.802 | 0 | 3 |
| PCSK5 | 0 | 2.965280907 | 0.877 | 0.22 | 0 | 3 |
| TOX | 0 | 2.559629889 | 0.606 | 0.027 | 0 | 3 |
| HTR4 | 0 | 2.39486833 | 0.547 | 0.011 | 0 | 3 |
| CHRM3 | 0 | 2.265291908 | 0.688 | 0.112 | 0 | 3 |
| VEGFC | 0 | 2.258646215 | 0.858 | 0.289 | 0 | 3 |
| NEBL | 0 | 2.132316306 | 0.719 | 0.152 | 0 | 3 |
| PCDH11X | 0 | 2.119751617 | 0.351 | 0.018 | 0 | 3 |
| COL8A1 | 0 | 2.081761546 | 0.594 | 0.093 | 0 | 3 |
| GUCY1A1 | 0 | 2.056426953 | 0.67 | 0.105 | 0 | 3 |
| SYT1 | 0 | 2.014885132 | 0.538 | 0.088 | 0 | 3 |
| PRDM16 | 0 | 1.959577885 | 0.765 | 0.102 | 0 | 3 |
| SH3RF2 | 0 | 1.943173459 | 0.604 | 0.058 | 0 | 3 |
| ITPR2 | 0 | 1.94289583 | 0.959 | 0.606 | 0 | 3 |
| IGFBP3 | 0 | 1.931505945 | 0.535 | 0.069 | 0 | 3 |
| PLPP1 | 4.00E-173 | 1.858391933 | 0.806 | 0.454 | 1.25E-168 | 3 |
| ELN | 2.14E-238 | 1.82298546 | 0.753 | 0.298 | 6.66E-234 | 3 |
| ATP2A3 | 0 | 1.81403515 | 0.659 | 0.104 | 0 | 3 |
| NOS1AP | 0 | 1.809927159 | 0.729 | 0.162 | 0 | 3 |
| EPHA3 | 0 | 1.757323333 | 0.471 | 0.036 | 0 | 3 |
| DKK2 | 3.79E-307 | 1.724600868 | 0.256 | 0.012 | 1.18E-302 | 3 |
| GJA5 | 0 | 1.650312123 | 0.46 | 0.016 | 0 | 3 |
| PDGFD | 1.35E-249 | 1.516375858 | 0.599 | 0.151 | 4.21E-245 | 3 |
| COL21A1 | 6.46E-223 | 1.48358828 | 0.581 | 0.158 | 2.01E-218 | 3 |
| WIPF3 | 0 | 1.461529634 | 0.44 | 0.021 | 0 | 3 |
| MGP | 6.63E-188 | 1.453227713 | 0.517 | 0.138 | 2.06E-183 | 3 |
| JAG1 | 1.17E-230 | 1.436154303 | 0.723 | 0.268 | 3.65E-226 | 3 |
| PTPRG | 1.11E-239 | 1.422431011 | 0.935 | 0.59 | 3.45E-235 | 3 |
| NOVA1 | 8.71E-226 | 2.868928805 | 0.772 | 0.169 | 2.71E-221 | 4 |
| ABI3BP | 1.52E-195 | 2.251755739 | 0.735 | 0.169 | 4.72E-191 | 4 |
| DCN | 1.93E-217 | 2.226035757 | 0.639 | 0.106 | 6.00E-213 | 4 |
| DCLK1 | 2.53E-198 | 2.197367458 | 0.631 | 0.112 | 7.88E-194 | 4 |
| BICC1 | 4.11E-222 | 2.11024733 | 0.475 | 0.053 | 1.28E-217 | 4 |
| KAZN | 5.69E-78 | 2.086468045 | 0.584 | 0.22 | 1.77E-73 | 4 |
| ITGBL1 | 3.79E-139 | 1.790616371 | 0.363 | 0.049 | 1.18E-134 | 4 |
| ANK2 | 2.89E-212 | 1.721658061 | 0.52 | 0.066 | 9.02E-208 | 4 |
| COL1A2 | 2.77E-173 | 1.687508 | 0.57 | 0.098 | 8.62E-169 | 4 |
| PID1 | 4.72E-191 | 1.681045569 | 0.395 | 0.041 | 1.47E-186 | 4 |
| ZNF385B | 4.79E-105 | 1.663901327 | 0.44 | 0.092 | 1.49E-100 | 4 |
| KCND2 | 3.08E-138 | 1.588108751 | 0.268 | 0.026 | 9.61E-134 | 4 |
| ABCA6 | 1.76E-153 | 1.585209318 | 0.464 | 0.072 | 5.48E-149 | 4 |
| ADH1B | 1.45E-86 | 1.578413025 | 0.366 | 0.075 | 4.53E-82 | 4 |
| CFD | 8.37E-102 | 1.570060105 | 0.469 | 0.105 | 2.61E-97 | 4 |
| DDR2 | 7.38E-187 | 1.561753973 | 0.493 | 0.067 | 2.30E-182 | 4 |
| NOX4 | 1.26E-60 | 1.548472925 | 0.39 | 0.114 | 3.94E-56 | 4 |
| TPRG1 | 1.72E-149 | 1.525283415 | 0.464 | 0.073 | 5.35E-145 | 4 |
| TRPS1 | 3.00E-87 | 1.524272617 | 0.621 | 0.216 | 9.34E-83 | 4 |
| ZEB2 | 3.03E-102 | 1.51478301 | 0.584 | 0.165 | 9.44E-98 | 4 |
| SEMA3C | 1.05E-120 | 1.513960229 | 0.427 | 0.079 | 3.26E-116 | 4 |
| LRP1B | 5.29E-46 | 1.506898943 | 0.424 | 0.158 | 1.65E-41 | 4 |
| LPAR1 | 6.94E-281 | 1.490948656 | 0.464 | 0.036 | 2.16E-276 | 4 |
| ABCA8 | 3.90E-167 | 1.463165248 | 0.408 | 0.05 | 1.21E-162 | 4 |
| PCDH9 | 5.23E-94 | 1.456067373 | 0.435 | 0.1 | 1.63E-89 | 4 |
| MGST1 | 5.78E-126 | 1.443608984 | 0.332 | 0.044 | 1.80E-121 | 4 |
| CCN5 | 3.56E-149 | 1.430447609 | 0.361 | 0.043 | 1.11E-144 | 4 |
| GPC6 | 1.25E-81 | 1.414827389 | 0.292 | 0.053 | 3.89E-77 | 4 |
| UST | 3.17E-248 | 1.414361068 | 0.414 | 0.033 | 9.89E-244 | 4 |
| PDGFRA | 2.73E-260 | 1.401296147 | 0.393 | 0.027 | 8.51E-256 | 4 |
