## Supplementary Table 3 for "Cell and Transcriptomic Diversity of Infrapatellar Fat Pad during Knee Osteoarthritis"

**Supplementary Table 3: CellChat putative cell-cell communications.** Inferred cell-cell communications at the level of ligands (source column is set to macrophages, adipocytes and endothelial cells) and receptors (target column set to fibroblasts) using CellChat software v2.

| | source | target | ligand | receptor | prob | pval | interaction_name | interaction_name_2 | pathway_name | annotation | evidence |
|---|---|---|---|---|---|---|---|---|---|---|---|
| 1 | Adipocytes | Fibroblasts | BMP5 | ACVR1_ACVR2A | 0.00013687 | 0 | BMP5_ACVR1_ACVR2A | BMP5 - (ACVR1+ACVR2A) | BMP | Secreted Signaling | KEGG: hsa04350; PMID:26893264 |
| 2 | Adipocytes | Fibroblasts | BMP5 | ACVR1_BMPR2 | 0.00024991 | 0 | BMP5_ACVR1_BMPR2 | BMP5 - (ACVR1+BMPR2) | BMP | Secreted Signaling | KEGG: hsa04350; PMID:26893264 |
| 3 | Adipocytes | Fibroblasts | BMP5 | BMPR1A_ACVR2A | 0.00024518 | 0 | BMP5_BMPR1A_ACVR2A | BMP5 - (BMPR1A+ACVR2A) | BMP | Secreted Signaling | KEGG: hsa04350; PMID:26893264 |
| 4 | Adipocytes | Fibroblasts | BMP5 | BMPR1A_BMPR2 | 0.00044719 | 0 | BMP5_BMPR1A_BMPR2 | BMP5 - (BMPR1A+BMPR2) | BMP | Secreted Signaling | KEGG: hsa04350; PMID:26893264 |
| 5 | Endothelial | Fibroblasts | BMP6 | ACVR1_ACVR2A | 0.00026028 | 0 | BMP6_ACVR1_ACVR2A | BMP6 - (ACVR1+ACVR2A) | BMP | Secreted Signaling | KEGG: hsa04350; PMID:26893264 |
| 6 | Endothelial | Fibroblasts | BMP6 | ACVR1_BMPR2 | 0.00047433 | 0 | BMP6_ACVR1_BMPR2 | BMP6 - (ACVR1+BMPR2) | BMP | Secreted Signaling | KEGG: hsa04350; PMID:26893264 |
| 7 | Endothelial | Fibroblasts | BMP6 | BMPR1A_ACVR2A | 0.00046539 | 0 | BMP6_BMPR1A_ACVR2A | BMP6 - (BMPR1A+ACVR2A) | BMP | Secreted Signaling | KEGG: hsa04350; PMID:26893264 |
| 8 | Endothelial | Fibroblasts | BMP6 | BMPR1A_BMPR2 | 0.00084594 | 0 | BMP6_BMPR1A_BMPR2 | BMP6 - (BMPR1A+BMPR2) | BMP | Secreted Signaling | KEGG: hsa04350; PMID:26893264 |
| 9 | Macrophages | Fibroblasts | WNT2B | FZD4_LRP6 | 0.00012895 | 0 | WNT2B_FZD4_LRP6 | WNT2B - (FZD4+LRP6) | WNT | Secreted Signaling | KEGG: hsa04310; PMID: 23209147 |
| 10 | Adipocytes | Fibroblasts | FGF1 | FGFR1 | 0.00072379 | 0 | FGF1_FGFR1 | FGF1 - FGFR1 | FGF | Secreted Signaling | PMC: 4393358 |
| 11 | Adipocytes | Fibroblasts | FGF2 | FGFR1 | 0.00289201 | 0 | FGF2_FGFR1 | FGF2 - FGFR1 | FGF | Secreted Signaling | PMC: 4393358 |
| 12 | Macrophages | Fibroblasts | PDGFB | PDGFRA | 0.00083101 | 0 | PDGFB_PDGFRA | PDGFB - PDGFRA | PDGF | Secreted Signaling | PMID: 15207812 |
| 13 | Endothelial | Fibroblasts | PDGFB | PDGFRA | 0.00070834 | 0 | PDGFB_PDGFRA | PDGFB - PDGFRA | PDGF | Secreted Signaling | PMID: 15207812 |
| 14 | Macrophages | Fibroblasts | PDGFB | PDGFRB | 0.00047672 | 0 | PDGFB_PDGFRB | PDGFB - PDGFRB | PDGF | Secreted Signaling | PMID: 15207812 |
| 15 | Endothelial | Fibroblasts | PDGFB | PDGFRB | 0.0004069 | 0 | PDGFB_PDGFRB | PDGFB - PDGFRB | PDGF | Secreted Signaling | PMID: 15207812 |
| 16 | Macrophages | Fibroblasts | PDGFC | PDGFRA | 0.00884738 | 0 | PDGFC_PDGFRA | PDGFC - PDGFRA | PDGF | Secreted Signaling | PMID: 15207812 |
| 17 | Adipocytes | Fibroblasts | PDGFC | PDGFRA | 0.00284928 | 0 | PDGFC_PDGFRA | PDGFC - PDGFRA | PDGF | Secreted Signaling | PMID: 15207812 |
| 18 | Adipocytes | Fibroblasts | PDGFD | PDGFRB | 0.00049412 | 0 | PDGFD_PDGFRB | PDGFD - PDGFRB | PDGF | Secreted Signaling | PMID: 15207812 |
| 19 | Macrophages | Fibroblasts | IGF1 | IGF1R | 0.00132658 | 0 | IGF1_IGF1R | IGF1 - IGF1R | IGF | Secreted Signaling | PMID: 14604834 |
| 20 | Endothelial | Fibroblasts | CXCL12 | ACKR3 | 0.00012319 | 0 | CXCL12_ACKR3 | CXCL12 - ACKR3 | CXCL | Secreted Signaling | KEGG: hsa04060 |
| 21 | Adipocytes | Fibroblasts | ADIPOQ | ADIPOR2 | 0.00049885 | 0 | ADIPOQ_ADIPOR2 | ADIPOQ - ADIPOR2 | ADIPONECTIN | Secreted Signaling | PMID: 21284979 |
| 22 | Adipocytes | Fibroblasts | ANGPTL4 | CDH11 | 0.0003761 | 0 | ANGPTL4_CDH11 | ANGPTL4 - CDH11 | ANGPTL | Secreted Signaling | PMID: 30049845 |
| 23 | Adipocytes | Fibroblasts | ANGPTL4 | SDC2 | 0.00052652 | 0 | ANGPTL4_SDC2 | ANGPTL4 - SDC2 | ANGPTL | Secreted Signaling | PMID: 29017031 |
| 24 | Macrophages | Fibroblasts | GAS6 | AXL | 0.00040239 | 0 | GAS6_AXL | GAS6 - AXL | GAS | Secreted Signaling | PMID: 27801848 |
| 25 | Adipocytes | Fibroblasts | COL4A1 | ITGA9_ITGB1 | 0.00125192 | 0 | COL4A1_ITGA9_ITGB1 | COL4A1 - (ITGA9+ITGB1) | COLLAGEN | ECM-Receptor | KEGG: hsa04512 |
| 26 | Endothelial | Fibroblasts | COL4A1 | ITGA9_ITGB1 | 0.00028185 | 0 | COL4A1_ITGA9_ITGB1 | COL4A1 - (ITGA9+ITGB1) | COLLAGEN | ECM-Receptor | KEGG: hsa04512 |
| 27 | Adipocytes | Fibroblasts | COL4A2 | ITGA9_ITGB1 | 0.00128854 | 0 | COL4A2_ITGA9_ITGB1 | COL4A2 - (ITGA9+ITGB1) | COLLAGEN | ECM-Receptor | KEGG: hsa04512 |
| 28 | Endothelial | Fibroblasts | COL4A2 | ITGA9_ITGB1 | 0.00063678 | 0 | COL4A2_ITGA9_ITGB1 | COL4A2 - (ITGA9+ITGB1) | COLLAGEN | ECM-Receptor | KEGG: hsa04512 |
| 29 | Adipocytes | Fibroblasts | COL6A1 | ITGA9_ITGB1 | 0.00029747 | 0 | COL6A1_ITGA9_ITGB1 | COL6A1 - (ITGA9+ITGB1) | COLLAGEN | ECM-Receptor | KEGG: hsa04512 |
| 30 | Endothelial | Fibroblasts | LAMA3 | ITGA9_ITGB1 | 0.00040963 | 0 | LAMA3_ITGA9_ITGB1 | LAMA3 - (ITGA9+ITGB1) | LAMININ | ECM-Receptor | KEGG: hsa04512 |
| 31 | Adipocytes | Fibroblasts | LAMA4 | ITGA9_ITGB1 | 0.0022216 | 0 | LAMA4_ITGA9_ITGB1 | LAMA4 - (ITGA9+ITGB1) | LAMININ | ECM-Receptor | KEGG: hsa04512 |
| 32 | Endothelial | Fibroblasts | LAMA5 | ITGA9_ITGB1 | 9.10E-05 | 0 | LAMA5_ITGA9_ITGB1 | LAMA5 - (ITGA9+ITGB1) | LAMININ | ECM-Receptor | KEGG: hsa04512 |
| 33 | Adipocytes | Fibroblasts | LAMB1 | ITGA9_ITGB1 | 0.00014273 | 0 | LAMB1_ITGA9_ITGB1 | LAMB1 - (ITGA9+ITGB1) | LAMININ | ECM-Receptor | KEGG: hsa04512 |
| 34 | Endothelial | Fibroblasts | LAMB1 | ITGA9_ITGB1 | 0.00015335 | 0 | LAMB1_ITGA9_ITGB1 | LAMB1 - (ITGA9+ITGB1) | LAMININ | ECM-Receptor | KEGG: hsa04512 |
| 35 | Adipocytes | Fibroblasts | LAMC1 | ITGA9_ITGB1 | 0.0006628 | 0 | LAMC1_ITGA9_ITGB1 | LAMC1 - (ITGA9+ITGB1) | LAMININ | ECM-Receptor | KEGG: hsa04512 |
| 36 | Adipocytes | Fibroblasts | COL1A2 | ITGA11_ITGB1 | 0.00012021 | 0 | COL1A2_ITGA11_ITGB1 | COL1A2 - (ITGA11+ITGB1) | COLLAGEN | ECM-Receptor | KEGG: hsa04512 |
| 37 | Adipocytes | Fibroblasts | COL4A1 | ITGA11_ITGB1 | 0.00194508 | 0 | COL4A1_ITGA11_ITGB1 | COL4A1 - (ITGA11+ITGB1) | COLLAGEN | ECM-Receptor | KEGG: hsa04512 |
| 38 | Endothelial | Fibroblasts | COL4A1 | ITGA11_ITGB1 | 0.00044051 | 0 | COL4A1_ITGA11_ITGB1 | COL4A1 - (ITGA11+ITGB1) | COLLAGEN | ECM-Receptor | KEGG: hsa04512 |
| 39 | Adipocytes | Fibroblasts | COL4A2 | ITGA11_ITGB1 | 0.00200148 | 0 | COL4A2_ITGA11_ITGB1 | COL4A2 - (ITGA11+ITGB1) | COLLAGEN | ECM-Receptor | KEGG: hsa04512 |
| 40 | Endothelial | Fibroblasts | COL4A2 | ITGA11_ITGB1 | 0.00099222 | 0 | COL4A2_ITGA11_ITGB1 | COL4A2 - (ITGA11+ITGB1) | COLLAGEN | ECM-Receptor | KEGG: hsa04512 |
| 41 | Adipocytes | Fibroblasts | COL6A1 | ITGA11_ITGB1 | 0.00046513 | 0 | COL6A1_ITGA11_ITGB1 | COL6A1 - (ITGA11+ITGB1) | COLLAGEN | ECM-Receptor | KEGG: hsa04512 |
| 42 | Adipocytes | Fibroblasts | COL6A2 | ITGA11_ITGB1 | 0.00019709 | 0 | COL6A2_ITGA11_ITGB1 | COL6A2 - (ITGA11+ITGB1) | COLLAGEN | ECM-Receptor | KEGG: hsa04512 |
| 43 | Adipocytes | Fibroblasts | COL6A3 | ITGA11_ITGB1 | 0.0001976 | 0 | COL6A3_ITGA11_ITGB1 | COL6A3 - (ITGA11+ITGB1) | COLLAGEN | ECM-Receptor | KEGG: hsa04512 |
| 44 | Macrophages | Fibroblasts | FN1 | ITGAV_ITGB8 | 0.00071038 | 0 | FN1_ITGAV_ITGB8 | FN1 - (ITGAV+ITGB8) | FN1 | ECM-Receptor | KEGG: hsa04512 |
| 45 | Adipocytes | Fibroblasts | FN1 | ITGAV_ITGB8 | 0.00011943 | 0 | FN1_ITGAV_ITGB8 | FN1 - (ITGAV+ITGB8) | FN1 | ECM-Receptor | KEGG: hsa04512 |
| 46 | Endothelial | Fibroblasts | FN1 | ITGAV_ITGB8 | 0.00046835 | 0 | FN1_ITGAV_ITGB8 | FN1 - (ITGAV+ITGB8) | FN1 | ECM-Receptor | KEGG: hsa04512 |
| 47 | Adipocytes | Fibroblasts | LAMA2 | ITGAV_ITGB8 | 0.00097368 | 0 | LAMA2_ITGAV_ITGB8 | LAMA2 - (ITGAV+ITGB8) | LAMININ | ECM-Receptor | KEGG: hsa04512 |
| 48 | Endothelial | Fibroblasts | LAMA3 | ITGAV_ITGB8 | 0.00070713 | 0 | LAMA3_ITGAV_ITGB8 | LAMA3 - (ITGAV+ITGB8) | LAMININ | ECM-Receptor | KEGG: hsa04512 |
| 49 | Adipocytes | Fibroblasts | LAMA4 | ITGAV_ITGB8 | 0.00377946 | 0 | LAMA4_ITGAV_ITGB8 | LAMA4 - (ITGAV+ITGB8) | LAMININ | ECM-Receptor | KEGG: hsa04512 |
| 50 | Endothelial | Fibroblasts | LAMA4 | ITGAV_ITGB8 | 0.00046889 | 0 | LAMA4_ITGAV_ITGB8 | LAMA4 - (ITGAV+ITGB8) | LAMININ | ECM-Receptor | KEGG: hsa04512 |
| 51 | Endothelial | Fibroblasts | LAMA5 | ITGAV_ITGB8 | 0.00015757 | 0 | LAMA5_ITGAV_ITGB8 | LAMA5 - (ITGAV+ITGB8) | LAMININ | ECM-Receptor | KEGG: hsa04512 |
| 52 | Adipocytes | Fibroblasts | LAMB1 | ITGAV_ITGB8 | 0.00024721 | 0 | LAMB1_ITGAV_ITGB8 | LAMB1 - (ITGAV+ITGB8) | LAMININ | ECM-Receptor | KEGG: hsa04512 |
| 53 | Endothelial | Fibroblasts | LAMB1 | ITGAV_ITGB8 | 0.00026547 | 0 | LAMB1_ITGAV_ITGB8 | LAMB1 - (ITGAV+ITGB8) | LAMININ | ECM-Receptor | KEGG: hsa04512 |
| 54 | Adipocytes | Fibroblasts | LAMC1 | ITGAV_ITGB8 | 0.0011428 | 0 | LAMC1_ITGAV_ITGB8 | LAMC1 - (ITGAV+ITGB8) | LAMININ | ECM-Receptor | KEGG: hsa04512 |
| 55 | Endothelial | Fibroblasts | LAMC1 | ITGAV_ITGB8 | 0.00042264 | 0 | LAMC1_ITGAV_ITGB8 | LAMC1 - (ITGAV+ITGB8) | LAMININ | ECM-Receptor | KEGG: hsa04512 |
| 56 | Adipocytes | Fibroblasts | COL1A2 | ITGAV_ITGB8 | 0.00013304 | 0 | COL1A2_ITGAV_ITGB8 | COL1A2 - (ITGAV+ITGB8) | COLLAGEN | ECM-Receptor | KEGG: hsa04512 |
| 57 | Adipocytes | Fibroblasts | COL4A1 | ITGAV_ITGB8 | 0.0021476 | 0 | COL4A1_ITGAV_ITGB8 | COL4A1 - (ITGAV+ITGB8) | COLLAGEN | ECM-Receptor | KEGG: hsa04512 |
| 58 | Endothelial | Fibroblasts | COL4A1 | ITGAV_ITGB8 | 0.00048723 | 0 | COL4A1_ITGAV_ITGB8 | COL4A1 - (ITGAV+ITGB8) | COLLAGEN | ECM-Receptor | KEGG: hsa04512 |
| 59 | Adipocytes | Fibroblasts | COL4A2 | ITGAV_ITGB8 | 0.00220972 | 0 | COL4A2_ITGAV_ITGB8 | COL4A2 - (ITGAV+ITGB8) | COLLAGEN | ECM-Receptor | KEGG: hsa04512 |
| 60 | Endothelial | Fibroblasts | COL4A2 | ITGAV_ITGB8 | 0.00109647 | 0 | COL4A2_ITGAV_ITGB8 | COL4A2 - (ITGAV+ITGB8) | COLLAGEN | ECM-Receptor | KEGG: hsa04512 |
| 61 | Adipocytes | Fibroblasts | COL6A1 | ITGAV_ITGB8 | 0.00051452 | 0 | COL6A1_ITGAV_ITGB8 | COL6A1 - (ITGAV+ITGB8) | COLLAGEN | ECM-Receptor | KEGG: hsa04512 |
| 62 | Adipocytes | Fibroblasts | COL6A2 | ITGAV_ITGB8 | 0.0002181 | 0 | COL6A2_ITGAV_ITGB8 | COL6A2 - (ITGAV+ITGB8) | COLLAGEN | ECM-Receptor | KEGG: hsa04512 |
| 63 | Adipocytes | Fibroblasts | COL6A3 | ITGAV_ITGB8 | 0.00021865 | 0 | COL6A3_ITGAV_ITGB8 | COL6A3 - (ITGAV+ITGB8) | COLLAGEN | ECM-Receptor | KEGG: hsa04512 |
| 64 | Macrophages | Fibroblasts | FN1 | CD44 | 0.00185942 | 0 | FN1_CD44 | FN1 - CD44 | FN1 | ECM-Receptor | KEGG: hsa04512 |
| 65 | Endothelial | Fibroblasts | FN1 | CD44 | 0.00122598 | 0 | FN1_CD44 | FN1 - CD44 | FN1 | ECM-Receptor | KEGG: hsa04512 |
| 66 | Adipocytes | Fibroblasts | COL4A1 | CD44 | 0.00545851 | 0 | COL4A1_CD44 | COL4A1 - CD44 | COLLAGEN | ECM-Receptor | KEGG: hsa04512 |
| 67 | Endothelial | Fibroblasts | COL4A1 | CD44 | 0.00127479 | 0 | COL4A1_CD44 | COL4A1 - CD44 | COLLAGEN | ECM-Receptor | KEGG: hsa04512 |
| 68 | Adipocytes | Fibroblasts | COL4A2 | CD44 | 0.00560985 | 0 | COL4A2_CD44 | COL4A2 - CD44 | COLLAGEN | ECM-Receptor | KEGG: hsa04512 |
| 69 | Endothelial | Fibroblasts | COL4A2 | CD44 | 0.00282618 | 0 | COL4A2_CD44 | COL4A2 - CD44 | COLLAGEN | ECM-Receptor | KEGG: hsa04512 |
| 70 | Adipocytes | Fibroblasts | COL6A1 | CD44 | 0.00134908 | 0 | COL6A1_CD44 | COL6A1 - CD44 | COLLAGEN | ECM-Receptor | KEGG: hsa04512 |
| 71 | Adipocytes | Fibroblasts | COL6A2 | CD44 | 0.00057515 | 0 | COL6A2_CD44 | COL6A2 - CD44 | COLLAGEN | ECM-Receptor | KEGG: hsa04512 |
| 72 | Adipocytes | Fibroblasts | COL6A3 | CD44 | 0.00057661 | 0 | COL6A3_CD44 | COL6A3 - CD44 | COLLAGEN | ECM-Receptor | KEGG: hsa04512 |
| 73 | Adipocytes | Fibroblasts | LAMA2 | CD44 | 0.00253051 | 0 | LAMA2_CD44 | LAMA2 - CD44 | LAMININ | ECM-Receptor | KEGG: hsa04512 |
| 74 | Endothelial | Fibroblasts | LAMA3 | CD44 | 0.00184012 | 0 | LAMA3_CD44 | LAMA3 - CD44 | LAMININ | ECM-Receptor | KEGG: hsa04512 |

| | | | | | | | | | | | |
|---|---|---|---|---|---|---|---|---|---|---|---|
| 75 | Adipocytes | Fibroblasts | LAMA4 | CD44 | 0.00932082 | 0 | LAMA4_CD44 | LAMA4 - CD44 | LAMININ | ECM-Receptor | KEGG: hsa04512 |
| 76 | Endothelial | Fibroblasts | LAMA5 | CD44 | 0.00041567 | 0 | LAMA5_CD44 | LAMA5 - CD44 | LAMININ | ECM-Receptor | KEGG: hsa04512 |
| 77 | Adipocytes | Fibroblasts | LAMB1 | CD44 | 0.00065157 | 0 | LAMB1_CD44 | LAMB1 - CD44 | LAMININ | ECM-Receptor | KEGG: hsa04512 |
| 78 | Endothelial | Fibroblasts | LAMB1 | CD44 | 0.00069842 | 0 | LAMB1_CD44 | LAMB1 - CD44 | LAMININ | ECM-Receptor | KEGG: hsa04512 |
| 79 | Adipocytes | Fibroblasts | LAMC1 | CD44 | 0.00296044 | 0 | LAMC1_CD44 | LAMC1 - CD44 | LAMININ | ECM-Receptor | KEGG: hsa04512 |
| 80 | Endothelial | Fibroblasts | LAMC1 | CD44 | 0.00110757 | 0 | LAMC1_CD44 | LAMC1 - CD44 | LAMININ | ECM-Receptor | KEGG: hsa04512 |
| 81 | Macrophages | Fibroblasts | ADGRE5 | CD55 | 0.00024759 | 0 | ADGRE5_CD55 | ADGRE5 - CD55 | ADGRE5 | Cell-Cell Contact | PMID: 11297558 |
| 82 | Macrophages | Fibroblasts | CD46 | JAG1 | 0.00027918 | 0 | CD46_JAG1 | CD46 - JAG1 | CD46 | Cell-Cell Contact | PMID: 23086448 |
| 83 | Adipocytes | Fibroblasts | CD46 | JAG1 | 0.00059526 | 0 | CD46_JAG1 | CD46 - JAG1 | CD46 | Cell-Cell Contact | PMID: 23086448 |
| 84 | Endothelial | Fibroblasts | CD46 | JAG1 | 0.00010881 | 0 | CD46_JAG1 | CD46 - JAG1 | CD46 | Cell-Cell Contact | PMID: 23086448 |
| 85 | Macrophages | Fibroblasts | CD99 | CD99 | 0.00135154 | 0 | CD99_CD99 | CD99 - CD99 | CD99 | Cell-Cell Contact | KEGG: hsa04514 |
| 86 | Adipocytes | Fibroblasts | CD99 | CD99 | 0.00093965 | 0 | CD99_CD99 | CD99 - CD99 | CD99 | Cell-Cell Contact | KEGG: hsa04514 |
| 87 | Endothelial | Fibroblasts | CD99 | CD99 | 0.00045578 | 0 | CD99_CD99 | CD99 - CD99 | CD99 | Cell-Cell Contact | KEGG: hsa04514 |
| 88 | Endothelial | Fibroblasts | JAM2 | ITGAV_ITGB1 | 0.00031178 | 0 | JAM2_ITGAV_ITGB1 | JAM2 - (ITGAV+ITGB1) | JAM | Cell-Cell Contact | KEGG: hsa04514 |
| 89 | Endothelial | Fibroblasts | JAM2 | JAM3 | 0.00010494 | 0 | JAM2_JAM3 | JAM2 - JAM3 | JAM | Cell-Cell Contact | PMC1237098; PMID: 11590146 |
| 90 | Adipocytes | Fibroblasts | F11R | JAM3 | 4.41E-05 | 0 | F11R_JAM3 | F11R - JAM3 | JAM | Cell-Cell Contact | KEGG: hsa04514 |
| 91 | Adipocytes | Fibroblasts | MPZL1 | MPZL1 | 0.00018352 | 0 | MPZL1_MPZL1 | MPZL1 - MPZL1 | MPZ | Cell-Cell Contact | KEGG: hsa04514 |
| 92 | Adipocytes | Fibroblasts | JAG1 | NOTCH2 | 0.00028865 | 0 | JAG1_NOTCH2 | JAG1 - NOTCH2 | NOTCH | Cell-Cell Contact | PMID: 22353464 |
| 93 | Endothelial | Fibroblasts | JAG1 | NOTCH2 | 0.0004096 | 0 | JAG1_NOTCH2 | JAG1 - NOTCH2 | NOTCH | Cell-Cell Contact | PMID: 22353464 |
| 94 | Adipocytes | Fibroblasts | NRXN3 | NLGN1 | 0.00020557 | 0 | NRXN3_NLGN1 | NRXN3 - NLGN1 | NRXN | Cell-Cell Contact | KEGG: hsa04514 |
| 95 | Endothelial | Fibroblasts | PTPRM | PTPRM | 0.00931703 | 0 | PTPRM_PTPRM | PTPRM - PTPRM | PTPRM | Cell-Cell Contact | KEGG: hsa04514 |
| 96 | Adipocytes | Fibroblasts | TENM3 | ADGRL2 | 0.00139766 | 0 | TENM3_ADGRL2 | TENM3 - ADGRL2 | ADGRL | Cell-Cell Contact | PMID: 25728924;PMID: 34022279;PMID: 27091502;PMID: 35907405 |
| 97 | Adipocytes | Fibroblasts | TENM4 | ADGRL2 | 0.00065863 | 0 | TENM4_ADGRL2 | TENM4 - ADGRL2 | ADGRL | Cell-Cell Contact | PMID: 25728924;PMID: 34022279;PMID: 27091502;PMID: 35907405 |
| 98 | Adipocytes | Fibroblasts | FLRT2 | FLRT2 | 0.00024521 | 0 | FLRT2_FLRT2 | FLRT2 - FLRT2 | FLRT | Cell-Cell Contact | PMID: 27091502 |
| 99 | Endothelial | Fibroblasts | FLRT2 | FLRT2 | 0.00036212 | 0 | FLRT2_FLRT2 | FLRT2 - FLRT2 | FLRT | Cell-Cell Contact | PMID: 27091502 |
| 100 | Adipocytes | Fibroblasts | FLRT2 | UNC5C | 0.0001232 | 0 | FLRT2_UNC5C | FLRT2 - UNC5C | UNC5 | Cell-Cell Contact | PMID: 27091502 |
| 101 | Endothelial | Fibroblasts | FLRT2 | UNC5C | 0.00018218 | 0 | FLRT2_UNC5C | FLRT2 - UNC5C | UNC5 | Cell-Cell Contact | PMID: 27091502 |
| 102 | Endothelial | Fibroblasts | NTN4 | NTRK2 | 0.00016574 | 0 | NTN4_NTRK2 | NTN4 - NTRK2 | Netrin | Cell-Cell Contact | PMID: 34531526 |
| 103 | Adipocytes | Fibroblasts | PGD2-AKR1C3 | PTGFR | 0.00020848 | 0 | PGF2a-AKR1C3_PTGFR | PGF2a-AKR1C3 - PTGFR | Prostaglandin | Non-protein Signaling | PMID: 34949672;PMID: 21508345 |
| 104 | Macrophages | Fibroblasts | Cholesterol-LIPA | RORA | 0.00181811 | 0 | Cholesterol-Cholesterol-LIPA_RORA | CHOLESTEROL-LIPA - RORA | Cholesterol | Non-protein Signaling | HMRbase;PMID:12467577;PDB:1N83 |
| 105 | Macrophages | Fibroblasts | DHEA-STS | ESR1 | 0.00039563 | 0 | Dehydroepiandrosterone-DHEA-STS_ESR1 | DHEA-STS - ESR1 | DHEA | Non-protein Signaling | HMRbase |
| 106 | Adipocytes | Fibroblasts | DHEAS-SULT2B | PPARG | 0.00013341 | 0 | DHEAsulfate-DHEAS-SULT2B_PPARG | DHEAS-SULT2B - PPARG | DHEAS | Non-protein Signaling | HMRbase |
