## Supplementary Table 4 for "Cell and Transcriptomic Diversity of Infrapatellar Fat Pad during Knee Osteoarthritis"

**Supplementary Table 4A: Fibroblast differentially expressed genes in knee OA (n=15) compared to healthy control donor (n=6) IFPs.** Upregulated genes are arranged by decreasing Log2 fold change (FC) and downregulated genes are arranged by increasing Log2FC. Genes were filtered by adjusted p < 0.05, log2FC ≥ 0.5, min.pct ≥ 0.5.

**Upregulated Genes**

| gene | p_val | avg_log2FC | pct.1 | pct.2 | p_val_adj |
|---|---|---|---|---|---|
| FN1 | 6.76E-261 | 1.879153272 | 0.752 | 0.543 | 2.11E-256 |
| ZNF385B | 0 | 1.325557746 | 0.686 | 0.432 | 0 |
| PRG4 | 3.26E-149 | 1.251924266 | 0.647 | 0.483 | 1.02E-144 |
| ANKH | 5.30E-282 | 1.127908845 | 0.532 | 0.286 | 1.65E-277 |
| KAZN | 1.13E-198 | 0.919284241 | 0.783 | 0.573 | 3.51E-194 |
| MAPK10 | 0 | 0.888625904 | 0.636 | 0.305 | 0 |
| LRP1B | 1.51E-208 | 0.775655017 | 0.605 | 0.375 | 4.69E-204 |
| SOX5 | 3.09E-199 | 0.741318119 | 0.954 | 0.841 | 9.62E-195 |
| SNED1 | 0 | 0.704108703 | 0.879 | 0.617 | 0 |
| SH3RF1 | 2.24E-280 | 0.686494708 | 0.623 | 0.353 | 6.99E-276 |
| MRC2 | 0 | 0.660429366 | 0.668 | 0.361 | 0 |
| ZNF385D | 4.54E-183 | 0.632874972 | 0.5 | 0.285 | 1.42E-178 |
| CUX1 | 7.99E-249 | 0.624454013 | 0.665 | 0.409 | 2.49E-244 |
| C1GALT1 | 1.50E-214 | 0.572584896 | 0.535 | 0.302 | 4.68E-210 |
| ZBTB7C | 3.23E-153 | 0.559690647 | 0.579 | 0.381 | 1.00E-148 |
| COLEC12 | 9.94E-177 | 0.544303735 | 0.564 | 0.337 | 3.10E-172 |
| COL5A2 | 4.54E-208 | 0.536422978 | 0.792 | 0.545 | 1.41E-203 |
| PPP3CA | 3.44E-207 | 0.530020898 | 0.813 | 0.561 | 1.07E-202 |
| MAGI2 | 2.44E-167 | 0.505833224 | 0.837 | 0.654 | 7.61E-163 |
| GULP1 | 8.96E-182 | 0.503825846 | 0.561 | 0.334 | 2.79E-177 |

**Downregulated Genes**

| gene | p_val | avg_log2FC | pct.1 | pct.2 | p_val_adj |
|---|---|---|---|---|---|
| JUN | 0 | -2.29920565 | 0.21 | 0.515 | 0 |
| ABCA1 | 3.97E-121 | -1.016921317 | 0.54 | 0.593 | 1.24E-116 |
| VIM | 1.01E-83 | -0.93646036 | 0.677 | 0.682 | 3.16E-79 |
| GSN | 4.11E-153 | -0.849921549 | 0.774 | 0.798 | 1.28E-148 |
| SAMD4A | 1.13E-82 | -0.842603864 | 0.549 | 0.584 | 3.50E-78 |
| CCN5 | 1.69E-30 | -0.831192309 | 0.543 | 0.534 | 5.25E-26 |
| ABLIM1 | 4.39E-66 | -0.818114541 | 0.603 | 0.611 | 1.37E-61 |
| SASH1 | 3.44E-41 | -0.768594475 | 0.674 | 0.63 | 1.07E-36 |
| ANXA1 | 1.64E-16 | -0.689412349 | 0.523 | 0.507 | 5.09E-12 |
| RBPJ | 5.86E-124 | -0.671019907 | 0.736 | 0.738 | 1.83E-119 |
| CFD | 8.65E-39 | -0.663947494 | 0.637 | 0.634 | 2.69E-34 |
| RAB1A | 4.37E-12 | -0.616238872 | 0.616 | 0.554 | 1.36E-07 |
| COL6A3 | 1.98E-67 | -0.591872006 | 0.725 | 0.709 | 6.16E-63 |
| TGFBR3 | 1.56E-55 | -0.559103196 | 0.679 | 0.662 | 4.85E-51 |
| GHR | 3.64E-61 | -0.550852322 | 0.565 | 0.6 | 1.13E-56 |
| EHBP1 | 2.69E-89 | -0.549666209 | 0.725 | 0.716 | 8.38E-85 |
| CBLB | 6.53E-25 | -0.502330606 | 0.76 | 0.693 | 2.03E-20 |
| RTN4 | 9.70E-17 | -0.502103173 | 0.654 | 0.599 | 3.02E-12 |

**Supplementary Table 4B: PathDIP pathways enriched for the upregulated differentially expressed gene-set list from fibroblasts of knee OA (n=15) vs heathy control donor (n=6) IFPs.** Pathways were filtered by adjusted p < 0.05.

| Pathway Source | Pathway Name | p-value | q-value (FDR: BH-method) | q-value (Bonferroni) | Ratio | Query mapped | Pathway size | Type | Category |
|---|---|---|---|---|---|---|---|---|---|
| WikiPathways | Focal adhesion | 0.00070341 | 0.00439633 | 0.0351707 | 0.23 | 3 | 198 | Cellular processes and organization | Cellular community |
| WikiPathways | miRNA targets in ECM and membrane receptors | 0.00069384 | 0.004956 | 0.034692 | 0.15 | 2 | 43 | Cellular processes and organization | Cellular community |
| Panther_Pathway | Integrin signalling pathway | 0.00041304 | 0.005163 | 0.020652 | 0.23 | 3 | 165 | Environmental information processing | Signal transduction |
| REACTOME | MET promotes cell motility | 0.00063072 | 0.00525601 | 0.0315361 | 0.15 | 2 | 41 | Environmental information processing | Signal transduction |
| WikiPathways | G13 signaling pathway | 0.00054157 | 0.00541572 | 0.0270786 | 0.15 | 2 | 38 | Environmental information processing | Signal transduction |
| REACTOME | MET activates PTK2 signaling | 0.0003365 | 0.00560838 | 0.0168252 | 0.15 | 2 | 30 | Environmental information processing | Signal transduction |
| Panther_Pathway | B cell activation | 0.0013038 | 0.00724333 | 0.06519 | 0.15 | 2 | 59 | Organismal systems | Immune system |
| REACTOME | Non-integrin membrane-ECM interactions | 0.0013038 | 0.00724333 | 0.06519 | 0.15 | 2 | 59 | Cellular processes and organization | Cellular community |
| WikiPathways | Nanoparticle-mediated activation of receptor signaling | 0.00029272 | 0.00731803 | 0.0146361 | 0.15 | 2 | 28 | Metabolism | Xenobiotics biodegradation and metabolism |
| REACTOME | Signaling by MET | 0.00232343 | 0.00968096 | 0.116172 | 0.15 | 2 | 79 | Environmental information processing | Signal transduction |
| REACTOME | ECM proteoglycans | 0.00215262 | 0.00978464 | 0.107631 | 0.15 | 2 | 76 | Cellular processes and organization | Cellular community |

**Supplementary Table 4C: PathDIP pathways enriched for the downregulated differentially expressed gene-set list from fibroblasts of knee OA (n=15) vs heathy control donor (n=6) IFPs.** Pathways were filered by adjusted p < 0.05.

| Pathway Source | Pathway Name | p-value | q-value (FDR: BH-method) | q-value (Bonferroni) | Ratio | Query mapped | Pathway size | Type | Category |
|---|---|---|---|---|---|---|---|---|---|
| WikiPathways | T-cell receptor signaling pathway | 0.00010829 | 0.00286976 | 0.00573953 | 0.2 | 3 | 90 | Organismal systems | Immune system |
| REACTOME | Caspase-mediated cleavage of cytoskeletal proteins | 6.93E-05 | 0.00367347 | 0.00367347 | 0.13 | 2 | 12 | Cellular processes and organization | Cell growth and death |
| SIGNOR 3.0 | EGFR Signaling | 0.0002639 | 0.00466227 | 0.0139868 | 0.13 | 2 | 23 | Environmental information processing | Signal transduction |
| WikiPathways | Fatty Acids and Lipoproteins Transport in Hepatocytes | 0.00058277 | 0.00617739 | 0.030887 | 0.27 | 4 | 382 | Organismal systems | Digestive system |
| Panther_Pathway | FAS signaling pathway | 0.00048266 | 0.00639525 | 0.025581 | 0.13 | 2 | 31 | Environmental information processing | Signal transduction |
| REACTOME | Apoptotic cleavage of cellular proteins | 0.00072657 | 0.00641803 | 0.0385082 | 0.13 | 2 | 38 | Cellular processes and organization | Cell growth and death |
| WikiPathways | TGF-beta receptor signaling | 0.00151885 | 0.00894434 | 0.0804991 | 0.13 | 2 | 55 | Environmental information processing | Signal transduction |
| REACTOME | Apoptotic execution phase | 0.00135874 | 0.00900165 | 0.0720132 | 0.13 | 2 | 52 | Cellular processes and organization | Cell growth and death |
| WikiPathways | TROP2 regulatory signaling | 0.00125675 | 0.00951539 | 0.0666078 | 0.13 | 2 | 49 | Environmental information processing | Signal transduction |
| REACTOME | Pre-NOTCH Transcription and Translation | 0.00186485 | 0.00988371 | 0.0988371 | 0.13 | 2 | 61 | Environmental information processing | Signal transduction |
