## Supplementary Table 5 for "Cell and Transcriptomic Diversity of Infrapatellar Fat Pad during Knee Osteoarthritis"

**Supplementary Table 5A: Fibroblast differentially expressed genes from female (n=8) compared to male (n=7) KOA-IFPs.** Upregulated genes are arranged by decreasing Log2 fold change (FC) and downregulated genes are arranged by increasing Log2FC. Genes were filered by adjusted $p < 0.05$, log2FC ≥ 0.5 and min.pct ≥ 0.5.

**Upregulated Genes**

| gene | p_val | avg_log2FC | pct.1 | pct.2 | p_val_adj |
|---|---|---|---|---|---|
| **CLU** | 0 | 2.823536437 | 0.573 | 0.161 | 0 |
| **HTRA1** | 0 | 2.00245381 | 0.516 | 0.199 | 0 |
| **PRG4** | 0 | 1.950146443 | 0.772 | 0.481 | 0 |
| **TMEM196** | 0 | 1.896350265 | 0.506 | 0.122 | 0 |
| **FN1** | 0 | 1.842165777 | 0.861 | 0.607 | 0 |
| **ITGB8** | 0 | 1.802278434 | 0.629 | 0.295 | 0 |
| **ADGRB3** | 0 | 1.6803444 | 0.511 | 0.18 | 0 |
| **PLA2G2A** | 0 | 1.544106989 | 0.564 | 0.212 | 0 |
| **ZNF385B** | 0 | 1.098239 | 0.777 | 0.564 | 0 |
| **ANKH** | 0 | 1.058261399 | 0.624 | 0.41 | 0 |
| **FGF10** | 0 | 1.031539149 | 0.55 | 0.287 | 0 |
| **KAZN** | 0 | 0.918656408 | 0.851 | 0.694 | 0 |
| **ITGBL1** | 8.74E-235 | 0.847622274 | 0.621 | 0.45 | 2.72E-230 |
| **GPC6** | 7.17E-272 | 0.78571472 | 0.605 | 0.393 | 2.23E-267 |
| **UGP2** | 3.17E-211 | 0.76677207 | 0.548 | 0.389 | 9.87E-207 |
| **SOX5** | 0 | 0.755540222 | 0.967 | 0.936 | 0 |
| **C1GALT1** | 0 | 0.753429123 | 0.625 | 0.414 | 0 |
| **COLEC12** | 0 | 0.729842968 | 0.663 | 0.432 | 0 |
| **FOXP2** | 0 | 0.699279019 | 0.723 | 0.442 | 0 |
| **ARHGAP28** | 0 | 0.693719481 | 0.517 | 0.278 | 0 |
| **MGP** | 3.89E-171 | 0.678295317 | 0.504 | 0.344 | 1.21E-166 |
| **CSGALNACT1** | 3.71E-272 | 0.655568088 | 0.537 | 0.318 | 1.16E-267 |
| **COL14A1** | 2.35E-158 | 0.638432676 | 0.514 | 0.357 | 7.31E-154 |
| **GLCCI1** | 0 | 0.62042888 | 0.525 | 0.288 | 0 |
| **CREB5** | 7.39E-307 | 0.608648464 | 0.816 | 0.636 | 2.30E-302 |
| **ITM2B** | 4.35E-181 | 0.582040029 | 0.675 | 0.519 | 1.36E-176 |
| **ARHGAP20** | 7.01E-175 | 0.574279416 | 0.571 | 0.414 | 2.18E-170 |
| **SAT1** | 7.78E-195 | 0.560326272 | 0.526 | 0.353 | 2.42E-190 |
| **ENAH** | 1.01E-178 | 0.544456499 | 0.566 | 0.403 | 3.15E-174 |
| **TRPS1** | 1.70E-297 | 0.542942511 | 0.92 | 0.811 | 5.28E-293 |
| **EXT1** | 3.81E-197 | 0.516625359 | 0.834 | 0.707 | 1.19E-192 |
| **LRP1B** | 2.18E-172 | 0.511820041 | 0.686 | 0.498 | 6.78E-168 |
| **CDK14** | 1.99E-275 | 0.506232772 | 0.846 | 0.705 | 6.21E-271 |

**Downregulated Genes**

| gene | p_val | avg_log2FC | pct.1 | pct.2 | p_val_adj |
|---|---|---|---|---|---|
| **UTY** | 0 | -2.203114529 | 0.001 | 0.691 | 0 |
| **USP9Y** | 0 | -2.047764036 | 0.002 | 0.623 | 0 |
| **LAMA2** | 0 | -1.20708494 | 0.536 | 0.795 | 0 |
| **AFF3** | 0 | -1.185273303 | 0.251 | 0.536 | 0 |
| **NAV3** | 0 | -1.025169009 | 0.249 | 0.52 | 0 |
| **ABCA9** | 0 | -0.938334345 | 0.311 | 0.574 | 0 |
| **ABCA6** | 0 | -0.928734196 | 0.41 | 0.628 | 0 |
| **NEGR1** | 0 | -0.917174734 | 0.613 | 0.823 | 0 |
| **FBLN1** | 0 | -0.906879107 | 0.395 | 0.647 | 0 |
| **FBLN2** | 0 | -0.883932967 | 0.29 | 0.551 | 0 |
| **VIT** | 0 | -0.880744006 | 0.479 | 0.682 | 0 |
| **BMP5** | 0 | -0.863569421 | 0.252 | 0.512 | 0 |
| **EBF2** | 0 | -0.851028575 | 0.41 | 0.609 | 0 |
| **GSN** | 2.54E-290 | -0.834030968 | 0.747 | 0.809 | 7.91E-286 |
| **NTM** | 0 | -0.831840834 | 0.357 | 0.622 | 0 |
| **ITGA11** | 0 | -0.830225076 | 0.352 | 0.609 | 0 |
| **MFAP5** | 0 | -0.829563391 | 0.289 | 0.538 | 0 |
| **ABCA8** | 1.52E-216 | -0.756407 | 0.572 | 0.683 | 4.73E-212 |
| **DOCK4** | 0 | -0.73812828 | 0.418 | 0.634 | 0 |
| **SESTD1** | 0 | -0.732353324 | 0.583 | 0.758 | 0 |
| **GHR** | 6.72E-217 | -0.71946496 | 0.507 | 0.641 | 2.09E-212 |
| **NOVA1** | 9.43E-288 | -0.699689959 | 0.877 | 0.886 | 2.94E-283 |
| **TGFBR3** | 4.21E-74 | -0.699391013 | 0.665 | 0.698 | 1.31E-69 |
| **ZEB1** | 0 | -0.698610629 | 0.858 | 0.922 | 0 |
| **TNXB** | 0 | -0.692919687 | 0.808 | 0.893 | 0 |
| **ADGRD1** | 1.14E-270 | -0.68721924 | 0.307 | 0.5 | 3.56E-266 |
| **LTBP4** | 3.65E-247 | -0.686662961 | 0.426 | 0.582 | 1.14E-242 |
| **BCL6** | 3.43E-159 | -0.682793555 | 0.391 | 0.515 | 1.07E-154 |
| **DCN** | 0 | -0.678448065 | 0.783 | 0.872 | 0 |
| **XG** | 3.72E-301 | -0.677911531 | 0.318 | 0.518 | 1.16E-296 |
| **SPTBN1** | 0 | -0.654464126 | 0.674 | 0.844 | 0 |
| **UTRN** | 0 | -0.652590689 | 0.71 | 0.869 | 0 |
| **OPHN1** | 0 | -0.651333722 | 0.529 | 0.7 | 0 |
| **COL12A1** | 2.62E-106 | -0.648112105 | 0.413 | 0.511 | 8.17E-102 |
| **FBN1** | 9.20E-285 | -0.64734426 | 0.768 | 0.852 | 2.87E-280 |
| **TRIO** | 2.01E-260 | -0.635234565 | 0.655 | 0.786 | 6.25E-256 |
| **NOX4** | 6.40E-263 | -0.630121901 | 0.479 | 0.668 | 1.99E-258 |
| **AHNAK** | 4.48E-235 | -0.626301382 | 0.642 | 0.751 | 1.40E-230 |
| **CELF2** | 0 | -0.615660928 | 0.736 | 0.897 | 0 |
| **STARD9** | 1.64E-288 | -0.609602312 | 0.535 | 0.716 | 5.11E-284 |
| **RBPJ** | 1.51E-271 | -0.604718456 | 0.692 | 0.794 | 4.71E-267 |
| **DST** | 1.93E-241 | -0.601335867 | 0.913 | 0.94 | 6.02E-237 |
| **PARD3B** | 0 | -0.59831534 | 0.826 | 0.889 | 0 |
| **SVEP1** | 1.66E-156 | -0.592532595 | 0.415 | 0.542 | 5.18E-152 |
| **CAMK1D** | 3.67E-70 | -0.581669532 | 0.724 | 0.728 | 1.14E-65 |
| **DDR2** | 2.25E-233 | -0.581093752 | 0.636 | 0.749 | 7.00E-229 |
| **ANKRD12** | 1.37E-275 | -0.579307236 | 0.725 | 0.817 | 4.28E-271 |
| **FMNL2** | 4.33E-203 | -0.566899193 | 0.338 | 0.512 | 1.35E-198 |
| **ABLIM1** | 5.72E-226 | -0.565362302 | 0.539 | 0.689 | 1.78E-221 |
| **CAB39L** | 6.39E-191 | -0.561735646 | 0.441 | 0.584 | 1.99E-186 |
| **PROCR** | 1.27E-89 | -0.555348452 | 0.431 | 0.506 | 3.94E-85 |
| **CCN5** | 1.34E-156 | -0.554220077 | 0.494 | 0.608 | 4.17E-152 |
| **DENND2A** | 5.38E-138 | -0.547063038 | 0.372 | 0.502 | 1.68E-133 |
| **DMD** | 9.54E-166 | -0.547031451 | 0.428 | 0.58 | 2.97E-161 |
| **PCNX2** | 9.95E-200 | -0.544252303 | 0.436 | 0.594 | 3.10E-195 |
| **ZBTB16** | 4.25E-105 | -0.542522861 | 0.459 | 0.545 | 1.32E-100 |
| **ANKRD36C** | 3.35E-181 | -0.540535882 | 0.555 | 0.674 | 1.04E-176 |
| **PID1** | 1.71E-90 | -0.529593359 | 0.571 | 0.63 | 5.32E-86 |
| **AGAP1** | 6.19E-115 | -0.522677307 | 0.697 | 0.745 | 1.93E-110 |
| **MGST1** | 3.70E-125 | -0.520048015 | 0.395 | 0.516 | 1.15E-120 |
| **RBM25** | 3.26E-137 | -0.519854489 | 0.666 | 0.735 | 1.02E-132 |
| **PLPP3** | 2.25E-152 | -0.519197634 | 0.568 | 0.671 | 7.01E-148 |
| **FBXL7** | 4.85E-188 | -0.518931222 | 0.75 | 0.825 | 1.51E-183 |
| **ARHGAP10** | 2.79E-189 | -0.517217803 | 0.505 | 0.638 | 8.70E-185 |
| **ZHX3** | 1.81E-138 | -0.514056093 | 0.398 | 0.519 | 5.64E-134 |
| **DCLK1** | 4.35E-243 | -0.512756272 | 0.616 | 0.794 | 1.35E-238 |
| **COL6A1** | 9.17E-190 | -0.511554841 | 0.587 | 0.699 | 2.86E-185 |
| **CFD** | 2.63E-13 | -0.509084517 | 0.636 | 0.638 | 8.20E-09 |
| **SRRM2** | 5.38E-155 | -0.505771615 | 0.749 | 0.815 | 1.68E-150 |
| **HSPG2** | 2.53E-196 | -0.502316029 | 0.569 | 0.687 | 7.88E-192 |

**Supplementary Table 5B: Gene ontology biological processes linked to upregulated genes in fibroblasts of female (n=8) compared to male (n=7) KOA-IFPs.** Biological functions were filtered by adjusted p < 0.05.

| ID | Description | GeneRatio | BgRatio | pvalue | p.adjust | qvalue | geneID | Count |
|---|---|---|---|---|---|---|---|---|
| GO:0051216 | cartilage development | 3/31 | 201/18903 | 1.75E-05 | 0.015478029 | 0.011231223 | ITGB8/SOX5/MGP/TRPS1/EXT1 | 5 |
| GO:0061448 | connective tissue development | 3/31 | 274/18903 | 7.70E-05 | 0.033948971 | 0.024634174 | ITGB8/SOX5/MGP/TRPS1/EXT1 | 5 |
| GO:0030210 | heparin biosynthetic process | 2/31 | 11/18903 | 0.000141843 | 0.037572093 | 0.027263197 | CSGALNACT1/EXT1 | 2 |
| GO:0033692 | cellular polysaccharide biosynthetic process | 3/31 | 66/18903 | 0.000170395 | 0.037572093 | 0.027263197 | UGP2/CSGALNACT1/EXT1 | 3 |
| GO:0000271 | polysaccharide biosynthetic process | 3/31 | 73/18903 | 0.000229806 | 0.038795151 | 0.028150677 | UGP2/CSGALNACT1/EXT1 | 3 |
| GO:0034637 | cellular carbohydrate biosynthetic process | 3/31 | 78/18903 | 0.000279536 | 0.038795151 | 0.028150677 | UGP2/CSGALNACT1/EXT1 | 3 |
| GO:0030202 | heparin metabolic process | 2/31 | 16/18903 | 0.000307898 | 0.038795151 | 0.028150677 | CSGALNACT1/EXT1 | 2 |
| GO:0033627 | cell adhesion mediated by integrin | 3/31 | 88/18903 | 0.000398771 | 0.042866325 | 0.031104816 | ITGB8/ITGBL1/EXT1 | 3 |
| GO:0001502 | cartilage condensation | 2/31 | 19/18903 | 0.000437411 | 0.042866325 | 0.031104816 | SOX5/MGP | 2 |
| GO:0044264 | cellular polysaccharide metabolic process | 3/31 | 99/18903 | 0.000563068 | 0.043297433 | 0.031417638 | UGP2/CSGALNACT1/EXT1 | 3 |
| GO:0002274 | myeloid leukocyte activation | 4/31 | 237/18903 | 0.000580946 | 0.043297433 | 0.031417638 | CLU/FN1/ITGB8/PLA2G2A | 4 |
| GO:0098743 | cell aggregation | 2/31 | 22/18903 | 0.000589081 | 0.043297433 | 0.031417638 | SOX5/MGP | 2 |
| GO:0005976 | polysaccharide metabolic process | 3/31 | 110/18903 | 0.000765382 | 0.049921713 | 0.036224372 | UGP2/CSGALNACT1/EXT1 | 3 |
| GO:0007229 | integrin-mediated signaling pathway | 3/31 | 113/18903 | 0.000827583 | 0.049921713 | 0.036224372 | FN1/ITGB8/ITGBL1 | 3 |
| GO:0002062 | chondrocyte differentiation | 3/31 | 114/18903 | 0.000849009 | 0.049921713 | 0.036224372 | SOX5/TRPS1/EXT1 | 3 |
| GO:0060441 | epithelial tube branching involved in lung morphogenesis | 2/31 | 29/18903 | 0.001027979 | 0.054543023 | 0.039577703 | FGF10/EXT1 | 2 |
| GO:0030204 | chondroitin sulfate metabolic process | 2/31 | 30/18903 | 0.001100282 | 0.054543023 | 0.039577703 | CSGALNACT1/EXT1 | 2 |
| GO:0015012 | heparan sulfate proteoglycan biosynthetic process | 2/31 | 31/18903 | 0.001174963 | 0.054543023 | 0.039577703 | CSGALNACT1/EXT1 | 2 |
| GO:0031069 | hair follicle morphogenesis | 2/31 | 31/18903 | 0.001174963 | 0.054543023 | 0.039577703 | FGF10/EXT1 | 2 |
| GO:0048565 | digestive tract development | 3/31 | 132/18903 | 0.001296712 | 0.056738974 | 0.041171138 | FGF10/C1GALT1/EXT1 | 3 |
| GO:0042476 | odontogenesis | 3/31 | 135/18903 | 0.001383258 | 0.056738974 | 0.041171138 | HTRA1/ANKH/FGF10 | 3 |
| GO:0048730 | epidermis morphogenesis | 2/31 | 35/18903 | 0.001497322 | 0.056738974 | 0.041171138 | FGF10/EXT1 | 2 |
| GO:0050654 | chondroitin sulfate proteoglycan metabolic process | 2/31 | 35/18903 | 0.001497322 | 0.056738974 | 0.041171138 | CSGALNACT1/EXT1 | 2 |
| GO:0055123 | digestive system development | 3/31 | 143/18903 | 0.001631593 | 0.056738974 | 0.041171138 | FGF10/C1GALT1/EXT1 | 3 |
| GO:0009225 | nucleotide-sugar metabolic process | 2/31 | 37/18903 | 0.001672577 | 0.056738974 | 0.041171138 | UGP2/CSGALNACT1 | 2 |
| GO:0010092 | specification of animal organ identity | 2/31 | 37/18903 | 0.001672577 | 0.056738974 | 0.041171138 | FGF10/EXT1 | 2 |
| GO:0009101 | glycoprotein biosynthetic process | 4/31 | 321/18903 | 0.001788208 | 0.058414782 | 0.042387144 | C1GALT1/CSGALNACT1/ITM2B/EXT1 | 4 |
| GO:0071634 | regulation of transforming growth factor beta production | 2/31 | 39/18903 | 0.001857138 | 0.05849984 | 0.042448864 | FN1/ITGB8 | 2 |
| GO:0030201 | heparan sulfate proteoglycan metabolic process | 2/31 | 40/18903 | 0.001952889 | 0.059394758 | 0.043098237 | CSGALNACT1/EXT1 | 2 |
| GO:0071604 | transforming growth factor beta production | 2/31 | 42/18903 | 0.002151296 | 0.06324811 | 0.04589432 | FN1/ITGB8 | 2 |

**Supplementary Table 5C: PathDIP pathways linked to the upregulated differentially expressed gene-set list from fibroblasts of female (n=7) compared to male (n=8) KOA-IFPs.** Pathways were filtered by adjusted p <0.05.

| Pathway Source | Pathway Name | p-value | q-value (FDR: BH-method) | q-value (Bonferroni) | Ratio | Query mapped | Pathway size | Type | Category |
|---|---|---|---|---|---|---|---|---|---|
| KEGG | Regulation of actin cytoskeleton | 0.000636514 | 0.00700165 | 0.0350083 | 0.15 | 4 | 218 | Organismal systems | Cell motility |
| WikiPathways | Proteoglycan biosynthesis | 0.0004914 | 0.009009 | 0.027027 | 0.08 | 2 | 18 | Metabolism | Carbohydrate metabolism |
| WikiPathways | miR-509-3p alteration of YAP1/ECM axis | 0.0004914 | 0.009009 | 0.027027 | 0.08 | 2 | 18 | Cellular processes and organization | Cellular community |
| WikiPathways | Focal adhesion: PI3K-Akt-mTOR-signaling pathway | 0.00215258 | 0.00910707 | 0.118392 | 0.15 | 4 | 303 | Cellular processes and organization | Cellular community |
| REACTOME | Degradation of the extracellular matrix | 0.00210808 | 0.00966203 | 0.115944 | 0.12 | 3 | 140 | Cellular processes and organization | Cellular community |
| WikiPathways | Regulation of actin cytoskeleton | 0.00256514 | 0.0100773 | 0.141083 | 0.12 | 3 | 150 | Organismal systems | Cell motility |
| REACTOME | Elastic fibre formation | 0.00295015 | 0.0101411 | 0.162258 | 0.08 | 2 | 44 | Cellular processes and organization | Cellular community |
| MetabolicAtlas | chondroitin heparan sulfate biosynthesis | 0.00281923 | 0.0103372 | 0.155058 | 0.08 | 2 | 43 | Metabolism | Carbohydrate metabolism |
| WikiPathways | PI3K-Akt signaling pathway | 0.00319697 | 0.0103431 | 0.175833 | 0.15 | 4 | 338 | Environmental information processing | Signal transduction |
| REACTOME | Molecules associated with elastic fibres | 0.00209341 | 0.0104671 | 0.115138 | 0.08 | 2 | 37 | Cellular processes and organization | Cellular community |
| ACSN2 | ECM | 0.00159526 | 0.0109674 | 0.0877393 | 0.12 | 3 | 127 | Cellular processes and organization | Cellular community |
| REACTOME | Chondroitin sulfate/dermatan sulfate metabolism | 0.00379434 | 0.0109836 | 0.208689 | 0.08 | 2 | 50 | Metabolism | Carbohydrate metabolism |
| KEGG | PI3K-Akt signaling pathway | 0.00366036 | 0.0111844 | 0.20132 | 0.15 | 4 | 351 | Environmental information processing | Signal transduction |
| REACTOME | Extracellular matrix organization | 0.00207595 | 0.0114177 | 0.114177 | 0.15 | 4 | 300 | Cellular processes and organization | Cellular community |
| REACTOME | Glycosaminoglycan metabolism | 0.00148946 | 0.0117029 | 0.0819203 | 0.12 | 3 | 124 | Metabolism | Carbohydrate metabolism |
| WikiPathways | NOTCH1 regulation of endothelial cell calcification | 0.000437285 | 0.0120253 | 0.0240507 | 0.08 | 2 | 17 | Environmental information processing | Signal transduction |
| REACTOME | Metabolism of carbohydrates | 0.00197679 | 0.0120804 | 0.108723 | 0.15 | 4 | 296 | Metabolism | Carbohydrate metabolism |
| Panther_Pathway | Integrin signalling pathway | 0.000221216 | 0.0121669 | 0.0121669 | 0.15 | 4 | 165 | Environmental information processing | Signal transduction |
| REACTOME | HS-GAG biosynthesis | 0.00147161 | 0.0134898 | 0.0809386 | 0.08 | 2 | 31 | Metabolism | Carbohydrate metabolism |
| ACSN2 | CELL MATRIX ADHESIONS | 0.00614627 | 0.0160974 | 0.338045 | 0.08 | 2 | 64 | Cellular processes and organization | Cellular community |
| WikiPathways | Endochondral ossification | 0.00596115 | 0.0163932 | 0.327863 | 0.08 | 2 | 63 | Organismal systems | Development and regeneration |
| REACTOME | Integrin cell surface interactions | 0.0106294 | 0.0265735 | 0.584617 | 0.08 | 2 | 85 | Cellular processes and organization | Cellular community |
| KEGG | ECM-receptor interaction | 0.0113592 | 0.0271633 | 0.624756 | 0.08 | 2 | 88 | Cellular processes and organization | Cellular community |
| REACTOME | Antimicrobial peptides | 0.0123657 | 0.0283381 | 0.680114 | 0.08 | 2 | 92 | Organismal systems | Immune system |
| ACSN2 | IMMUNOSTIMULATORY CORE PATHWAYS | 0.0150457 | 0.0331005 | 0.827514 | 0.08 | 2 | 102 | Organismal systems | Immune system |
| WikiPathways | Complement system in neuronal development and plasticity | 0.0158944 | 0.0336228 | 0.874192 | 0.08 | 2 | 105 | Organismal systems | Nervous system |
| REACTOME | Platelet degranulation | 0.0230579 | 0.0469698 | 1 | 0.08 | 2 | 128 | Organismal systems | Immune system |
| REACTOME | Response to elevated platelet cytosolic Ca2+ | 0.0247622 | 0.04864 | 1 | 0.08 | 2 | 133 | Organismal systems | Immune system |

**Supplementary Table 5D: Gene ontology biological functions enriched for the downregulated differentially expressed genes from fibroblasts of female (n=8) comared to male (n=7) KOA-IFPs.** Biological functions were filtered by adjusted p < 0.05.

| ID | Description | GeneRatio | BgRatio | pvalue | p.adjust | qvalue | geneID | Count |
|---|---|---|---|---|---|---|---|---|
| GO:0007178 | transmembrane receptor protein serine/threonine kinase signaling pathway | 8/68 | 390/18903 | 7.67E-05 | 0.046282891 | 0.040353296 | USP9Y/BMP5/TGFBR3/ZEB1/LTBP4/SPTBN1/FBN1/RBPJ | 8 |
| GO:0035994 | response to muscle stretch | 3/68 | 26/18903 | 0.000109098 | 0.046282891 | 0.040353296 | GSN/DDR2/DMD | 3 |
| GO:0030198 | extracellular matrix organization | 7/68 | 318/18903 | 0.000143685 | 0.046282891 | 0.040353296 | LAMA2/FBLN1/FBLN2/VIT/TNXB/COL12A1/DDR2 | 7 |
| GO:0043062 | extracellular structure organization | 7/68 | 319/18903 | 0.000146494 | 0.046282891 | 0.040353296 | LAMA2/FBLN1/FBLN2/VIT/TNXB/COL12A1/DDR2 | 7 |
| GO:0045229 | external encapsulating structure organization | 7/68 | 321/18903 | 0.000152246 | 0.046282891 | 0.040353296 | LAMA2/FBLN1/FBLN2/VIT/TNXB/COL12A1/DDR2 | 7 |
| GO:0061384 | heart trabecula morphogenesis | 3/68 | 32/18903 | 0.000204936 | 0.051917094 | 0.045265665 | BMP5/TGFBR3/RBPJ | 3 |
| GO:0048048 | embryonic eye morphogenesis | 3/68 | 35/18903 | 0.000268346 | 0.056075825 | 0.048891595 | MFAP5/ZEB1/FBN1 | 3 |
| GO:0030509 | BMP signaling pathway | 5/68 | 167/18903 | 0.000337149 | 0.056075825 | 0.048891595 | USP9Y/BMP5/TGFBR3/FBN1/RBPJ | 5 |
| GO:0031589 | cell-substrate adhesion | 7/68 | 369/18903 | 0.000355662 | 0.056075825 | 0.048891595 | FBLN1/FBLN2/VIT/ITGA11/TNXB/BCL6/UTRN | 7 |
| GO:0071560 | cellular response to transforming growth factor beta stimulus | 6/68 | 271/18903 | 0.000427157 | 0.056075825 | 0.048891595 | USP9Y/TGFBR3/ZEB1/LTBP4/FBN1/DDR2 | 6 |
| GO:0071772 | response to BMP | 5/68 | 178/18903 | 0.00045162 | 0.056075825 | 0.048891595 | USP9Y/BMP5/TGFBR3/FBN1/RBPJ | 5 |
| GO:0071773 | cellular response to BMP stimulus | 5/68 | 178/18903 | 0.00045162 | 0.056075825 | 0.048891595 | USP9Y/BMP5/TGFBR3/FBN1/RBPJ | 5 |
| GO:0071559 | response to transforming growth factor beta | 6/68 | 277/18903 | 0.000479596 | 0.056075825 | 0.048891595 | USP9Y/TGFBR3/ZEB1/LTBP4/FBN1/DDR2 | 6 |

**Supplementary Table 5E: PathDIP Pathways downregulated differentially expressed gene-set list from fibroblasts of female (n=8) comared to male (n=7) KOA-IFPs.** Pathways were filtered by adjusted p < 0.05.

| Pathway Source | Pathway Name | p-value | q-value (FDR: BH-method) | q-value (Bonferroni) | Ratio | Query mapped | Pathway size | Type | Category |
|---|---|---|---|---|---|---|---|---|---|
| REACTOME | Extracellular matrix organization | 2.39E-14 | 2.75E-12 | 2.75E-12 | 0.32 | 15 | 300 | Cellular processes and organization | Cellular community |
| REACTOME | Molecules associated with elastic fibres | 1.33E-07 | 7.66E-06 | 1.53E-05 | 0.11 | 5 | 37 | Cellular processes and organization | Cellular community |
| REACTOME | Elastic fibre formation | 3.26E-07 | 1.25E-05 | 3.75E-05 | 0.11 | 5 | 44 | Cellular processes and organization | Cellular community |
| REACTOME | ECM proteoglycans | 5.12E-06 | 0.000147257 | 0.000589029 | 0.11 | 5 | 76 | Cellular processes and organization | Cellular community |
| KEGG | ECM-receptor interaction | 1.05E-05 | 0.000242475 | 0.00121238 | 0.11 | 5 | 88 | Cellular processes and organization | Cellular community |
| REACTOME | Non-integrin membrane-ECM interactions | 4.31E-05 | 0.000826902 | 0.00496141 | 0.09 | 4 | 59 | Cellular processes and organization | Cellular community |
| REACTOME | Degradation of the extracellular matrix | 9.86E-05 | 0.00162011 | 0.0113407 | 0.11 | 5 | 140 | Cellular processes and organization | Cellular community |
| REACTOME | Integrin cell surface interactions | 0.000180155 | 0.00230198 | 0.0207178 | 0.09 | 4 | 85 | Cellular processes and organization | Cellular community |
| Panther_Pathway | Integrin signalling pathway | 0.000213095 | 0.00245059 | 0.0245059 | 0.11 | 5 | 165 | Environmental information processing | Signal transduction |
| BioCarta | BIOCARTA AGR PATHWAY | 0.00017663 | 0.00253906 | 0.0203125 | 0.06 | 3 | 33 | Organismal systems | Nervous system |
| WikiPathways | miRNA targets in ECM and membrane receptors | 0.0003903 | 0.00408041 | 0.0448845 | 0.06 | 3 | 43 | Cellular processes and organization | Cellular community |
| KEGG | ABC transporters | 0.00044668 | 0.00428068 | 0.0513682 | 0.06 | 3 | 45 | Environmental information processing | Membrane transport |
| REACTOME | RHO GTPase cycle | 0.000670936 | 0.0059352 | 0.0771576 | 0.15 | 7 | 450 | Environmental information processing | Signal transduction |
| ACSN2 | ECM | 0.000830212 | 0.0068196 | 0.0954744 | 0.09 | 4 | 127 | Cellular processes and organization | Cellular community |
| REACTOME | DCC mediated attractive signaling | 0.000964978 | 0.00739816 | 0.110972 | 0.04 | 2 | 14 | Organismal systems | Development and regeneration |
| REACTOME | Assembly of collagen fibrils and other multimeric structures | 0.00109145 | 0.0078448 | 0.125517 | 0.06 | 3 | 61 | Cellular processes and organization | Cellular community |
| REACTOME | ABC transporters in lipid homeostasis | 0.00160871 | 0.0108825 | 0.185002 | 0.04 | 2 | 18 | Environmental information processing | Membrane transport |
| REACTOME | CDC42 GTPase cycle | 0.00173601 | 0.0110912 | 0.199641 | 0.09 | 4 | 155 | Environmental information processing | Signal transduction |
| REACTOME | RHOG GTPase cycle | 0.00190719 | 0.0115435 | 0.219327 | 0.06 | 3 | 74 | Environmental information processing | Signal transduction |
| BioCarta | BIOCARTA RHO PATHWAY | 0.00219402 | 0.0126156 | 0.252312 | 0.04 | 2 | 21 | Organismal systems | Cell motility |
| REACTOME | A tetrasaccharide linker sequence is required for GAG synthesis | 0.00335971 | 0.0154547 | 0.386367 | 0.04 | 2 | 26 | Metabolism | Carbohydrate metabolism |
| REACTOME | Collagen formation | 0.00333024 | 0.0159574 | 0.382978 | 0.06 | 3 | 90 | Cellular processes and organization | Cellular community |
| WikiPathways | Focal adhesion: PI3K-Akt-mTOR-signaling pathway | 0.00325132 | 0.0162566 | 0.373902 | 0.11 | 5 | 303 | Cellular processes and organization | Cellular community |
| REACTOME | EGR2 and SOX10-mediated initiation of Schwann cell myelination | 0.00389101 | 0.0165728 | 0.447466 | 0.04 | 2 | 28 | Organismal systems | Development and regeneration |
| KEGG | TGF-beta signaling pathway | 0.00376462 | 0.0166512 | 0.432931 | 0.06 | 3 | 94 | Environmental information processing | Signal transduction |
| REACTOME | RAC1 GTPase cycle | 0.00323338 | 0.0169018 | 0.371839 | 0.09 | 4 | 184 | Environmental information processing | Signal transduction |
| REACTOME | RAC2 GTPase cycle | 0.00312526 | 0.0171145 | 0.359405 | 0.06 | 3 | 88 | Environmental information processing | Signal transduction |
| REACTOME | Laminin interactions | 0.00445884 | 0.0176816 | 0.512767 | 0.04 | 2 | 30 | Cellular processes and organization | Cellular community |
| KEGG | Focal adhesion | 0.00443105 | 0.018199 | 0.509571 | 0.09 | 4 | 201 | Cellular processes and organization | Cellular community |
| REACTOME | ABC-family proteins mediated transport | 0.00486423 | 0.0186462 | 0.559386 | 0.06 | 3 | 103 | Environmental information processing | Membrane transport |
| WikiPathways | PI3K-Akt signaling pathway | 0.00516848 | 0.0191734 | 0.594375 | 0.11 | 5 | 338 | Environmental information processing | Signal transduction |
| WikiPathways | Oxidative stress response | 0.0053779 | 0.0193268 | 0.618458 | 0.04 | 2 | 33 | Cellular processes and organization | Cellular response to stimuli |
| KEGG | PI3K-Akt signaling pathway | 0.00605127 | 0.0210878 | 0.695896 | 0.11 | 5 | 351 | Environmental information processing | Signal transduction |
| REACTOME | Signaling by Rho GTPases | 0.00651722 | 0.0220435 | 0.74948 | 0.15 | 7 | 673 | Environmental information processing | Signal transduction |
| BioCarta | BIOCARTA ALK PATHWAY | 0.00672614 | 0.0221002 | 0.773506 | 0.04 | 2 | 37 | Organismal systems | Cardiovascular system |
| REACTOME | RHOV GTPase cycle | 0.00672614 | 0.0221002 | 0.773506 | 0.04 | 2 | 37 | Environmental information processing | Signal transduction |
| REACTOME | Signaling by Rho GTPases, Miro GTPases and RHOBTB3 | 0.00738529 | 0.0229543 | 0.849308 | 0.15 | 7 | 689 | Environmental information processing | Signal transduction |

| | | | | | | | | | |
|---|---|---|---|---|---|---|---|---|---|
| REACTOME | RHOU GTPase cycle | 0.00782778 | 0.0236893 | 0.900195 | 0.04 | 2 | 40 | Environmental information processing | Signal transduction |
| MetabolicAtlas | eicosanoid metabolism | 0.00900487 | 0.0265528 | 1 | 0.04 | 2 | 43 | Metabolism | Xenobiotics biodegradation and metabolism |
| WikiPathways | TGF-beta signaling pathway | 0.00963566 | 0.0270269 | 1 | 0.06 | 3 | 132 | Environmental information processing | Signal transduction |
| REACTOME | Collagen chain trimerization | 0.00941382 | 0.0270647 | 1 | 0.04 | 2 | 44 | Cellular processes and organization | Cellular community |
| REACTOME | TGF-beta receptor signaling activates SMADs | 0.0102562 | 0.0280825 | 1 | 0.04 | 2 | 46 | Environmental information processing | Signal transduction |
| REACTOME | Nervous system development | 0.0120997 | 0.0296056 | 1 | 0.13 | 6 | 580 | Organismal systems | Development and regeneration |
| WikiPathways | Ectoderm differentiation | 0.0117407 | 0.0313995 | 1 | 0.06 | 3 | 142 | Organismal systems | Development and regeneration |
| MetabolicAtlas | prostaglandin biosynthesis | 0.0120368 | 0.0314598 | 1 | 0.04 | 2 | 50 | Metabolism | Lipid metabolism |
| REACTOME | Chondroitin sulfate/dermatan sulfate metabolism | 0.0120368 | 0.0314598 | 1 | 0.04 | 2 | 50 | Metabolism | Carbohydrate metabolism |
| REACTOME | Netrin-1 signaling | 0.0120368 | 0.0314598 | 1 | 0.04 | 2 | 50 | Organismal systems | Development and regeneration |
| REACTOME | RHOA GTPase cycle | 0.0133601 | 0.0320086 | 1 | 0.06 | 3 | 149 | Environmental information processing | Signal transduction |
| REACTOME | RHOJ GTPase cycle | 0.0144389 | 0.0332095 | 1 | 0.04 | 2 | 55 | Environmental information processing | Signal transduction |
| REACTOME | Sensory processing of sound by outer hair cells of the cochlea | 0.0144389 | 0.0332095 | 1 | 0.04 | 2 | 55 | Organismal systems | Sensory system |
| ACSN2 | EMT REGULATORS | 0.0142266 | 0.033389 | 1 | 0.13 | 6 | 599 | Environmental information processing | Signal transduction |
| ACSN2 | CELL MATRIX ADHESIONS | 0.0192337 | 0.0409607 | 1 | 0.04 | 2 | 64 | Cellular processes and organization | Cellular community |
| REACTOME | Collagen degradation | 0.0192337 | 0.0409607 | 1 | 0.04 | 2 | 64 | Cellular processes and organization | Cellular community |
| REACTOME | NCAM signaling for neurite out-growth | 0.0186721 | 0.0412941 | 1 | 0.04 | 2 | 63 | Organismal systems | Development and regeneration |
| WikiPathways | Endochondral ossification | 0.0186721 | 0.0412941 | 1 | 0.04 | 2 | 63 | Organismal systems | Development and regeneration |
| REACTOME | FOXO-mediated transcription | 0.0203786 | 0.0418489 | 1 | 0.04 | 2 | 66 | Genetic information processing | Transcription |
| REACTOME | Collagen biosynthesis and modifying enzymes | 0.0209618 | 0.0422914 | 1 | 0.04 | 2 | 67 | Cellular processes and organization | Cellular community |
| WikiPathways | Glucocorticoid receptor pathway | 0.0227524 | 0.0451125 | 1 | 0.04 | 2 | 70 | Organismal systems | Endocrine system |
| REACTOME | RHOC GTPase cycle | 0.0252337 | 0.0491843 | 1 | 0.04 | 2 | 74 | Environmental information processing | Signal transduction |
