## Supplementary Table 6 for "Cell and Transcriptomic Diversity of Infrapatellar Fat Pad during Knee Osteoarthritis"

**Supplementary Table 6A: Fibroblast differentially expressed genes from obese BMI (n=8) compared to normal BMI (n=7) KOA-IFPs.** Upregulated genes are arranged by decreasing Log2 fold change (FC) and downregulated genes are arranged by increasing Log2 FC. Genes were filered by adjusted p < 0.05, log2FC ≥ 0.5 and min.pct ≥ 0.5.

**Upregulated Genes**

| gene | p_val | avg_log2FC | pct.1 | pct.2 | p_val_adj |
|---|---|---|---|---|---|
| **HMCN1** | 7.37E-207 | 0.947700428 | 0.571 | 0.416 | 2.30E-202 |
| **LHFPL6** | 5.29E-260 | 0.67108206 | 0.755 | 0.612 | 1.65E-255 |
| **ABCA6** | 1.30E-150 | 0.66055404 | 0.584 | 0.451 | 4.05E-146 |
| **PRRX1** | 0 | 0.618804282 | 0.829 | 0.681 | 0 |
| **ABCA8** | 2.47E-121 | 0.584290938 | 0.672 | 0.585 | 7.69E-117 |
| **ITGA11** | 2.82E-191 | 0.545358663 | 0.572 | 0.391 | 8.78E-187 |
| **DST** | 9.96E-262 | 0.540137038 | 0.943 | 0.912 | 3.10E-257 |
| **STARD9** | 1.12E-213 | 0.529031426 | 0.713 | 0.547 | 3.47E-209 |
| **SRRM2** | 9.36E-137 | 0.520312524 | 0.826 | 0.745 | 2.92E-132 |
| **FBLN1** | 4.11E-150 | 0.508126167 | 0.593 | 0.444 | 1.28E-145 |

**Downregulated Genes**

| gene | p_val | avg_log2FC | pct.1 | pct.2 | p_val_adj |
|---|---|---|---|---|---|
| **PRG4** | 0 | -1.8148305 | 0.493 | 0.749 | 0 |
| **CLU** | 0 | -1.819643444 | 0.22 | 0.513 | 0 |
| **FN1** | 7.19E-171 | -0.855025986 | 0.687 | 0.795 | 2.24E-166 |
| **ITGB8** | 1.86E-130 | -0.831616351 | 0.414 | 0.533 | 5.78E-126 |
| **FKBP5** | 9.38E-268 | -0.679533488 | 0.309 | 0.511 | 2.92E-263 |
| **UGP2** | 1.36E-129 | -0.675889959 | 0.415 | 0.522 | 4.24E-125 |
| **TIMP3** | 7.45E-181 | -0.646089085 | 0.702 | 0.815 | 2.32E-176 |
| **GPC6** | 3.57E-184 | -0.597113093 | 0.406 | 0.585 | 1.11E-179 |
| **ZBTB16** | 1.72E-235 | -0.559000991 | 0.378 | 0.574 | 5.36E-231 |
| **ITM2B** | 6.31E-155 | -0.539169995 | 0.533 | 0.658 | 1.97E-150 |
| **CREB5** | 3.23E-162 | -0.518138991 | 0.663 | 0.788 | 1.01E-157 |

**Supplementary Table 6B: Upregulated genes in fibroblasts from obese BMI (n=6) vs normal BMI (n=6) KOA-IFPs identified by spatial transcriptomics.** The list of upregulated differentially expressed genes in fibroblasts from obese BMI vs normal BMI KOA-IFPs identified using snRNA-seq DE analysis (Figure 6D) was searched against the entire spatially-resolved DEG list, identifying 9 genes. All 9 genes were upregulated in fibroblasts of obese vs. normal BMI KOA-IFPs, with 3 significantly upregulated, by spatial transcriptomics (adjusted p < 0.05, bolded).

| Gene | p-value | Average Log2FC | pct.1 | pct.2 | Adjusted p-value |
|---|---|---|---|---|---|
| **ITGA11** | 6.95232616793603e-09 | 0.701168467 | 0.334 | 0.268 | **0.000125163** |
| **STARD9** | 4.45859207347807e-08 | 0.569447103 | 0.136 | 0.083 | **0.00080268** |
| **ABCA6** | 1.4638478239807e-07 | 1.067385404 | 0.113 | 0.068 | **0.002635365** |
| **SRRM2** | 5.62272627957108e-05 | 0.203989396 | 0.377 | 0.331 | 1 |
| **HMCN1** | 6.57892093247976e-05 | 0.71502616 | 0.085 | 0.053 | 1 |
| **DST** | 0.011122594 | 0.113093422 | 0.225 | 0.196 | 1 |
| **LHFPL6** | 0.025832481 | 0.311569943 | 0.357 | 0.343 | 1 |
| **PRRX1** | 0.268042072 | 0.127942034 | 0.49 | 0.485 | 1 |
| **ABCA8** | 0.743731056 | 0.108130772 | 0.134 | 0.132 | 1 |

**Supplementary Table 6C: Gene ontology biological processes enriched in the upregulated differentially expressed genes from fibroblasts of obese BMI (n=8) vs normal BMI (n=7) KOA-IFPs.** Biological processes were filtered by adjusted p < 0.05.

| ID | Description | GeneRatio | BgRatio | pvalue | p.adjust | qvalue | geneID | Count |
|---|---|---|---|---|---|---|---|---|
| GO:0072091 | regulation of stem cell proliferation | 2/9 | 91/18614 | 8.32E-04 | 0.06931282 | 0.040077094 | PRRX1/FBLN1 | 2 |
| GO:0007229 | integrin-mediated signaling pathway | 2/9 | 112/18614 | 0.001256614 | 0.06931282 | 0.040077094 | ITGA11/DST | 2 |
| GO:0072089 | stem cell proliferation | 2/9 | 121/18614 | 1.46E-03 | 0.06931282 | 0.040077094 | PRRX1/FBLN1 | 2 |
| GO:0030198 | extracellular matrix organization | 2/9 | 314/18614 | 0.009442712 | 0.080149586 | 0.046342978 | HMCN1/FBLN1 | 2 |
| GO:0043062 | extracellular structure organization | 2/9 | 315/18614 | 0.00950066 | 0.080149586 | 0.046342978 | HMCN1/FBLN1 | 2 |
| GO:0045229 | external encapsulating structure organization | 2/9 | 317/18614 | 0.009617042 | 0.080149586 | 0.046342978 | HMCN1/FBLN1 | 2 |
| GO:0031589 | cell-substrate adhesion | 2/9 | 359/18614 | 0.012209165 | 0.080149586 | 0.046342978 | ITGA11/FBLN1 | 2 |
| GO:0007018 | microtubule-based movement | 2/9 | 417/18614 | 0.016240581 | 0.082200506 | 0.047528832 | DST/STARD9 | 2 |
| GO:0042060 | wound healing | 2/9 | 439/18614 | 0.017902212 | 0.082200506 | 0.047528832 | DST/FBLN1 | 2 |
| GO:0006869 | lipid transport | 2/9 | 447/18614 | 0.018524058 | 0.082200506 | 0.047528832 | ABCA6/ABCA8 | 2 |

**Supplementary Table 6D: PathDIP pathways enriched in the upregulated differentially expressed genes from fibroblasts of obese BMI (n=8) vs normal BMI (n=7) KOA-IFPs.** Pathways were filtered by adjusted p < 0.05.

| Pathway Source | Pathway Name | p-value | q-value (FDR: BH-method) | q-value (Bonferroni) | Ratio | Query mapped | Pathway size | Type | Category |
|---|---|---|---|---|---|---|---|---|---|
| KEGG | ABC transporters | 9.90E-05 | 0.000297 | 0.000594122 | 0.4 | 2 | 45 | Environmental information processing | Membrane transport |
| REACTOME | Extracellular matrix organization | 9.24E-05 | 0.000555 | 0.000554561 | 0.6 | 3 | 300 | Cellular processes and organization | Cellular community |
| REACTOME | ABC-family proteins mediated transport | 0.0005211 | 0.001042 | 0.00312661 | 0.4 | 2 | 103 | Environmental information processing | Membrane transport |
| ACSN2 | ECM | 0.000791 | 0.001186 | 0.00474593 | 0.4 | 2 | 127 | Cellular processes and organization | Cellular community |
| ACSN2 | EMT REGULATORS | 0.0166506 | 0.019981 | 0.0999036 | 0.4 | 2 | 599 | Environmental information processing | Signal transduction |
| REACTOME | Transport of small molecules | 0.0239292 | 0.023929 | 0.143575 | 0.4 | 2 | 726 | Environmental information processing | Membrane transport |

**Supplementary Table 6E: Gene ontology biological processes enriched in the downregulated differentially expressed genes from fibroblasts of obese BMI (n=8) vs normal BMI (n=7) KOA-IFPs.** Biological processes were filtered by adjusted p < 0.05.

| ID | Description | GeneRatio | BgRatio | pvalue | p.adjust | qvalue | geneID | Count |
|---|---|---|---|---|---|---|---|---|
| GO:0071634 | regulation of transforming growth factor beta production | 2/11 | 40/18614 | 0.00024463 | 0.042493668 | 0.026111525 | FN1/ITGB8 | 2 |
| GO:0071604 | transforming growth factor beta production | 2/11 | 43/18614 | 0.000282933 | 0.042493668 | 0.026111525 | FN1/ITGB8 | 2 |
| GO:0002274 | myeloid leukocyte activation | 3/11 | 240/18614 | 0.000323556 | 0.042493668 | 0.026111525 | CLU/FN1/ITGB8 | 3 |
| GO:0061077 | chaperone-mediated protein folding | 2/11 | 69/18614 | 0.000728925 | 0.07179911 | 0.044119143 | CLU/FKBP5 | 2 |

**Supplementary Table 6F: PathDIP pathways enriched in the downregulated differentially expressed genes from fibroblasts of obese BMI (n=8) vs normal BMI (n=7) KOA-IFPs.** Pathways were filtered by adjusted p < 0.05.

| Pathway Source | Pathway Name | p-value | q-value (FDR: BH-method) | q-value (Bonferroni) | Ratio | Query mapped | Pathway size | Type | Category |
|---|---|---|---|---|---|---|---|---|---|
| REACTOME | Response to elevated platelet cytosolic Ca2+ | 9.38E-05 | 0.0006569 | 0.00262769 | 0.3 | 3 | 133 | Organismal systems | Immune system |
| REACTOME | Platelet degranulation | 8.37E-05 | 0.0007815 | 0.00234464 | 0.3 | 3 | 128 | Organismal systems | Immune system |
| ACSN2 | ECM | 8.18E-05 | 0.0011453 | 0.00229056 | 0.3 | 3 | 127 | Cellular processes and organization | Cellular community |
| REACTOME | Molecules associated with elastic fibres | 0.0002976 | 0.0016667 | 0.00833358 | 0.2 | 2 | 37 | Cellular processes and organization | Cellular community |
| WikiPathways | miR-509-3p alteration of YAP1/ECM axis | 6.89E-05 | 0.0019283 | 0.00192832 | 0.2 | 2 | 18 | Cellular processes and organization | Cellular community |
| REACTOME | Elastic fibre formation | 0.0004216 | 0.0019677 | 0.0118059 | 0.2 | 2 | 44 | Cellular processes and organization | Cellular community |
| REACTOME | Platelet activation, signaling and aggregation | 0.0006912 | 0.002765 | 0.019355 | 0.3 | 3 | 262 | Organismal systems | Immune system |
| ACSN2 | CELL MATRIX ADHESIONS | 0.0008918 | 0.0031212 | 0.0249694 | 0.2 | 2 | 64 | Cellular processes and organization | Cellular community |
| WikiPathways | Focal adhesion: PI3K-Akt-mTOR-signaling pathway | 0.0010545 | 0.0032807 | 0.029526 | 0.3 | 3 | 303 | Cellular processes and organization | Cellular community |
| KEGG | ECM-receptor interaction | 0.001678 | 0.0036142 | 0.0469843 | 0.2 | 2 | 88 | Cellular processes and organization | Cellular community |
| KEGG | PI3K-Akt signaling pathway | 0.0016121 | 0.0037615 | 0.0451382 | 0.3 | 3 | 351 | Environmental information processing | Signal transduction |
| REACTOME | Integrin cell surface interactions | 0.0015667 | 0.003988 | 0.0438679 | 0.2 | 2 | 85 | Cellular processes and organization | Cellular community |
| WikiPathways | PI3K-Akt signaling pathway | 0.0014461 | 0.004049 | 0.0404902 | 0.3 | 3 | 338 | Environmental information processing | Signal transduction |
| KEGG | Estrogen signaling pathway | 0.004182 | 0.008364 | 0.117097 | 0.2 | 2 | 137 | Organismal systems | Endocrine system |
| Panther_Pathway | Integrin signalling pathway | 0.0057605 | 0.0107529 | 0.161294 | 0.2 | 2 | 165 | Environmental information processing | Signal transduction |
| ACSN2 | EMT REGULATORS | 0.0073891 | 0.0129308 | 0.206893 | 0.3 | 3 | 599 | Environmental information processing | Signal transduction |
| KEGG | Focal adhesion | 0.0084417 | 0.0131315 | 0.236366 | 0.2 | 2 | 201 | Cellular processes and organization | Cellular community |
| WikiPathways | Focal adhesion | 0.0082004 | 0.0135066 | 0.229611 | 0.2 | 2 | 198 | Cellular processes and organization | Cellular community |
| KEGG | Regulation of actin cytoskeleton | 0.0098702 | 0.0145455 | 0.276364 | 0.2 | 2 | 218 | Organismal systems | Cell motility |
| REACTOME | Hemostasis | 0.0113846 | 0.0159384 | 0.318769 | 0.3 | 3 | 701 | Organismal systems | Immune system |
| WikiPathways | IL-18 signaling pathway | 0.0151736 | 0.0202315 | 0.424861 | 0.2 | 2 | 272 | Organismal systems | Immune system |
| REACTOME | Extracellular matrix organization | 0.018143 | 0.0220871 | 0.508004 | 0.2 | 2 | 300 | Cellular processes and organization | Cellular community |
| REACTOME | Metabolism of carbohydrates | 0.0176885 | 0.0225126 | 0.495278 | 0.2 | 2 | 296 | Metabolism | Carbohydrate metabolism |
