## Supplementary Table 7 for "Cell and Transcriptomic Diversity of Infrapatellar Fat Pad during Knee Osteoarthritis"

**Supplementary Table 7A: Analysis of metabolites detected in supernatents from naive fibroblast cultures of obese BMI (n=5) compared to normal BMI (n=5) KOA-IFPs.** Out of 646 metabolites targeted, 445 metabolites were detected. Metabolites with no variance or having extreme outliers were removed prior to statistical analysis. Data are arranged by increasing p-value.

| Metabolite | Fold.Change | log2FC | t.stat | p.value |
|---|---|---|---|---|
| TG(50:2) | 3.4152 | 1.772 | -2.9382 | 0.018762 |
| TG(54:2) | 9.8011 | 3.2929 | -2.5911 | 0.032057 |
| C18:2 | 4.356 | 2.123 | -2.4894 | 0.037557 |
| Choline | 1.0586 | 0.0822 | -2.449 | 0.039997 |
| Homoarginine | 0.21642 | -2.2081 | 2.3365 | 0.047678 |
| TG(48:2) | 8.1721 | 3.0307 | -2.2471 | 0.054813 |
| CE(18:2) | 2.3733 | 1.2469 | -2.0075 | 0.079589 |
| Fumaric acid | 0.28954 | -1.7882 | 1.8584 | 0.10017 |
| cis-4-Hydroxyproline | 1.2563 | 0.32913 | -1.8312 | 0.10445 |
| trans-4-Hydroxyproline | 1.2563 | 0.32913 | -1.8312 | 0.10445 |
| Methylhistidine | 1.3617 | 0.4454 | -1.8199 | 0.10627 |
| N-Acetyl-Histidine | 4.9311 | 2.3019 | -1.7991 | 0.1097 |
| C3:1 | 2.6267 | 1.3933 | -1.7642 | 0.11571 |
| 3-Hydroxyisovaleric acid | 1.2466 | 0.31797 | -1.7267 | 0.1225 |
| DG(32:1) | 0.12906 | -2.9539 | 1.6703 | 0.1334 |
| TG(51:4) | 0.3669 | -1.4466 | 1.6537 | 0.13679 |
| TG(54:7) | 3.4954 | 1.8055 | -1.6514 | 0.13727 |
| 2,5-Furandicarboxylic acid | 0.39911 | -1.3251 | 1.6278 | 0.14223 |
| Cer(40:1) | 0.23105 | -2.1137 | 1.5775 | 0.15333 |
| CE(20:5) | 0.29819 | -1.7457 | 1.5563 | 0.15825 |
| N2-Acetyl-Ornithine | 1.1786 | 0.23703 | -1.5522 | 0.15922 |
| Succinic acid | 1.3293 | 0.41067 | -1.4724 | 0.17912 |
| Cer(43:1) | 0.16423 | -2.6062 | 1.436 | 0.18894 |
| LysoPC a C18:0 | 1.1353 | 0.18313 | -1.4045 | 0.19778 |
| Caprylic acid | 8.1017 | 3.0182 | -1.3472 | 0.21484 |
| Ethanolamine | 2.0596 | 1.0423 | -1.3405 | 0.21689 |
| C18:1OH | 11.401 | 3.5111 | -1.3352 | 0.21856 |
| PC aa C38:0 | 0.42547 | -1.2329 | 1.3263 | 0.22136 |
| 2-hydroxyglutaric acid | 2.0034 | 1.0024 | -1.3035 | 0.22865 |
| 2-Hydroxyisobutyric acid | 1.3636 | 0.44746 | -1.2409 | 0.24979 |
| Indolelactic acid | 3.2875 | 1.717 | -1.2212 | 0.25677 |
| CE(18:1) | 1.1901 | 0.25108 | -1.212 | 0.2601 |
| LysoPC a C28:0 | 1.314 | 0.39396 | -1.19 | 0.26817 |
| TG(46:2) | 7.5193 | 2.9106 | -1.1589 | 0.27992 |
| N1-Acetylspermidine | 1.9138 | 0.93646 | -1.1545 | 0.28163 |
| Acetoacetic acid | 2.0347 | 1.0248 | -1.1535 | 0.282 |
| Citric acid | 0.63146 | -0.66324 | 1.1521 | 0.28256 |
| Deoxyadenosine | 1.812 | 0.85762 | -1.1322 | 0.29035 |
| CE(17:0) | 1.812 | 0.85762 | -1.1322 | 0.29035 |
| DG(35:1) | 1.812 | 0.85762 | -1.1322 | 0.29035 |
| DG(36:3) | 1.812 | 0.85762 | -1.1322 | 0.29035 |
| TG(44:2) | 1.812 | 0.85762 | -1.1322 | 0.29035 |
| TG(44:4) | 1.812 | 0.85762 | -1.1322 | 0.29035 |
| TG(50:3) | 1.812 | 0.85762 | -1.1322 | 0.29035 |
| TG(51:2) | 1.812 | 0.85762 | -1.1322 | 0.29035 |
| TG(56:7) | 1.812 | 0.85762 | -1.1322 | 0.29035 |
| CE(16:1) | 0.18675 | -2.4208 | 1.1133 | 0.29791 |
| Quinolinic acid | 1.8352 | 0.87594 | -1.1058 | 0.30096 |
| CE(22:5) | 1.8352 | 0.87594 | -1.1058 | 0.30096 |
| Guanine | 0.6007 | -0.73527 | 1.1035 | 0.30187 |
| Glutaric acid | 1.8389 | 0.87885 | -1.1018 | 0.30259 |
| C16:1OH | 1.8389 | 0.87885 | -1.1018 | 0.30259 |
| HexCer(41:2) | 0.60698 | -0.72027 | 1.0865 | 0.30889 |
| CE(22:6) | 1.8799 | 0.91068 | -1.0703 | 0.31573 |
| N-Acetylputrescine | 1.886 | 0.91535 | -1.0559 | 0.32185 |
| SMOH C22:1 | 1.886 | 0.91535 | -1.0559 | 0.32185 |
| TG(56:8) | 1.886 | 0.91535 | -1.0559 | 0.32185 |
| TG(54:6) | 1.8938 | 0.92131 | -1.0491 | 0.3248 |
| Isovaleric acid | 3.3777 | 1.7561 | -1.0466 | 0.32586 |
| Guanidinopropionic acid | 1.2273 | 0.29545 | -1.033 | 0.33182 |
| C4OH | 1.4838 | 0.56929 | -1.0148 | 0.3399 |
| N-Acetyl-Glutamine | 1.2718 | 0.34686 | -0.99394 | 0.34936 |
| alpha-Aminoadipic acid | 0.60194 | -0.73231 | 0.98866 | 0.35179 |
| DG(34:2) | 0.60194 | -0.73231 | 0.98866 | 0.35179 |
| HexCer(44:1) | 0.60194 | -0.73231 | 0.98866 | 0.35179 |
| SMOH C14:1 | 0.60194 | -0.73231 | 0.98866 | 0.35179 |
| C12DC | 0.60194 | -0.73231 | 0.98866 | 0.35179 |
| Homocitrulline | 1.1648 | 0.22004 | -0.94593 | 0.37188 |
| Hypoxanthine | 1.0418 | 0.059112 | 0.93378 | 0.37774 |

| | | | | |
|---|---|---|---|---|
| N1-Acetyl-Lysine + N6-Acetyl-Lysine | 1.0842 | 0.11666 | -0.93008 | 0.37954 |
| 2-Hydroxybutyric acid | 0.91976 | -0.12068 | 0.91034 | 0.38925 |
| Putrescine | 0.50922 | -0.97364 | 0.90818 | 0.39032 |
| Phenylalanine | 0.90097 | -0.15046 | 0.89989 | 0.39446 |
| Creatinine | 1.096 | 0.13222 | -0.88466 | 0.40215 |
| CE(22:2) | 0.56767 | -0.81687 | 0.88349 | 0.40274 |
| TG(49:2) | 0.56767 | -0.81687 | 0.88349 | 0.40274 |
| TG(53:3) | 0.56767 | -0.81687 | 0.88349 | 0.40274 |
| 3-Hydroxybutyric acid | 0.79919 | -0.32339 | 0.84147 | 0.42452 |
| 3-Hydroxyisobutyric acid | 0.79919 | -0.32339 | 0.84147 | 0.42452 |
| alpha-Ketoglutaric acid | 1.4281 | 0.51412 | -0.8332 | 0.4289 |
| TG(50:0) | 0.54577 | -0.87364 | 0.83215 | 0.42946 |
| Butyric acid + Isobutyric acid | 0.64899 | -0.62373 | 0.8303 | 0.43045 |
| C14:1OH | 0.48637 | -1.0399 | 0.82408 | 0.43377 |
| C6 | 1.0314 | 0.044605 | -0.82363 | 0.43401 |
| Asymmetric dimethylarginine + Symmetric dimethylarginine | 0.51568 | -0.95545 | 0.80644 | 0.4433 |
| TG(48:1) | 2.7498 | 1.4593 | -0.79878 | 0.44748 |
| N-Acetyl-Glycine | 1.636 | 0.71018 | -0.79569 | 0.44918 |
| LysoPC a C20:3 | 1.0293 | 0.041708 | -0.75362 | 0.47267 |
| TG(50:4) | 2.7493 | 1.4591 | -0.74541 | 0.47735 |
| Glutamine | 0.95772 | -0.062331 | 0.72069 | 0.49162 |
| Agmatine | 1.5319 | 0.61534 | -0.70873 | 0.49862 |
| Cadaverine | 1.5153 | 0.59961 | -0.69042 | 0.50946 |
| Serine | 1.0283 | 0.040312 | -0.67291 | 0.51997 |
| HexCer(40:2) | 0.59938 | -0.73846 | -0.66416 | 0.52526 |
| Betaine | 0.11932 | -3.0671 | 0.65964 | 0.52802 |
| Propionic acid | 0.67495 | -0.56715 | 0.65561 | 0.53047 |
| Norepinephrine | 1.3795 | 0.46416 | -0.65165 | 0.53289 |
| Lactic acid | 1.0536 | 0.075323 | -0.65039 | 0.53367 |
| Cer(38:0) | 0.048923 | -4.3533 | 0.63777 | 0.54144 |
| Glyceric acid | 1.2536 | 0.32605 | -0.63691 | 0.54198 |
| 2-Hydroxyisovaleric acid | 0.096946 | -3.3667 | 0.60179 | 0.56398 |
| C5 | 1.3101 | 0.38965 | -0.59343 | 0.56929 |
| SMOH C16:1 | 0.11482 | -3.1226 | 0.58497 | 0.57469 |
| Tyrosine | 1.0242 | 0.034497 | -0.58477 | 0.57482 |
| Methylmalonic acid | 1.3206 | 0.40117 | -0.57949 | 0.57821 |
| Ornithine | 0.76545 | -0.38562 | 0.57791 | 0.57922 |
| TG(52:5) | 1.4251 | 0.51104 | -0.56524 | 0.58741 |
| Cytosine | 1.4199 | 0.50583 | -0.55034 | 0.59711 |
| alpha-Aminobutyric acid | 5.3551 | 2.4209 | -0.54885 | 0.59809 |
| Malonic acid | 0.70862 | -0.49691 | 0.54427 | 0.60109 |
| C16OH | 0.12507 | -2.9992 | 0.54208 | 0.60253 |
| 3,4-Dihydroxybutyric acid | 0.72481 | -0.46432 | 0.53198 | 0.60919 |
| SM C18:1 | 0.15289 | -2.7094 | 0.49496 | 0.63393 |
| Methionine sulfoxide | 0.77671 | -0.36456 | 0.48408 | 0.6413 |
| C14:2 | 2.7212 | 1.4442 | -0.45595 | 0.66055 |
| Threonine | 1.0202 | 0.028889 | -0.4546 | 0.66148 |
| TG(51:1) | 1.2665 | 0.3408 | -0.44887 | 0.66543 |
| HexCer(34:2) | 0.61535 | -0.70052 | -0.44411 | 0.66873 |
| CE(15:0) | 2.4907 | 1.3165 | -0.41855 | 0.68655 |
| Lysine | 1.01 | 0.01431 | -0.40073 | 0.69911 |
| Taurine | 1.1172 | 0.15992 | -0.39259 | 0.70488 |
| C14:2OH | 2.0815 | 1.0576 | -0.38813 | 0.70804 |
| SM C16:1 | 0.45945 | -1.122 | -0.38506 | 0.71023 |
| SM C16:0 | 0.32719 | -1.6118 | -0.37137 | 0.72 |
| TG(49:1) | 0.95379 | -0.068253 | 0.35291 | 0.73327 |
| Pyruvic acid | 1.0285 | 0.040563 | -0.34556 | 0.73859 |
| TG(48:3) | 1.6825 | 0.75057 | -0.33791 | 0.74412 |
| 5-Oxoproline | 1.0126 | 0.018071 | -0.31797 | 0.75865 |
| Benzoic acid | 0.77206 | -0.37321 | 0.30762 | 0.76623 |
| SM C24:0 | 0.033318 | -4.9076 | 0.30612 | 0.76733 |
| TG(51:3) | 1.4796 | 0.56524 | -0.28973 | 0.77939 |
| CE(20:4) | 0.57867 | -0.78918 | 0.28858 | 0.78024 |
| HexCer(38:2) | 0.58473 | -0.77415 | -0.28154 | 0.78544 |
| Cer(41:1) | 1.4415 | 0.52756 | -0.27888 | 0.78741 |
| Asparagine | 0.77595 | -0.36597 | 0.25511 | 0.80507 |
| Valine | 1.0067 | 0.0096931 | -0.25048 | 0.80853 |
| TG(50:1) | 1.3252 | 0.40623 | -0.24107 | 0.81557 |
| Caproic acid | 1.768 | 0.82215 | -0.24013 | 0.81627 |
| Sarcosine+beta-Alanine | 1.0046 | 0.0065941 | -0.23169 | 0.82259 |
| Histidine | 1.0063 | 0.0090284 | -0.22266 | 0.82938 |
| Cer(42:0) | 1.2735 | 0.34882 | -0.22126 | 0.83043 |

| | | | | |
|---|---|---|---|---|
| alpha-Ketoisovaleric acid | 0.9645 | -0.052152 | 0.21134 | 0.8379 |
| Alanine | 1.029 | 0.041269 | -0.20448 | 0.84308 |
| N-Acetyl-Alanine | 0.63363 | -0.65829 | 0.19883 | 0.84735 |
| Inosine | 0.66858 | -0.58084 | -0.17775 | 0.86334 |
| Arginine | 1.0036 | 0.0051482 | -0.1675 | 0.87113 |
| C12:1 | 0.76051 | -0.39496 | 0.16517 | 0.87291 |
| CE(20:1) | 0.60537 | -0.72412 | 0.15897 | 0.87763 |
| Methionine | 1.0029 | 0.0042201 | -0.15085 | 0.88383 |
| PC aa C26:0 | 0.14241 | -2.8119 | 0.14768 | 0.88625 |
| 2-oxoisocaproic acid | 0.94885 | -0.075744 | 0.1456 | 0.88784 |
| N-Acetyl-Serine | 0.81869 | -0.28861 | 0.13992 | 0.89218 |
| 1-Methylnicotinamide | 0.88948 | -0.16897 | 0.12307 | 0.90509 |
| TG(53:4) | 1.2629 | 0.33677 | 0.1222 | 0.90575 |
| Glutamic acid | 0.73579 | -0.44263 | 0.10946 | 0.91553 |
| Hippuric acid | 1.0283 | 0.040267 | -0.090161 | 0.93038 |
| 5-Hydroxylysine | 1.0677 | 0.094457 | -0.078947 | 0.93901 |
| Leucine+Isoleucine | 1.0044 | 0.0063113 | -0.07692 | 0.94058 |
| Proline | 0.92132 | -0.11822 | 0.073331 | 0.94334 |
| Kynurenine | 1.0356 | 0.050513 | -0.069477 | 0.94631 |
| Malic acid | 0.90932 | -0.13714 | 0.064322 | 0.95029 |
| Hex2Cer(36:1) | 0.84163 | -0.24874 | 0.062434 | 0.95175 |
| PC aa C36:0 | 1.0113 | 0.016247 | -0.062397 | 0.95178 |
| Tryptophan | 0.99967 | -0.000473 | -0.060808 | 0.953 |
| Carnosine | 0.63028 | -0.66594 | -0.058186 | 0.95503 |
| Glycine | 0.99642 | -0.005168 | 0.029338 | 0.97731 |
| C5DC | 0.95996 | -0.05896 | 0.024612 | 0.98097 |
| PC aa C38:3 | 0.98105 | -0.027608 | -0.008392 | 0.99351 |

**Supplementary Table 7B: Analysis of the change in metabolites in supernatants of fibroblasts from obese BMI (n=5) vs normal BMI (n=5) KOA-IFPs induced by TGFβ compared to vehicle treatment.** Out of 646 metabolites targeted, 445 metabolites were detected. Metabolites with no variance or having extreme outliers were removed prior to statistical analysis. Data are arranged by increasing p-value.

| Metabolite | change in log2FC | AveExpr | t | P.Value |
|---|---|---|---|---|
| TG(52:5) | 1.6895 | 2.17E-16 | 2.8324 | 0.014864 |
| CE(17:0) | -1.9679 | 1.60E-16 | -2.7271 | 0.018095 |
| PC aa C26:0 | -1.6721 | -1.58E-16 | -2.5731 | 0.024098 |
| Ethanolamine | 1.0291 | 1.93E-16 | 2.4253 | 0.031661 |
| CE(20:1) | -1.4141 | 5.69E-17 | -2.3719 | 0.034919 |
| Homocitrulline | 1.1228 | 2.22E-17 | 2.352 | 0.036213 |
| C16:1OH | 0.97545 | -3.86E-16 | 2.2327 | 0.044997 |
| Hex2Cer(34:1) | 0.99706 | -2.29E-17 | 2.2319 | 0.045062 |
| alpha-Ketoglutaric acid | -0.99375 | 2.65E-17 | -2.2129 | 0.046631 |
| Uric acid | 1.0943 | 1.47E-17 | 2.1399 | 0.053185 |
| PC aa C38:0 | 1.1424 | 2.50E-17 | 2.0938 | 0.057753 |
| SM C24:0 | -1.6769 | 1.39E-18 | -2.0089 | 0.067141 |
| N-Acetylputrescine | 1.1792 | -3.61E-17 | 1.8683 | 0.085861 |
| TG(51:1) | -0.89487 | -8.40E-17 | -1.8573 | 0.087503 |
| alpha-Aminoadipic acid | 0.74797 | -2.14E-16 | 1.8215 | 0.09307 |
| Methylmalonic acid | 0.73575 | 7.81E-18 | 1.8197 | 0.093359 |
| TG(50:2) | 1.0612 | -8.33E-17 | 1.8175 | 0.09371 |
| Norepinephrine | 0.83835 | -3.75E-17 | 1.8005 | 0.096478 |
| TG(46:2) | 0.75184 | 2.75E-16 | 1.797 | 0.09705 |
| DG(32:1) | 0.73199 | -6.34E-16 | 1.7615 | 0.1031 |
| Phenylalanine | -0.4537 | -7.05E-17 | -1.7147 | 0.1116 |
| C18:1 | 0.81971 | 6.23E-16 | 1.7113 | 0.11225 |
| SMOH C14:1 | -0.98669 | -2.78E-18 | -1.6606 | 0.12219 |
| C5DC | 0.64873 | -5.13E-17 | 1.6291 | 0.12876 |
| C3:1 | 0.72451 | 2.01E-17 | 1.6163 | 0.13152 |
| SMOH C16:1 | -0.92405 | 2.78E-17 | -1.6097 | 0.13295 |
| CE(20:4) | 0.4931 | -2.26E-18 | 1.5699 | 0.14194 |
| 5-Hydroxylysine | -0.66237 | -1.88E-16 | -1.5278 | 0.152 |
| TG(54:5) | 0.51148 | -1.98E-16 | 1.4999 | 0.15902 |
| Methylamine | 0.51148 | 8.31E-17 | 1.4999 | 0.15902 |
| SM C26:0 | 0.51148 | 1.04E-16 | 1.4999 | 0.15902 |
| Benzoic acid | 0.63606 | 3.40E-17 | 1.4478 | 0.1728 |
| HexCer(40:2) | 1.1549 | 2.50E-16 | 1.443 | 0.17414 |
| DG(34:2) | -0.85698 | -3.41E-16 | -1.4203 | 0.1805 |
| SM C16:0 | -0.87173 | 4.61E-17 | -1.4154 | 0.18192 |
| TG(50:0) | 0.50928 | -4.28E-16 | 1.4091 | 0.18372 |
| DG(36:1) | 0.50928 | 4.04E-16 | 1.4091 | 0.18372 |
| TG(53:5) | 0.50928 | -3.20E-16 | 1.4091 | 0.18372 |
| Hex2Cer(38:1) | 0.5144 | 3.82E-18 | 1.4007 | 0.18616 |
| DG(35:1) | 0.5144 | -4.82E-16 | 1.4007 | 0.18616 |
| Deoxyadenosine | 0.51272 | -8.67E-17 | 1.3979 | 0.18699 |
| HexCer(36:1) | 0.49459 | 4.12E-16 | 1.3922 | 0.18866 |
| Methionine sulfoxide | -0.58222 | -1.11E-17 | -1.3649 | 0.19687 |
| TG(50:3) | 0.48777 | -5.53E-17 | 1.3418 | 0.20404 |
| TG(53:3) | 0.48777 | -1.28E-16 | 1.3418 | 0.20404 |
| TG(54:1) | 0.48777 | -2.31E-16 | 1.3418 | 0.20404 |
| C14:2OH | -0.73187 | -4.30E-17 | -1.3121 | 0.21357 |
| Butyric acid + Isobutyric acid | 0.64938 | 1.28E-16 | 1.3006 | 0.21738 |
| Propionic acid | 0.42026 | 2.05E-17 | 1.2991 | 0.21786 |
| TG(48:1) | -0.78826 | -1.94E-17 | -1.2944 | 0.21944 |
| N1-Acetyl-Lysine + N6-Acetyl-Lysine | -0.61219 | -1.15E-17 | -1.2843 | 0.22283 |
| Carnosine | -0.56366 | 1.23E-17 | -1.273 | 0.22669 |
| TG(50:4) | -0.68938 | -1.14E-16 | -1.2462 | 0.23603 |
| N2-Acetyl-Ornithine | 0.37765 | 2.13E-16 | 1.2393 | 0.2385 |
| Hippuric acid | 0.81127 | 6.94E-17 | 1.2358 | 0.23974 |
| Urea | 0.51336 | -1.44E-17 | 1.2042 | 0.25128 |
| Dopamine | 0.51336 | 1.91E-16 | 1.2042 | 0.25128 |
| Epinephrine | 0.51336 | -3.26E-16 | 1.2042 | 0.25128 |
| 3-Hydroxyisobutyric acid | 0.43785 | -1.54E-16 | 1.1998 | 0.25295 |
| 3-Hydroxybutyric acid | 0.43785 | 6.84E-17 | 1.1998 | 0.25295 |
| Cer(41:1) | -0.49653 | -1.39E-16 | -1.1982 | 0.25354 |
| C5OH | 0.46962 | 1.51E-16 | 1.182 | 0.25969 |
| SM C16:1 | -0.74603 | -5.07E-17 | -1.1609 | 0.26783 |
| PC aa C36:0 | 0.35304 | -8.33E-18 | 1.1565 | 0.26955 |
| TG(56:8) | -0.51515 | 1.64E-16 | -1.1495 | 0.27234 |
| PC aa C38:3 | 0.35054 | 1.80E-17 | 1.1439 | 0.27456 |
| SM C20:2 | 0.3389 | 3.71E-17 | 1.1418 | 0.27539 |

| | | | | |
|---|---|---|---|---|
| C18:1OH | 0.67533 | 4.45E-16 | 1.1274 | 0.28122 |
| 2-Hydroxyisobutyric acid | 0.5176 | -9.58E-17 | 1.0465 | 0.31557 |
| C18:2 | 0.44693 | -3.89E-17 | 1.0158 | 0.3294 |
| TG(56:7) | 0.48851 | -1.35E-16 | 1.0036 | 0.33502 |
| Guanine | 0.44523 | -4.44E-17 | 0.98388 | 0.34424 |
| LysoPC a C28:0 | 0.4548 | -8.19E-17 | 0.98147 | 0.34538 |
| Homoarginine | 0.53046 | -1.18E-17 | 0.97987 | 0.34613 |
| TG(53:4) | -0.64178 | -5.62E-17 | -0.95912 | 0.35607 |
| Fumaric acid | 0.50167 | 1.04E-18 | 0.94318 | 0.36385 |
| Isovaleric acid | -0.40581 | 9.16E-17 | -0.93192 | 0.36941 |
| Betaine | -0.47724 | -8.33E-18 | -0.87568 | 0.39807 |
| 5-Oxoproline | 0.25031 | -1.64E-16 | 0.86596 | 0.40318 |
| C5 | -0.72889 | -9.85E-17 | -0.85867 | 0.40704 |
| Cer(38:0) | 0.4914 | -6.28E-17 | 0.84996 | 0.41167 |
| CE(16:1) | -0.56753 | 7.63E-17 | -0.84813 | 0.41266 |
| 2-Hydroxybutyric acid | 0.29432 | 1.92E-16 | 0.84129 | 0.41633 |
| Hypoxanthine | 0.26825 | 1.53E-17 | 0.84084 | 0.41657 |
| Ornithine | 0.41468 | 5.12E-18 | 0.83138 | 0.4217 |
| PC aa C40:2 | -0.36573 | 2.28E-16 | -0.80386 | 0.43683 |
| C12DC | -0.2695 | -4.58E-17 | -0.80024 | 0.43884 |
| Putrescine | -0.41039 | 6.11E-17 | -0.79224 | 0.44331 |
| Succinic acid | 0.27432 | 6.18E-17 | 0.78963 | 0.44478 |
| CE(15:0) | -0.43619 | -8.60E-17 | -0.7859 | 0.44689 |
| Cer(40:1) | -0.55628 | 5.55E-17 | -0.78325 | 0.44838 |
| Cer(43:1) | -0.53821 | -1.85E-16 | -0.78196 | 0.44911 |
| CE(20:5) | -0.28543 | -5.04E-17 | -0.7548 | 0.46465 |
| Taurine | 0.44659 | 1.83E-16 | 0.75338 | 0.46548 |
| TG(56:9) | 0.36186 | 1.82E-16 | 0.747 | 0.46918 |
| CE(18:1) | -0.48649 | -4.51E-17 | -0.7286 | 0.47997 |
| Guanidinopropionic acid | 0.47018 | 5.97E-17 | 0.72013 | 0.48498 |
| Glutamic acid | 0.24678 | -4.16E-17 | 0.6998 | 0.49715 |
| Proline | 0.18265 | -6.07E-18 | 0.67983 | 0.50927 |
| Lactic acid | 0.192 | 3.99E-17 | 0.67397 | 0.51287 |
| Caprylic acid | 0.33959 | -1.55E-16 | 0.67366 | 0.51306 |
| TG(51:4) | -0.37265 | 2.95E-16 | -0.66986 | 0.51539 |
| Acetoacetic acid | 0.3179 | -4.89E-17 | 0.66848 | 0.51624 |
| Glutaric acid | 0.33956 | 1.87E-16 | 0.66187 | 0.52033 |
| Asparagine | -0.26226 | -1.80E-17 | -0.65315 | 0.52575 |
| HexCer(44:1) | 0.42219 | -7.77E-17 | 0.64766 | 0.52917 |
| 1-Methylnicotinamide | 0.20764 | 1.20E-16 | 0.64599 | 0.53022 |
| C4OH | -0.53797 | -9.16E-17 | -0.63616 | 0.5364 |
| Citric acid | -0.16994 | -2.08E-17 | -0.62345 | 0.54444 |
| C16OH | -0.39173 | 5.44E-16 | -0.60672 | 0.55513 |
| HexCer(34:2) | 0.35715 | -7.20E-17 | 0.59056 | 0.56557 |
| LysoPC a C18:0 | 0.14212 | 7.11E-17 | 0.56915 | 0.57956 |
| Malonic acid | 0.34023 | 5.83E-17 | 0.56673 | 0.58115 |
| HexCer(34:1) | -0.18568 | 2.48E-16 | -0.55286 | 0.59032 |
| TG(54:6) | -0.18568 | 5.64E-17 | -0.55286 | 0.59032 |
| HexCer(38:2) | 0.53525 | 1.67E-17 | 0.54811 | 0.59348 |
| N-Acetyl-Serine | -0.24302 | -4.20E-17 | -0.54233 | 0.59733 |
| Cer(42:0) | 0.28986 | 7.92E-16 | 0.54073 | 0.5984 |
| Lysine | 0.16103 | -2.09E-16 | 0.53966 | 0.59912 |
| C6 | -0.16766 | 4.72E-17 | -0.5298 | 0.60574 |
| SM C24:1 | -0.17055 | 5.67E-17 | -0.51953 | 0.61267 |
| LysoPC a C20:4 | -0.16425 | 7.11E-17 | -0.5195 | 0.61268 |
| LysoPC a C18:1 | -0.16365 | 1.18E-16 | -0.51786 | 0.61379 |
| Serine | 0.15366 | -3.76E-16 | 0.50983 | 0.61924 |
| Tyrosine | 0.15036 | 2.17E-16 | 0.46684 | 0.64882 |
| CE(22:6) | 0.24454 | -1.16E-16 | 0.4664 | 0.64913 |
| Glyceric acid | 0.23527 | -1.18E-16 | 0.46621 | 0.64926 |
| C14:1OH | -0.35147 | -1.30E-16 | -0.46522 | 0.64995 |
| DG(32:2) | -0.18414 | 5.34E-16 | -0.45442 | 0.65749 |
| 2-hydroxyglutaric acid | -0.18414 | 6.31E-17 | -0.45442 | 0.65749 |
| Valeric acid | -0.18414 | 2.78E-18 | -0.45442 | 0.65749 |
| C5:1 | -0.18414 | -5.90E-18 | -0.45442 | 0.65749 |
| Inosine | 0.19481 | -9.85E-17 | 0.44788 | 0.66207 |
| LysoPC a C20:3 | -0.12525 | -9.71E-18 | -0.4392 | 0.66817 |
| CE(22:5) | -0.16382 | 3.47E-18 | -0.43712 | 0.66964 |
| TG(49:1) | -0.16382 | 2.66E-16 | -0.43712 | 0.66964 |
| Alanine | 0.10994 | -4.53E-17 | 0.43453 | 0.67147 |
| Sarcosine+beta-Alanine | 0.18988 | 6.38E-17 | 0.42777 | 0.67625 |

| | | | | |
|---|---|---|---|---|
| 2-Hydroxyisovaleric acid | -0.16265 | 4.72E-16 | -0.4272 | 0.67666 |
| Guanosine | -0.16265 | 8.31E-17 | -0.4272 | 0.67666 |
| SMOH C22:1 | -0.16223 | 9.54E-16 | -0.42705 | 0.67677 |
| TG(50:1) | -0.25352 | 7.77E-17 | -0.42593 | 0.67756 |
| Agmatine | 0.20045 | -6.72E-18 | 0.42507 | 0.67817 |
| Cer(38:1) | -0.16317 | -4.23E-16 | -0.42373 | 0.67912 |
| TG(52:6) | -0.16317 | -1.90E-16 | -0.42373 | 0.67912 |
| TG(51:3) | -0.16023 | 1.81E-16 | -0.42103 | 0.68104 |
| Threonine | 0.13724 | -2.37E-16 | 0.41948 | 0.68214 |
| Tryptophan | 0.12135 | -1.26E-16 | 0.41285 | 0.68686 |
| CE(22:2) | 0.2283 | 6.42E-17 | 0.41087 | 0.68827 |
| Methylhistidine | 0.14584 | 1.37E-16 | 0.41075 | 0.68836 |
| C4:1 | -0.15916 | -2.36E-16 | -0.40913 | 0.68952 |
| Leucine+Isoleucine | 0.11282 | 1.43E-16 | 0.39614 | 0.69883 |
| TG(54:7) | 0.17817 | -1.26E-16 | 0.3804 | 0.71017 |
| Methionine | 0.10916 | -8.22E-17 | 0.37987 | 0.71056 |
| 3,4-Dihydroxybutyric acid | 0.17489 | -6.89E-17 | 0.36348 | 0.72245 |
| PC ae C42:1 | 0.15501 | 3.97E-16 | 0.36032 | 0.72475 |
| 2-oxoisocaproic acid | 0.11735 | -1.27E-17 | 0.35988 | 0.72507 |
| Choline | 0.099098 | -2.75E-17 | 0.34143 | 0.73857 |
| N-Acetyl-Alanine | -0.22844 | -5.97E-17 | -0.34015 | 0.73952 |
| Kynurenine | 0.16234 | -9.44E-17 | 0.33978 | 0.73979 |
| Arginine | 0.10077 | -8.07E-17 | 0.33388 | 0.74412 |
| Glycine | 0.098361 | 9.12E-17 | 0.33279 | 0.74493 |
| alpha-Ketoisovaleric acid | 0.10622 | 8.83E-18 | 0.33073 | 0.74645 |
| Cadaverine | -0.27783 | 2.27E-17 | -0.31991 | 0.75444 |
| TG(48:2) | 0.12946 | 1.09E-16 | 0.30899 | 0.76253 |
| N-Acetyl-Glutamine | -0.17248 | 8.31E-18 | -0.29481 | 0.77308 |
| Cytosine | 0.12564 | 1.25E-16 | 0.28812 | 0.77808 |
| Creatinine | 0.1314 | -1.11E-17 | 0.27311 | 0.78933 |
| TG(48:0) | 0.10113 | 8.61E-17 | 0.25777 | 0.80087 |
| C12:1 | -0.13187 | -1.67E-17 | -0.23984 | 0.81443 |
| Histidine | 0.072605 | -6.56E-17 | 0.23384 | 0.81898 |
| SM C18:1 | -0.11366 | -1.39E-16 | -0.22447 | 0.8261 |
| Asymmetric dimethylarginine + Symmetric dimethylarginine | 0.085686 | -1.34E-16 | 0.21297 | 0.83486 |
| CE(18:2) | -0.12264 | 5.57E-17 | -0.21106 | 0.83631 |
| TG(44:4) | 0.11911 | 5.83E-17 | 0.21075 | 0.83655 |
| Glutamine | 0.057404 | -7.21E-16 | 0.19946 | 0.84518 |
| 3-Hydroxyisovaleric acid | 0.092832 | 1.35E-16 | 0.18815 | 0.85385 |
| Malic acid | 0.049035 | -1.21E-16 | 0.1833 | 0.85756 |
| Pyruvic acid | 0.049562 | 3.80E-17 | 0.17858 | 0.86119 |
| Caproic acid | -0.059523 | -1.06E-16 | -0.17644 | 0.86284 |
| 2,5-Furandicarboxylic acid | 0.046556 | -4.28E-17 | 0.1725 | 0.86587 |
| Valine | 0.04366 | -1.34E-16 | 0.15754 | 0.87739 |
| TG(54:2) | -0.10773 | -2.64E-17 | -0.1547 | 0.87958 |
| DG(40:7) | 0.063566 | -4.81E-17 | 0.1471 | 0.88545 |
| cis-4-Hydroxyproline | 0.042862 | -4.10E-17 | 0.14557 | 0.88663 |
| trans-4-Hydroxyproline | 0.042862 | -4.10E-17 | 0.14557 | 0.88663 |
| HexCer(41:2) | -0.13849 | 1.20E-16 | -0.13798 | 0.8925 |
| N-Acetyl-Glycine | 0.034488 | 5.75E-17 | 0.093237 | 0.92723 |
| N1-Acetylspermidine | -0.018722 | -1.84E-16 | -0.072315 | 0.94352 |
| C14:2 | -0.033994 | -1.03E-16 | -0.055503 | 0.95663 |

**Supplementary Table 7C: Analysis of the change in metabolites in supernatents of fibroblasts from obese BMI (n=5) vs normal BMI (n=5) KOA-IFPs induced by TNFα compared to vehicle treatment.** Out of 646 metabolites targeted, 445 metabolites were detected. Metabolites with no variance or having extreme outliers were removed prior to statistical analysis. Data are arranged by increasing p-value.

| Metabolite | change in log2FC | AveExpr | t | P.Value |
|---|---|---|---|---|
| TG(54:7) | 1.4074 | -4.30E-17 | 3.4606 | 0.0038693 |
| TG(52:5) | 2.0463 | -9.37E-18 | 3.4136 | 0.0042469 |
| Serotonin | 1.1796 | 2.87E-17 | 3.2201 | 0.0062307 |
| SM C16:1 | -1.6502 | -1.21E-17 | -2.7168 | 0.016807 |
| 3-Hydroxybutyric acid | 1.0773 | 4.86E-18 | 2.5892 | 0.021545 |
| 3-Hydroxyisobutyric acid | 1.0773 | -9.25E-17 | 2.5892 | 0.021545 |
| SM C16:0 | -1.4904 | 2.12E-17 | -2.5554 | 0.023002 |
| Glutaric acid | 1.0206 | 1.35E-16 | 2.3982 | 0.031116 |
| SM C24:0 | -1.6688 | 7.01E-17 | -2.3728 | 0.032661 |
| N-Acetyl-Glycine | 1.1636 | 8.81E-17 | 2.3375 | 0.034932 |
| N-Acetylputrescine | 1.3111 | 1.71E-16 | 2.0885 | 0.055675 |
| TG(50:2) | 1.0581 | -2.22E-17 | 2.0304 | 0.06194 |
| TG(50:3) | 0.94021 | -1.18E-16 | 2.0293 | 0.062064 |
| SMOH C14:1 | -1.0698 | -3.61E-17 | -1.9872 | 0.067008 |
| Ethanolamine | -0.94675 | -1.19E-16 | -1.9672 | 0.069491 |
| SMOH C16:1 | -1.1212 | -6.52E-17 | -1.9072 | 0.07742 |
| Malonic acid | -0.93399 | 4.44E-17 | -1.869 | 0.082879 |
| CE(18:1) | 1.585 | -4.72E-17 | 1.8447 | 0.086527 |
| CE(16:1) | -0.92373 | -1.17E-16 | -1.8183 | 0.090671 |
| alpha-Ketoglutaric acid | -0.82238 | -6.17E-17 | -1.7991 | 0.093793 |
| Taurine | 0.79846 | -1.04E-16 | 1.7903 | 0.095245 |
| TG(48:3) | 0.73544 | -5.07E-17 | 1.7129 | 0.10898 |
| TG(54:5) | 0.73263 | -2.17E-16 | 1.7025 | 0.11095 |
| DG(36:3) | 0.77895 | -3.67E-16 | 1.6896 | 0.11343 |
| TG(44:2) | 0.71397 | 1.30E-19 | 1.6843 | 0.11447 |
| Hex2Cer(34:1) | 0.69058 | 1.32E-16 | 1.6729 | 0.11674 |
| TG(53:3) | 0.74507 | -1.84E-16 | 1.6685 | 0.11762 |
| TG(51:4) | -0.79024 | -1.32E-17 | -1.657 | 0.11996 |
| C4OH | 0.83718 | 9.99E-17 | 1.5914 | 0.13405 |
| Cer(42:0) | 0.76019 | -2.44E-16 | 1.5775 | 0.13721 |
| C16OH | 0.98412 | -2.87E-16 | 1.5153 | 0.15215 |
| CE(17:0) | 1.0457 | -1.61E-16 | 1.5078 | 0.15404 |
| N2-Acetyl-Ornithine | 0.51761 | 6.34E-17 | 1.4851 | 0.15988 |
| Putrescine | 0.64421 | -2.53E-16 | 1.4831 | 0.1604 |
| Methionine sulfoxide | -0.63008 | 2.78E-18 | -1.4581 | 0.16706 |
| C18:2 | 0.76258 | -2.90E-16 | 1.4467 | 0.17021 |
| Benzoic acid | 0.74755 | 4.51E-18 | 1.4445 | 0.1708 |
| Homocitrulline | 0.54106 | -5.83E-17 | 1.4242 | 0.17648 |
| SM C20:2 | 0.51241 | -2.60E-16 | 1.3599 | 0.19556 |
| N-Acetyl-Glutamine | -0.4951 | -1.01E-17 | -1.357 | 0.19646 |
| HexCer(41:2) | -0.86513 | 5.83E-17 | -1.3542 | 0.19732 |
| Propionic acid | 0.56615 | 6.11E-17 | 1.3419 | 0.20119 |
| PC aa C38:3 | 0.61999 | -6.66E-17 | 1.3414 | 0.20133 |
| Cer(41:1) | -0.50799 | 3.53E-16 | -1.3063 | 0.2127 |
| 3,4-Dihydroxybutyric acid | 0.48992 | -1.01E-16 | 1.2872 | 0.21909 |
| DG(32:2) | 0.48992 | -1.04E-16 | 1.2872 | 0.21909 |
| 2,5-Furandicarboxylic acid | 0.48456 | -1.13E-16 | 1.2533 | 0.2308 |
| TG(56:7) | 0.48456 | 2.47E-16 | 1.2533 | 0.2308 |
| Methylamine | 0.48456 | -1.69E-16 | 1.2533 | 0.2308 |
| SM C26:0 | 0.48456 | -1.67E-16 | 1.2533 | 0.2308 |
| SMOH C22:1 | -0.76057 | -4.82E-16 | -1.2457 | 0.23349 |
| CE(20:5) | 0.65447 | -5.83E-17 | 1.2424 | 0.23466 |
| TG(56:6) | 0.4745 | -7.61E-17 | 1.2372 | 0.23655 |
| N1-Acetyl-Lysine + N6-Acetyl-Lysine | -0.71613 | -4.43E-17 | -1.2254 | 0.24082 |
| 2-Hydroxybutyric acid | 0.5308 | -5.52E-17 | 1.217 | 0.24389 |
| Urea | 0.47497 | 4.36E-17 | 1.2137 | 0.24511 |
| SM C24:1 | 0.47497 | -4.83E-16 | 1.2137 | 0.24511 |
| Hex2Cer(32:1) | 0.47497 | 1.55E-16 | 1.2137 | 0.24511 |
| Methylmalonic acid | 0.46689 | -2.52E-18 | 1.2074 | 0.24747 |
| HexCer(36:1) | 0.46689 | 4.22E-16 | 1.2074 | 0.24747 |
| PC aa C38:0 | 0.58821 | -7.77E-17 | 1.1769 | 0.25903 |
| C14:2OH | -0.61036 | -3.33E-17 | -1.1765 | 0.25921 |
| CE(20:1) | -0.98265 | 1.33E-16 | -1.158 | 0.26643 |
| Norepinephrine | 0.62 | 1.39E-17 | 1.1444 | 0.2718 |
| C4:1 | -0.48007 | 2.64E-16 | -1.1408 | 0.27328 |
| PC aa C26:0 | -0.85503 | -8.33E-18 | -1.1228 | 0.28059 |
| 5-Hydroxylysine | -0.46406 | -1.85E-16 | -1.1087 | 0.28641 |

| | | | | |
|---|---|---|---|---|
| TG(49:2) | 0.48288 | -5.20E-17 | 1.1015 | 0.28941 |
| TG(52:4) | 0.48288 | 1.35E-16 | 1.1015 | 0.28941 |
| TG(44:1) | 0.48288 | -1.36E-16 | 1.1015 | 0.28941 |
| Indolelactic acid | 0.48288 | -2.76E-16 | 1.1015 | 0.28941 |
| CE(15:0) | 0.67596 | 2.18E-16 | 1.0873 | 0.29544 |
| Inosine | 0.50893 | 9.16E-17 | 1.0812 | 0.29805 |
| C12DC | 0.66188 | -1.28E-16 | 1.0809 | 0.29819 |
| DG(32:1) | 0.75978 | 4.66E-16 | 1.0534 | 0.31015 |
| TG(46:2) | 0.63428 | -4.16E-18 | 1.0422 | 0.31515 |
| Succinic acid | -0.37921 | -1.86E-17 | -1.0332 | 0.31918 |
| LysoPC a C28:0 | 0.35912 | -4.58E-17 | 1.0329 | 0.3193 |
| LysoPC a C18:0 | 0.72076 | -1.67E-17 | 1.0166 | 0.32674 |
| Hippuric acid | 0.54458 | -1.83E-16 | 1.0124 | 0.3287 |
| Isovaleric acid | 0.57103 | -1.14E-16 | 1.0029 | 0.3331 |
| LysoPC a C20:3 | -0.76937 | 2.78E-18 | -1.0012 | 0.33385 |
| TG(51:1) | -0.43266 | 5.35E-17 | -0.95735 | 0.35478 |
| Caprylic acid | -0.47768 | -1.08E-16 | -0.92822 | 0.36916 |
| Acetoacetic acid | -0.44173 | -3.28E-17 | -0.91552 | 0.37556 |
| Lactic acid | 0.2952 | 4.31E-16 | 0.89789 | 0.38456 |
| PC aa C36:0 | 0.31811 | -5.55E-18 | 0.89684 | 0.3851 |
| TG(56:8) | 0.59061 | -1.25E-16 | 0.89304 | 0.38706 |
| Proline | 0.27576 | 2.78E-18 | 0.8899 | 0.38869 |
| TG(54:6) | 0.30984 | -6.45E-17 | 0.87543 | 0.39625 |
| N-Acetyl-Serine | 0.42925 | 1.80E-16 | 0.84968 | 0.40992 |
| TG(54:2) | 0.55544 | 1.09E-16 | 0.83393 | 0.41845 |
| LysoPC a C20:4 | 0.39453 | 7.98E-18 | 0.83314 | 0.41888 |
| Homoarginine | 0.46517 | -5.97E-17 | 0.81969 | 0.42625 |
| C5DC | 0.2877 | -5.41E-17 | 0.81787 | 0.42725 |
| Cer(43:1) | -0.46512 | -4.17E-16 | -0.81769 | 0.42736 |
| HexCer(38:2) | 0.69537 | -7.29E-18 | 0.81323 | 0.42982 |
| CE(20:4) | 0.34787 | 1.94E-17 | 0.78632 | 0.4449 |
| Carnosine | -0.39744 | -3.34E-17 | -0.7686 | 0.45501 |
| Butyric acid + Isobutyric acid | 0.43019 | -3.14E-17 | 0.75092 | 0.46525 |
| TG(48:2) | 0.35295 | 5.55E-18 | 0.74394 | 0.46933 |
| Guanidinopropionic acid | 0.39178 | 1.64E-17 | 0.73507 | 0.47454 |
| Fumaric acid | 0.43466 | 3.12E-18 | 0.73385 | 0.47526 |
| Guanine | 0.26448 | 7.49E-17 | 0.72046 | 0.4832 |
| C18:1 | 0.43459 | -3.77E-16 | 0.71885 | 0.48416 |
| Methylhistidine | 0.30159 | -1.17E-16 | 0.71111 | 0.4888 |
| HexCer(34:2) | 0.43987 | 2.33E-16 | 0.70478 | 0.49261 |
| 5-Oxoproline | 0.22855 | 4.32E-17 | 0.69639 | 0.49768 |
| Phenylalanine | -0.26048 | 1.41E-16 | -0.69332 | 0.49955 |
| 2-oxoisocaproic acid | 0.24166 | 3.10E-17 | 0.67644 | 0.50988 |
| Asymmetric dimethylarginine + Symmetric dimethylarginine | -0.31099 | 3.82E-18 | -0.64102 | 0.53195 |
| HexCer(34:1) | -0.23444 | -2.55E-16 | -0.6379 | 0.53393 |
| Cer(38:0) | -0.23444 | -4.99E-16 | -0.6379 | 0.53393 |
| TG(53:4) | 0.30785 | 1.02E-16 | 0.63465 | 0.53598 |
| C3:1 | 0.32154 | -2.64E-17 | 0.62791 | 0.54026 |
| Betaine | 0.30178 | 5.48E-17 | 0.60124 | 0.55738 |
| LysoPC a C18:1 | 0.29911 | -1.46E-17 | 0.53301 | 0.60246 |
| HexCer(40:2) | -0.51505 | 2.48E-16 | -0.53196 | 0.60317 |
| C18:1OH | 0.30407 | -3.33E-17 | 0.5228 | 0.60936 |
| DG(36:2) | -0.20181 | 4.27E-16 | -0.52096 | 0.61061 |
| DG(40:7) | -0.2099 | -3.07E-16 | -0.5188 | 0.61208 |
| 2-Hydroxyisobutyric acid | 0.24714 | -5.55E-18 | 0.51857 | 0.61223 |
| 2-Hydroxyisovaleric acid | -0.20178 | -1.12E-16 | -0.5098 | 0.6182 |
| Guanosine | -0.20178 | 9.54E-19 | -0.5098 | 0.6182 |
| Cer(38:1) | -0.20105 | 1.08E-15 | -0.50693 | 0.62016 |
| Hex2Cer(36:1) | -0.20603 | 3.80E-16 | -0.50472 | 0.62168 |
| C16:1 | -0.20603 | -3.88E-16 | -0.50472 | 0.62168 |
| CE(22:0) | -0.20603 | -9.52E-17 | -0.50472 | 0.62168 |
| Citrulline | -0.20603 | 2.25E-16 | -0.50472 | 0.62168 |
| TG(51:2) | -0.20603 | -6.70E-17 | -0.50472 | 0.62168 |
| Ornithine | 0.18501 | 8.33E-18 | 0.48861 | 0.63275 |
| alpha-Ketoisovaleric acid | 0.18362 | -1.46E-17 | 0.48842 | 0.63288 |
| TG(48:0) | -0.2193 | 5.69E-17 | -0.47945 | 0.63909 |
| 1-Methylnicotinamide | 0.15902 | -2.88E-17 | 0.47428 | 0.64268 |
| Glutamic acid | 0.13854 | 3.82E-17 | 0.4472 | 0.66163 |
| Glyceric acid | -0.23009 | 1.62E-16 | -0.44536 | 0.66293 |
| cis-4-Hydroxyproline | -0.1376 | 2.57E-16 | -0.39994 | 0.69529 |

| | | | | |
|---|---|---|---|---|
| trans-4-Hydroxyproline | -0.1376 | 2.57E-16 | -0.39994 | 0.69529 |
| C5OH | 0.11629 | 2.26E-16 | 0.39605 | 0.69809 |
| Threonine | -0.13377 | -2.01E-16 | -0.39054 | 0.70207 |
| Cytosine | -0.17573 | -1.97E-16 | -0.35334 | 0.72914 |
| Sarcosine+beta-Alanine | -0.12529 | -5.55E-17 | -0.33849 | 0.74006 |
| Agmatine | 0.13376 | -2.74E-16 | 0.33357 | 0.74369 |
| Glutamine | -0.10679 | -2.91E-16 | -0.31637 | 0.75643 |
| Leucine+Isoleucine | 0.10074 | 2.05E-16 | 0.30824 | 0.76248 |
| TG(50:4) | -0.26714 | -6.38E-17 | -0.30591 | 0.76422 |
| Histidine | -0.10342 | -4.04E-17 | -0.30016 | 0.76851 |
| TG(54:1) | 0.16468 | -2.93E-16 | 0.29206 | 0.77456 |
| Valeric acid | 0.12395 | 7.91E-17 | 0.28996 | 0.77614 |
| Valine | -0.095897 | -9.69E-17 | -0.28855 | 0.77719 |
| CE(22:6) | 0.11648 | 6.87E-17 | 0.28596 | 0.77913 |
| Serine | -0.092583 | 3.46E-16 | -0.27209 | 0.78956 |
| PC aa C40:2 | 0.11918 | -4.67E-16 | 0.27162 | 0.78992 |
| Methionine | -0.091499 | 2.21E-16 | -0.26935 | 0.79162 |
| CE(18:2) | 0.20894 | -2.43E-17 | 0.26494 | 0.79495 |
| C6 | 0.11972 | 5.52E-17 | 0.26159 | 0.79748 |
| C16:1OH | 0.16171 | -3.22E-16 | 0.25533 | 0.80222 |
| TG(51:3) | 0.12438 | 5.52E-17 | 0.25215 | 0.80462 |
| Kynurenine | -0.096643 | -3.61E-17 | -0.23675 | 0.81631 |
| C12:1 | -0.11435 | -5.55E-18 | -0.23513 | 0.81754 |
| Deoxyadenosine | 0.10379 | 6.57E-17 | 0.23428 | 0.81819 |
| CE(22:2) | 0.093472 | -4.91E-17 | 0.22479 | 0.82542 |
| TG(50:1) | -0.129 | -7.77E-17 | -0.22291 | 0.82685 |
| Cer(40:1) | -0.17971 | -1.42E-16 | -0.22057 | 0.82864 |
| Pyruvic acid | -0.072483 | 1.62E-16 | -0.22005 | 0.82904 |
| Alanine | 0.073662 | -6.09E-17 | 0.21694 | 0.83142 |
| TG(56:9) | 0.093094 | -1.82E-16 | 0.21183 | 0.83532 |
| Citric acid | 0.073083 | 1.21E-18 | 0.20563 | 0.84007 |
| TG(44:4) | 0.10657 | -2.78E-18 | 0.19304 | 0.84973 |
| N1-Acetylspermidine | -0.07916 | -2.92E-16 | -0.17697 | 0.86209 |
| Arginine | -0.059177 | -1.21E-18 | -0.17288 | 0.86524 |
| N-Acetyl-Alanine | -0.051744 | -7.98E-17 | -0.15674 | 0.87771 |
| C14:1OH | -0.061159 | 1.24E-16 | -0.11663 | 0.90882 |
| Cadaverine | 0.082098 | -2.78E-17 | 0.1134 | 0.91134 |
| C5 | 0.060182 | 5.26E-17 | 0.11312 | 0.91156 |
| Asparagine | -0.049627 | -4.37E-17 | -0.11105 | 0.91317 |
| Choline | -0.034868 | -7.42E-18 | -0.10276 | 0.91963 |
| Glycine | -0.028929 | -2.69E-16 | -0.084232 | 0.93408 |
| Malic acid | -0.029329 | -4.22E-17 | -0.074766 | 0.94147 |
| 3-Hydroxyisovaleric acid | -0.022684 | 2.01E-16 | -0.063719 | 0.9501 |
| Tyrosine | -0.018043 | -1.28E-16 | -0.049347 | 0.96135 |
| Tryptophan | -0.013506 | -6.48E-17 | -0.039176 | 0.96931 |
| DG(35:1) | 0.015006 | 1.75E-16 | 0.033006 | 0.97414 |
| HexCer(44:1) | 0.014253 | -1.31E-16 | 0.02901 | 0.97727 |
| Creatinine | 0.013024 | 8.05E-17 | 0.027846 | 0.97818 |
| Lysine | -0.0085233 | 4.00E-16 | -0.025379 | 0.98011 |
| DG(34:2) | 0.011344 | 1.67E-16 | 0.025236 | 0.98023 |
| Hypoxanthine | 0.007401 | 5.13E-17 | 0.019455 | 0.98475 |
| Caproic acid | -0.0059435 | 1.82E-16 | -0.015874 | 0.98756 |
| TG(48:1) | 0.0060918 | -6.66E-17 | 0.010189 | 0.99202 |
| Uric acid | 0.0028368 | -2.84E-17 | 0.0061319 | 0.99519 |
| C14:2 | 0.0032539 | -9.30E-17 | 0.0058531 | 0.99541 |
