## Supplementary Table 8 for "Cell and Transcriptomic Diversity of Infrapatellar Fat Pad during Knee Osteoarthritis"

**Supplementary Table 8: RNA quality control for all IFP used for spatial sequencing.**

| Spatial Sample | RIN Number | DV200 Percentage |
|---|---|---|
| 1 | 9.1 | 97% |
| 2 | 9 | 96% |
| 3 | 8 | 96% |
| 4 | 7.9 | 96% |
| 5 | 9.3 | 97% |
| 6 | 9 | 98% |
| 7 | 6.5 | 95% |
| 8 | 7.8 | 88% |
| 9 | 8.9 | 97% |
| 10 | 9.5 | 99% |
| 11 | 9.8 | 97% |
| 12 | 8.8 | 90% |
